# Lanthipeptide Synthetases Participate the Biosynthesis of 2-Aminovinyl-Cysteine Motifs in Thioamitides

**DOI:** 10.1101/2020.08.21.260323

**Authors:** Jingxia Lu, Yuan Wu, Jiao Li, Yuqing Li, Yingying Zhang, Zengbing Bai, Jie Zheng, Jiapeng Zhu, Huan Wang

**Affiliations:** State Key Laboratory of Coordination Chemistry, Chemistry and Biomedicine Innovation Center of Nanjing University, Jiangsu Key Laboratory of Advanced Organic Materials, School of Chemistry and Chemical Engineering, Nanjing University, Nanjing 210093, China; School of Medicine and Life Sciences, State Key Laboratory Cultivation Base for TCM Quality and Efficacy, Jiangsu Key Laboratory for Pharmacology and Safety Evaluation of Chinese Materia Medica, Nanjing University of Chinese Medicine, Nanjing 210023, China

## Abstract

Thioamitides are a group of ribosomally synthesized and post-translational modified peptides with potent antiproliferative and pro-apoptotic activities. Their biosynthesis remains largely unknown, especially for the characteristic C-terminal 2-<u>a</u>mino<u>vi</u>nyl-<u>Cys</u>teine (AviCys) motifs. Herein, we report the discovery that homologs of class III lanthipeptide synthetases (LanKC_t_s)encoded *outside* putative thioamitide biosynthetic gene clusters (BGCs) fully dehydrate the precursor peptides. Remarkably, LanKC_t_ enzymes bind tightly to cysteine decarboxylases encoded *inside* thioamitide BGCs, and the resulting complex complete the macrocyclization of AviCys rings. Furthermore, LanKC_t_ enzymes are present in the genomes of many thioamitide-producing strains and are functional when in complex with cysteine decarboxylases to produce AviCys macrocycles. Thus, our study reveals the participation of lanthipeptide synthetases as a general strategy for dehydration and AviCys formation during thioamitides biosynthesis and thus paves the way for the bioengineering of this class of bioactive natural products.

## Introduction

Ribosomally synthesized and post-translationally modified peptides (*RiPPs*) have emerged as a major family of natural products with diverse bioactivities.^1^ Thioamitides^2^ are a subgroup of *RiPPs* that contain thioamides in place of amides in peptide backbones as their class-defining feature.^3–7^ As the first example of this group of compounds, apoptosis inducer thioviridamide contains a number of unnatural amino acids, including backbone thioamides, a *β*-hydroxy-*N1,N3*-dimethylhistidinium (hdmHis) residue, a N-terminal δ-hydroxy-δ-methyl-4-oxopentanoyl group and a C-terminal 2-aminovinyl-cysteine (AviCys) motif (Fig. 1A).^8–10^ Posttranslational modification enzymes are identified in the *tva* gene cluster (Fig. S1A), including TvaH/TvaI as a pair of homologs to YcaO/TfuA proteins for the thioamidation of peptide backbone and TvaG as a putative methyltransferase for the double methylation of His(12) residue.^11^ The N-terminal 2-hydroxy-2-methyl-4-oxopentanoyl group is the product of acetone addition to a pyruvate motif derived from the hydrolysis of a dehydroalanine (Dha) residue during purification.^6, 12^ The C-terminal 2-aminovinyl-cysteine (AviCys) motif is another structural feature of thioamitides and exists in a variety of *RiPPs*, including lanthipeptides (e.g., epidermin),^13–15^ lipolanthines (e.g., microvionin)^16^ and linaridins (e.g., cypemycin)^17^ (Fig. 1A). All AviCys-containing compounds reported to date exhibit potent bioactivities, implying the biological importance of this cyclic scaffold. Current proposal for the formation of AviCys and 2-aminovinyl-3-methyl-Cysteine (AviMeCys) motifs in thioamitides involves three consecutive enzymatic modifications. First, Ser/Thr residues in the precursor peptide are converted into Dha/Dhb residues by a dehydratase. A cysteine decarboxylase then oxidatively decarboxylated the C-terminal cysteine to generate a thioenol motif. In the final step, a putative cyclase catalyzes the Michael-type addition between the thioenol group and a Dha/Dhb residue to yield an AviCys/AviMeCys motif (Fig. 1B).^17, 18^ Although the function of cysteine decarboxylases has been well characterized in several cases,^19–21^ putative dehydratases and cyclases are not yet identified in the biosynthetic gene clusters (BGCs) of thio-amitides, leaving the AviCys biosynthesis unresolved. Herein, we report the discovery that homologs of class III lanthipeptide synthetases (LanKC_t_s) locating *outside* the putative thioamitide BGCs function as dehydratases to dehydrate Ser/Thr residues in thioamitide precursor peptides. Remarkably, these LanKC_t_ enzymes bind tightly to cysteine decarboxylases (LanD_t_, LanD-like enzymes from thioamitide biosynthesis) encoded *inside* thioamitide BGCs and cooperatively catalyze the formation of AviCys/AviMeCys motifs at the C-terminus of peptide substrates. The combination of LanKC_t_-LanD_t_ displays remarkable tolerance toward peptide substrates by generating AviCys/AviMeCys rings of various sizes and sequences. This study reveals an unprecedented example that lanthipeptide synthetases encoded *outside* thioamitide BGCs participate the macrocyclization of AviCys/AviMeCys rings through direct association with corresponding LanD_t_ enzymes.

**Figure 1.**
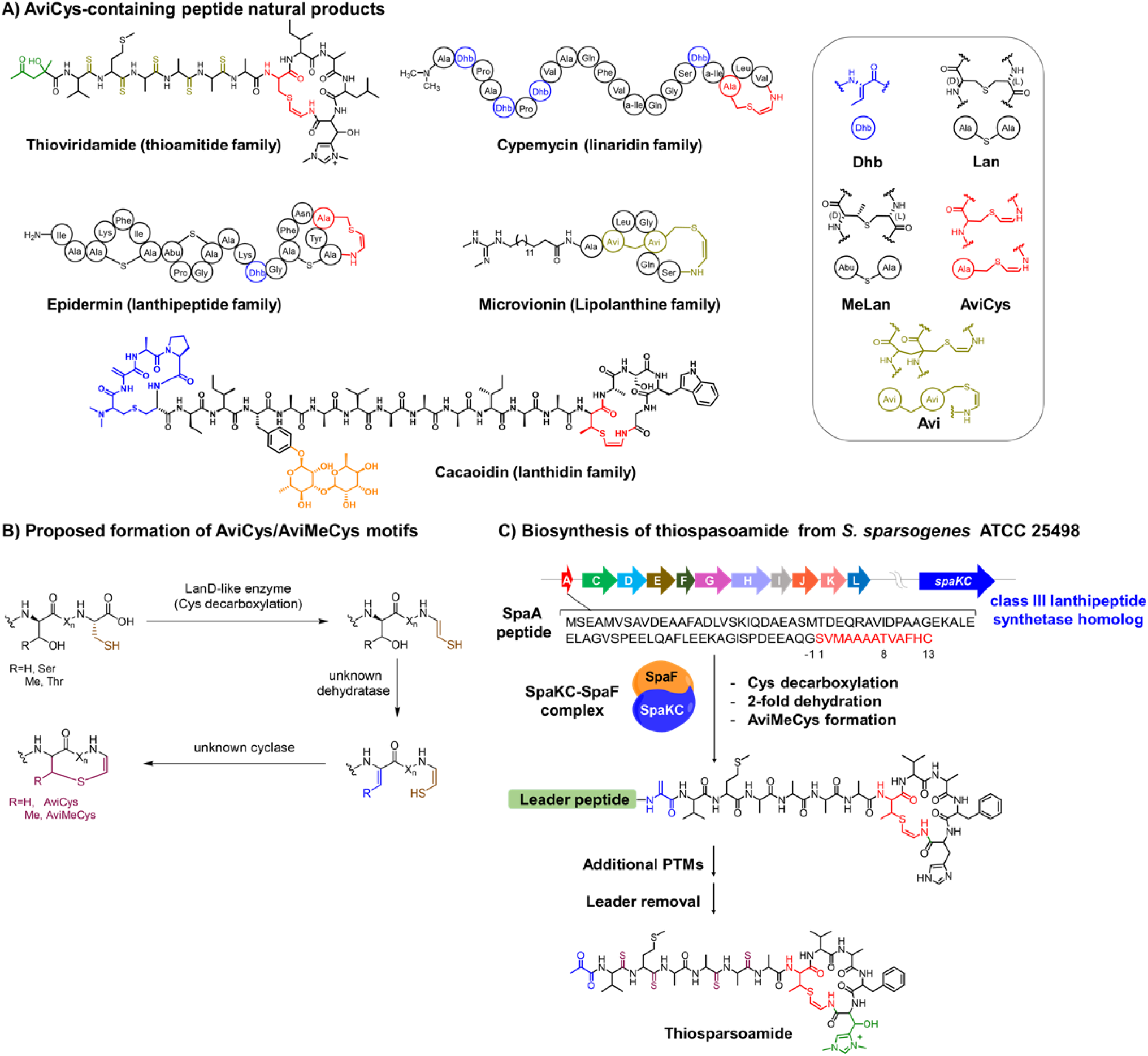
Structures and biosynthesis of AviCys/AviMeCys-containing compounds. A). AviCys-containing peptide natural products. B). Proposed biosynthesis of AviCys/AviMeCys motifs. C). Biosynthesis of thiosparsoamide, the putative product of the *spa* gene cluster. SpaA core peptide is marked in red.

## RESULTS

### Discovery of a new thioamitide natural product thiosparsoamide

To investigate the biosynthesis of thioamitides, we focused on a *spa* gene cluster from the strain of *S. sparsogenes* ATCC 25498, which shares high homology to the thioviridamide BGC (*tva* gene cluster) (Fig. 1C, Fig. S1-S2). We speculated that the final product of *spa* gene cluster, named thio-sparsoamide, contains multiple thioamide motifs, an hdmHis residue, a C-terminal AviMeCys macrocycle and an N-terminal Dha residue, which could be hydrolyzed to a pyruvate motif in culture medium (Fig. 1C, Fig. S1). Indeed, fermentation of *S. sparsogenes* strain led to the isolation of a peptide product with a mass of 1305.493 Da (Fig. S3), which matches with predicted mass of thiosparsoamide. Tandem MS analysis further confirmed the presence of expected unnatural amino acids in this peptide product (Fig. S3). Together, these results confirm the function of *spa* gene cluster in producing a new thioamitide compound thiosparsoamide.

### Class III lanthipeptide synthetase SpaKC and decarboxylase SpaF catalyze the formation of AviMeCys motif

To understand the biosynthesis of thiosparsoamide, SpaF, a putative cysteine decarboxylase encoded in the *spa* gene cluster, was co-expressed with precursor peptide SpaA in *E. coli*. SpaF oxidatively decarboxylates SpaA by producing an unstable thioenol product **SpaA-(a)** with a mass loss of 46 Da (Fig. 2A, Fig. S4). **SpaA-(b)** and **SpaA-(c)** with mass losses of 62 Da and 104 Da, respectively, were detected as the hydrolysis products of **SpaA-(a)**(Fig. 2A, Fig. S4-S5). Degradation of thioenol products from peptide decarboxylation is spontaneous in buffer conditions and precedent in the cases of several LanD-like enzymes, including TvaF from thioviridamide biosynthesis, EpiD from epidermin biosynthesis and MicD from microvionin biosynthesis.^16, 22^ To facilitate the removal of leader peptide, SpaA_G-1K_ peptide was used as peptide substrate for further studies (Fig. S6-S7). To provide mechanistic insight into the function of SpaF, we resolved its crystal structure at 2.15 Å resolution. In solution, SpaF forms a complex with a molecular weight ~ 300 kDa with FMN bound as a cofactor (Fig. S8). Consistent with this observation, crystal-packing shows that SpaF assembles into a homododecamer, in which trimers locate at the vertices of a tetrahedron (Fig. 2B). Each trimer unit binds with one FMN molecule in a cavity at the interface of two monomers (Fig. S9). The overall structure of SpaF is highly similar to those of TvaF, CypD and EpiD despite their low sequence similarity, suggesting the convergent evolution of cysteine decarboxylases of this class.^23^ Together, our results showed that SpaF is a FMN-dependent enzyme that catalyzes the cysteine decarboxylation of the precursor peptide SpaA.

**Figure 2.**
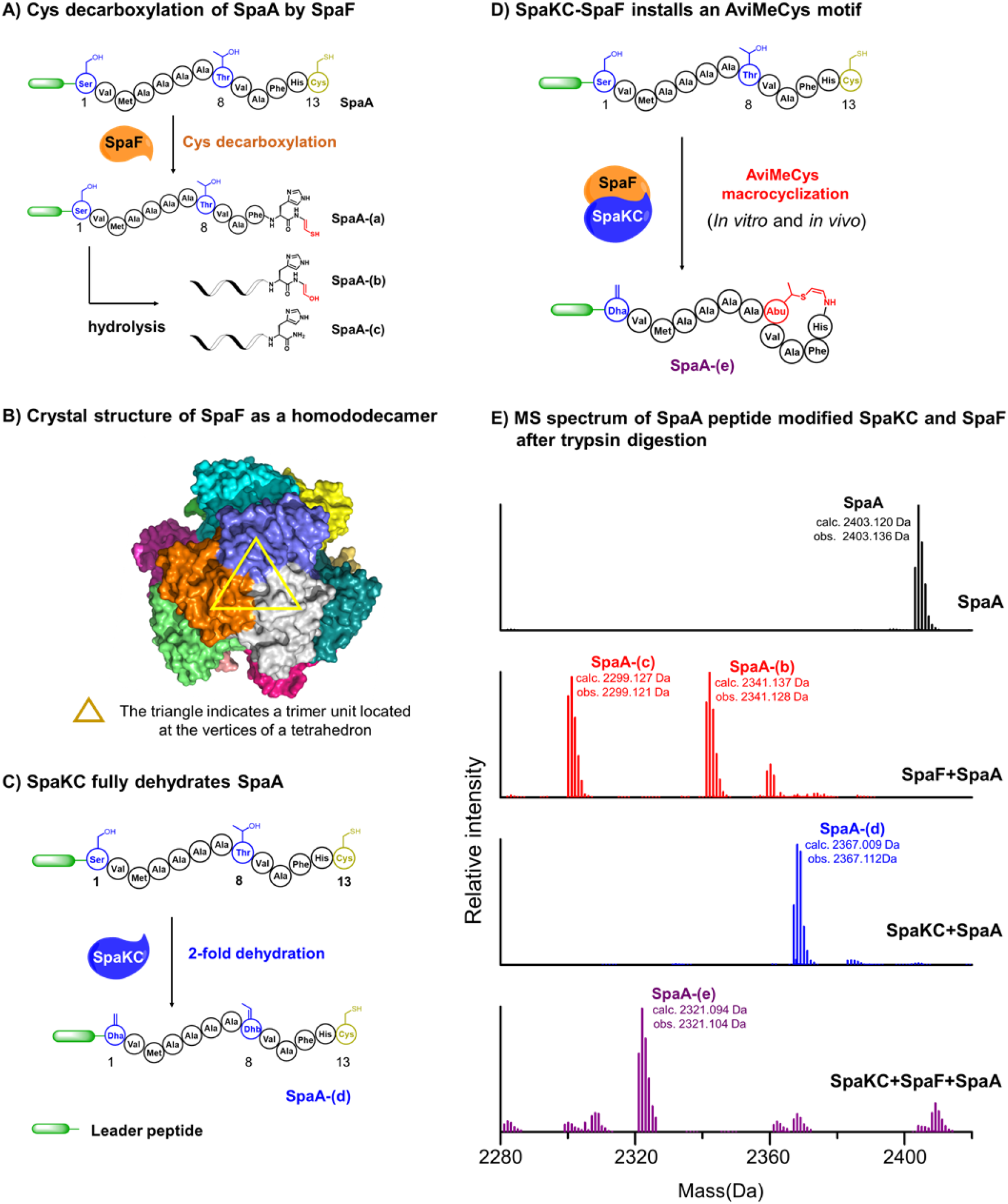
Enzymatic modification of SpaA peptide by SpaF and SpaKC *via* a co-expression system in *E. coli*. (A). SpaF converts SpaA into decarboxylation product **SpaA-(a)**, which spontaneously hydrolyzes into **SpaA-(b)** and **SpaA-(c)** in buffer. (B) Crystal structure of SpaF as a homododecamer (PDB ID: 7CFU). (C) LC-MS analysis of the SpaKC-modified SpaA_G-1K_ peptide **SpaA-(d)** with 2-fold dehydration. **SpaA**_**G-1K**_**-(d)**_(−11)−13)_ is the core peptide fragment of peptide **SpaA**_**G-1K**_−**(d)** after trypsin digestion. (D) SpaKC-SpaF combination converts SpaA_G-1K_ peptide into **SpaA-(e)** with a Dha residue and an AviCys motif. (E) MS spectrum of modified SpaA peptides after trypsin digestion.

Next, we focused on elucidating the identity of dehydratase and cyclase for the dehydration and AviMeCys formation in thiosparsoamide biosynthesis (Fig. 1C). We noticed that leader peptides of thioamitide precursors contain highly conserved PE<u>E(L/A)Q</u> motifs, which shares similarity to the L<u>ELQ</u>G motif conserved in leader peptides of class III lanthipeptide precursors for enzymatic recognition (Fig. S2 and S10).^24, 25^ In addition, class III lanthipeptide synthetase (LanKC) AplKC from NAI-112 biosynthesis is able to phosphorylate SpaA peptide *in vitro* (Fig. S11), indicating that SpaA can be recognized by a LanKC enzyme.^26, 27^ Together, these data raise the possibility that SpaA peptide could be modified by a LanKC enzyme during thiosparsoamide biosynthesis. By scanning the genome of *S. sparsogenes* ATCC 25498, we found a gene, named *spaKC*, which encodes a LanKC homolog and is located far outside the *spa* gene cluster without association with any putative biosynthetic gene clusters of secondary metabolites (Fig. 1C, Table S1). Co-expression of *spaKC* and *spaA_G-1K_* genes produced **SpaA**_**G-1K**_-**(d)** with 2-fold dehydration at Ser(1) and Thr(8) residues as the only product, showing that SpaKC efficiently recognizes SpaA_G-1K_ as a substrate for dehydration (Fig. 2B, Fig. S12). No ring structure was formed in **SpaA**_**G-1K**_-**(d)**, suggesting that SpaKC is not capable of catalyzing the formation of a Lan/MeLan macrocycle between Cys(13) and Dha(1)/Dhb(8) or **SpaA**_**G-1K**_-**(d)** is not a substrate for macrocyclization. Next, SpaA_G-1K_ peptide was co-expressed with SpaKC and SpaF enzymes in *E. coli*, yielding a peptide product **SpaA**_**G-1K**_-**(e)** with a mass loss of 82 Da compared with SpaA peptide, matching two-fold dehydration (−36 Da) and one oxidative decarboxylation (−46 Da) (Fig. 2C). Tandem MS analysis showed that **SpaA**_**G-1K**_-**(e)** contains a Dha(1) and a C-terminal macrocycle between the 8^th^ and 13^th^ residues (Fig. 2D, Fig. S13). In addition, iodoacetamide (IAA) treatment of **SpaA**_**G-1K**_-**(e)** resulted in no mass change, supporting that the thiol group of Cys(13) was consumed during the ring formation (Fig. S14).

To further verify the identity of the C-terminal macrocycle, **SpaA**_**G-1K**_-**(e)** peptide was reduced and desulfurized by NiCl_2_ and NaBH_4_ treatment in H_2_O following an established protocol (Fig. 3A, Fig. S15).^28^ Tandem MS analysis of the desulfurization product **SpaA**_**G-1K**_-**(f)** revealed the presence of an Ala(1) residue, two homoalanine (hAla) residues at the 3^rd^ and 8^th^ positions and an ethylamine-derived His residue at the C-terminus, supporting the structural assignment of **SpaA**_**G-1K**_-**(e)** with a Dha(1) residue and a C-terminal AviMeCys motif. The hAla(3) residue of **SpaA**_**G-1K**_-**(f)** is derived from the Met(3) of **SpaA**_**G-1K**_-**(e)** via sidechain cleavage during desulfurization. When NaBD_4_ and D_2_O were employed to linearize **SpaA**_**G-1K**_-**(e)**, Dha(1) was reduced to an Ala residue with double deuteration, whereas Met(8) was converted into a hAla residue with one deuterium substitution in product **SpaA**_**(G-1K)**_-**(g)** (Fig. 3B, Fig. S16). Importantly, single deuterium and triple deuterium substitutions were observed in the hAla(8) residue and the C-terminal 12^th^ residue in **SpaA**_**(G-1K)**_-**(g)**, respectively, further supporting the installation of a C-terminal AviMeCys motif in SpaA peptide. Collectively, our results showed that the combination of SpaF-SpaKC enzymes is capable of catalyzing the AviMeCys formation in precursor peptide SpaA.

**Figure 3.**
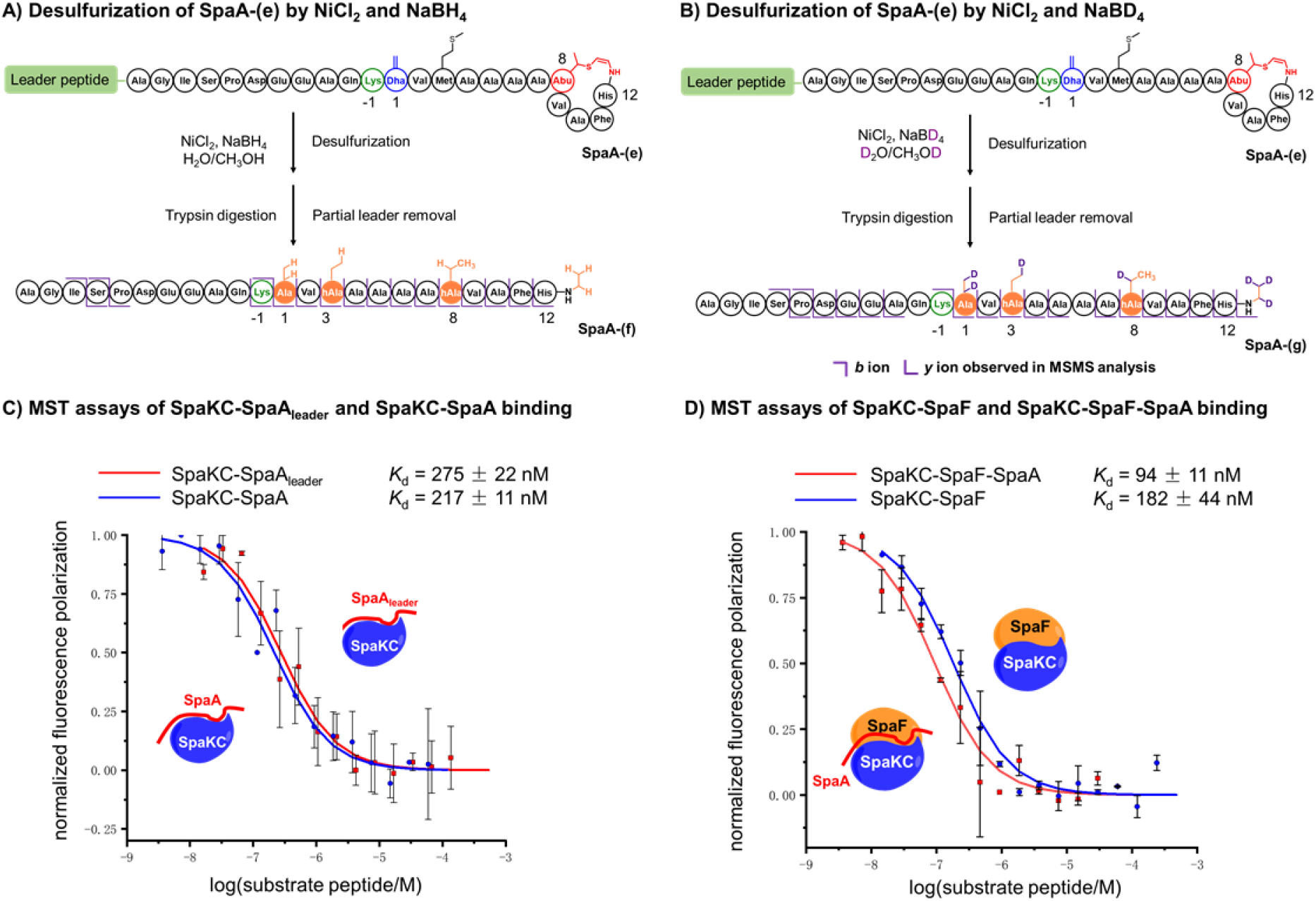
Characterization of AviMeCys formation and binding affinities between SpaKC, SpaF and SpaA. (A-B) Characterization of AviMeCys formation in **SpaA-(e)** peptide by reductive desulferization and MS/MS analysis of the resulting linearized product. *b* ions and *y* ions resulted from fragmentation are shown in purple. (C-D) MST analysis of the binding affinity between SpaKC and SpaA, SpaA leader peptide, SpaF. Error bars, s.e.m. of three independent measurements. Experimental details are described in the supporting information.

### SpaKC and SpaF function as a complex

The formation of AviMeCys motif includes dehydration of Ser/Thr residues, cysteine decarboxylation and AviMeCys macrocyclization. To probe the order of modifications, we examined the *in vitro* activity of SpaKC and SpaF toward various peptide substrates. SpaKC is fully functional as a dehydratase *in vitro* with ATP as a cofactor to dehydrate SpaA and decarboxylated SpaA peptides (Fig. S17-S18). Similarly, SpaF is highly efficient in decarboxylating SpaA peptide, but less efficient with dehydrated SpaA peptide **SpaA**_**(G-1K)-**_**(d)** (Fig. S19). No cyclization product was detected in the abovementioned assays when SpaKC or SpaF functions alone. Only when SpaA, SpaKC and SpaF were incubated together, cyclized product **SpaA**_**G-1K**_-**(e)** was successfully produced *in vitro* (Fig. 2D, Fig. S20), indicating that the combination of SpaKC and SpaF enzymes was strictly required for AviMeCys installation. In order to understand the cooperative nature of SpaKC-SpaF enzymes, the interactions between SpaKC, SpaF and SpaA peptides were measured by microscale thermophoresis (MST) assays (Fig. 3C and 3D). Results showed that SpaKC binds to SpaA leader peptide and SpaA peptide with similar *K*_d_ values of (275 ± 23) nM and (217 ± 12) nM, indicating that SpaKC functions in a leader dependent manner. Remarkably, SpaKC and SpaF bind to each other tightly with a *K*_d_ of (182 ± 44) nM, showing that SpaKC and SpaF function as a complex in solution. Remarkably, the binding between SpaKC-SpaF complex and SpaA peptide exhibited a *K*_d_ of (94 ± 11) nM, which is 2-fold lower than that of SpaKC-SpaA binding, indicating that SpaA peptide enhances the stability of the enzyme complex during modification. Together, our results show that SpaKC and SpaF function as a complex and process peptide substrate in a leader dependent manner to achieve AviMeCys macrocyclization (Fig. 4).

**Figure 4.**
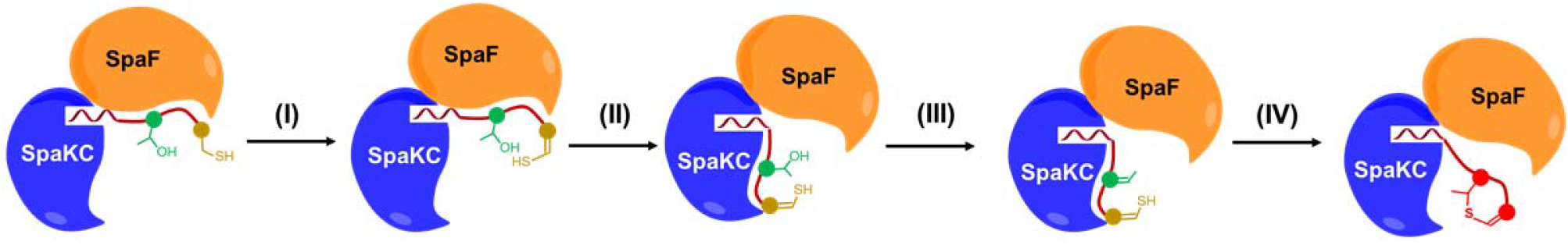
Current model of AviMeCys formation catalyzed by SpaKC-SpaF complex by the following steps: (I). First, SpaA peptide binds to the SpaKC-SpaF complex via leader-SpaKC association; (II). As the first modification, SpaF oxidatively decarboxylates the C-terminal Cys residue in SpaA; (III). The decarboxylated core peptide is relocated to SpaKC for dehydration; (IV). The dehydrated and decarboxylated SpaA peptide is cyclized to form an AviMeCys motif.

Next, we examined the versatility of SpaKC-SpaF complex in generating AviCys/AviMeCys motifs in various SpaA mutants. Mutation of AviMeCys precursor residues Thr(8) and Cys(13) to Cys and Ala, respectively, abolished the AviMeCys formation while dehydration of Ser/Thr occurs efficiently (Fig. 5A, entry 1-2, Fig. S21-22). Ala mutation at any other position of SpaA, including residues in the ring region, has little impact on the macrocyclization of AviMeCys motif in the corresponding SpaA peptide (Fig. 5A, entry 3-5, Fig. S23-25). Furthermore, SpaKC-SpaF combination exhibited remarkable flexibility when the size of the AviMeCys ring was altered. Deletion or insertion of residues between Thr(8) and Cys(13) did not significantly affect macrocyclization, and AviMeCys rings of sizes varying from four to eight amino acids were efficiently generated (Fig. 5B, entry 6-11, Fig. S26-31). Similarly, SpaA peptides carrying mutations or deletion of non-ring-forming residues in the core peptide were well tolerated by SpaKC-SpaF combination by yielding corresponding cyclized products (Fig. 5A, entry 12-14, Fig. S32-34). In addition, SpaA_T8S_ peptide were efficiently cyclized by forming an AviCys motif, further demonstrating the substrate flexibility of the SpaKC-SpaF combination (Fig. 5B, entry 15, Fig. S35). Overall, SpaKC-SpaF combination displayed remarkable substrate tolerance toward SpaA peptides for AviCys/AviMeCys installation.

**Figure 5.**
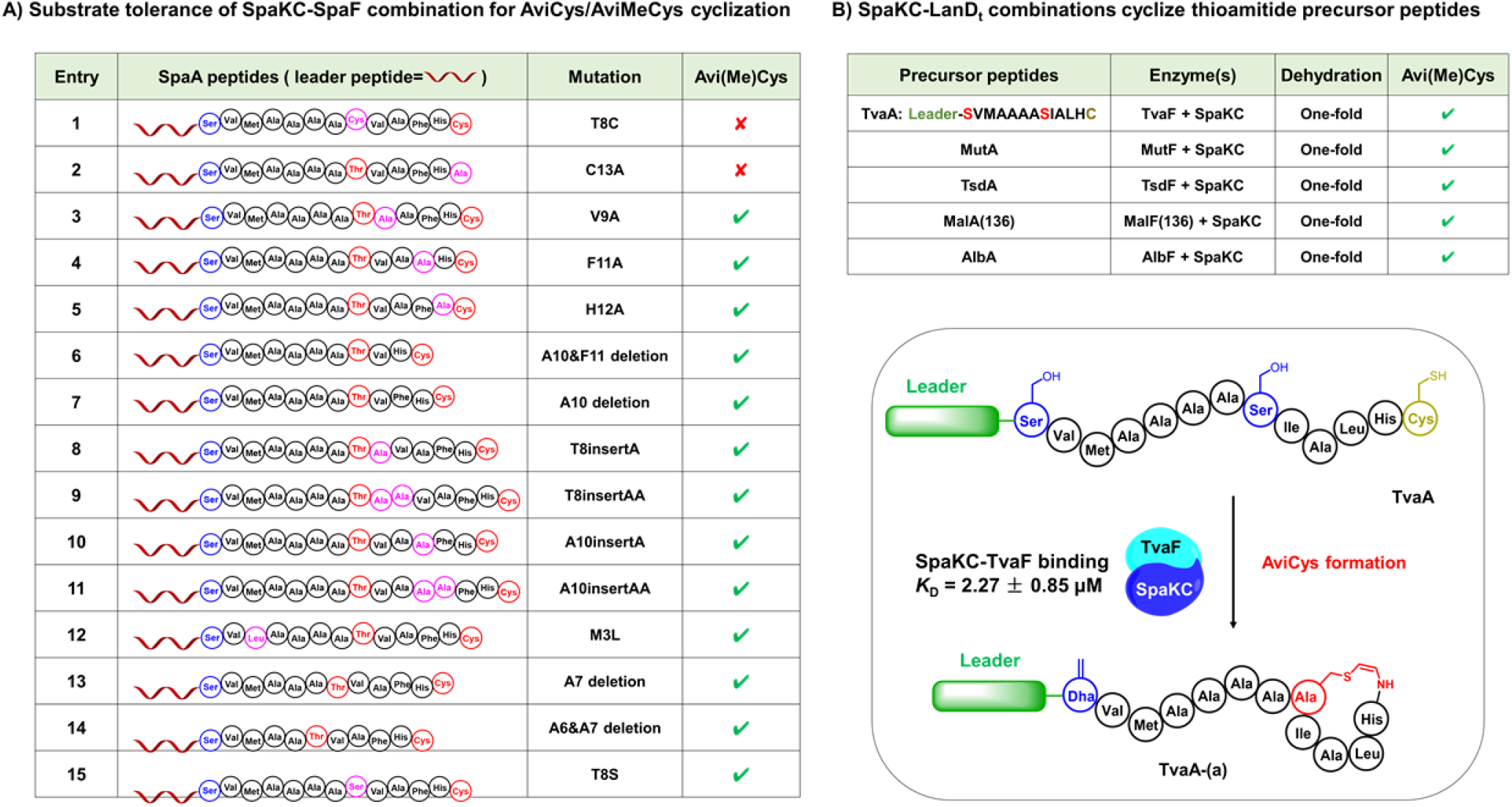
SpaKC and SpaF function as a complex and allows generation of AviCys/AviMeCys rings of various sizes and sequences. SpaKC installs Dha residues and AviCys/AviMeCys motifs in precursor peptides of thioviridamide-like natural products. (A). Precursor peptides of thioviridamide-like natural products are modified by SpaKC or the enzyme pair of SpaKC-(cysteine decarboxylase). (B). SpaKC and TavF form a complex and modify TvaA peptide by installing a Dha residue and a C-terminal AviCys motif.

### Utilization of LanKC enzymes for dehydration and AviCys cyclization is a general strategy for thioamitide biosynthesis

The thioamitide precursor peptides share high sequence similarity (Fig. S2A), raising the possibility to utilize SpaKC as an enzymatic tool to dehydrate these peptides. Five precursor peptides, including TvaA from *S. olivoridis* NA05001, TsdA from *S. sp* NRRL S-87, MutA from *S. mutomycini* NRRL B65393, AlbA from *A. alba* DSM 44262, and MalA(136) from *S. malaysiense* MUSC 136 (Fig. S2A), were co-expressed with SpaKC in *E. coli*. As expected, SpaKC efficiently modified all five precursor peptides with two-fold dehydration (Fig. S36-40), demonstrating the substrate tolerance of SpaKC as a dehydratase.

Next, we examined whether SpaKC could function with LanD_t_ decarboxylases from thioamitide BGCs by installing AviCys/AviMeCys motifs in corresponding precursor peptides. TvaF from thioviridamide biosynthesis was selected as a model and shown to bind to SpaKC with a *K*_d_ of (2.27 ± 0.85) μM (Fig. S41), indicating that SpaKC and TvaF can function as a complex. When co-expressed with TvaA peptide in *E. coli*, SpaKC-TvaF combination efficiently modified the peptide substrate by installing one Dha at the 1^st^ position and one AviCys motif at the C-terminus (Fig. 5B, Fig. S42). Similarly, when SpaKC was co-expressed with MutF/MutA, TsdF/TsdA, MalF(136)/MalA(136) and AlbF/AlbA pairs, AviCys motifs were correctly installed at the C-terminus in all peptide substrates (Fig. 5A, Fig. S43-S46). Therefore, SpaKC serves as a general dehydratase and functions cooperatively with LanD_t_ enzymes by forming protein complexes to install AviCys/AviMeCys motifs in thioamitide precursor peptides.

The capability of SpaKC to function with various LanD_t_ enzymes for AviCys/AviMeCys formation implies that the utilization of class III lanthipeptide synthetase (LanKC) homologs might be a general strategy for thioamitide biosynthesis. To date, 18 bacterial strains have been identified to contain putative thioamitide BGCs in their genomes. By scanning the genomes of these bacteria (17 out of 18 are fully sequenced), we identified 12 genes encoding LanKC homologs from 10 strains, which are all located far *outside* the putative thioamitide BGCs (Table 1, Table S1). Both sequence Similarity Networks (SSNs) analysis and maximum-likehood phylogeny analysis show that these LanKC homologs (named LanKC_t_, <u>LanKC</u> homologs in thioamitide biosynthesis), including SpaKC, are closely related to known class III lanthipeptide synthetases phylogenetically (Fig. S47-S48). To examine the participation of LanKC_t_ enzymes in thioamitide biosynthesis, we selected MalA(136), the precursor peptide from the thioamitide BGC in *S. malaysiense* MUSC 136 strain, as well as the LanKC_t_ enzyme MalKC(136)-1 discovered from its genome, as a model (Fig. S49). Results showed that MalKC(136)-1 efficiently dehydrated MalA(136) peptide by up to 2-fold during co-expression in *E. coli* (Fig. S50), confirming the role of MalKC(136)-1 as a dehydratse to modify the corresponding thioamitide precursor peptide. Together, our results indicate that the participation of a LanKC_t_ enzymes encoded outside corresponding thioamitide BGCs is a general strategy employed during thioamitide biosynthesis.

**Table 1.** LanKC_t_ genes discovered in the genomes of thioamitide-producing strains. All genes are located outside corresponding thioamitide BGCs.

| Strain | LanKC<sub>t</sub> | Accession number |
| --- | --- | --- |
| *S. sparsogenes* ATCC 25498 | SpaKC | WP_065963996.1 |
| *S. sp.* MUSC 14 | SmuKC(14) | WP_071371419.1 |
| *S. sp.* MUSC 125 | SmuKC(125) | WP_039650135.1 |
| *S. malaysiense* MUSC 136 | MalKC(136)-1 | OIK27458.1 |
|  | MalKC(136)-2 | OIK27459.1 |
| *S. malaysiense* | MalKC | WP_071381943.1 |
| *S. sp.* CNB091 | ScnKC-1 | WP_026290248.1 |
|  | ScnKC-2 | WP_018959331.1 |
| *S. sp.* NRRL S-4 | SnrKC(S-4) | WP_053928640.1 |
| *S. mutomycini* NRRL B-65393 | MutKC | WP_065848356.1 |
| *Amycolatopsis alba* DSM 44262 | AmyKC | WP_020633044.1 |
| *Nocardiopsis potens* DSM 45234 | NocKC | WP_017595403.1 |

## Discussion

AviCys/AviMeCys motifs exist in several classes of *RiPPs* and are essential for their bioactivities. Elucidation of the multiple enzymatic transformations during AviCys/AviMeCys formation is a prerequist for the bioengineeing of these bioactive compounds. Our investigation reveals that LanKC_t_ enzymes encoded *outside* putative thioamitide BGCs fully dehydrate the precursor peptides. LanKC_t_ and LanD_t_ enzymes encoded inside thioamitide BGCs may form complexes and construct the AviCys/AviMeCys rings cooperatively. Regarding the order of modifications during AviCys formation, our results suggest that cysteine decarboxylation occurs before the dehydration of precursor peptides. The formation of a LabKC_t_-LanD_t_ complex is therefore benefical for the unstable thioenol intermediate to quickly relocate from LanD_t_ to the LanKC_t_ enzyme for dehydration and subsequent cyclization by minimizing possibility of hydrolysis. We further show that LanKC_t_ enzymes are widespread in the genomes of most thioamitide-producing strains and are all functional to dehydrate corresponding thioamitide precursor peptides, suggesting LanKC_t_-LanD_t_ combination is a general strategy for AviCys/AviMeCys formation. Together, our study reveals a very rare system in the biosynthesis of *RiPPs* that a modification enzyme encoded *outside* the BGC function cooperatively with an enzyme encoded *inside* the BGC through specific association to construct a key structural element in the final product. The detailed catalytic mechanism for AviCys/AviMeCys cyclization in LanKC_t_-LanD_t_ complex requires further investigation.

It is highly possible that proteins of unknown functions encoded in the putative thioamitide BGCs possess dehydratase function for AviCys formation, since 8 out of 18 thioamitide-producing strains do not contain LanKC_t_ enzymes in their genomes. In addition, successful heterologous production of thioviridamide and derivatives by the putative *tva* gene cluster has been reported in *S. lividans* and *S. avermitilis*, suggesting the *tva* gene cluster is fully capable of producing thioviridamides.^10, 29–31^ However, it is noteworthy that LanKC_t_ enzymes are encoded in the genomes of both *S. lividans* and *S. avermitilis* strains, and their functions remain unknown (Table S2). Therefore, we hypothesize that LanKC_t_ enzymes may not be essential for thioamitide biosynthesis, but serve as complementary dehydratases to support the production of thioamitides when neccesasry.

Recently, coorpoerative function of MicKC-MicD enzyme pair is reported in the formation of avionin motifs during microvionin biosynthesis, in which both enzymes are encoded *in* the BGCs (Fig. 1A, Fig. S51).^29, 32^ MicKC and MicD function in a mutual regulotory manner, where the activity of MicKC is highly dependent on the presence of MicD. In contrast, SpakC and SpaF are fully functional saparately for dehydration and decarboxylation, respectively, desipte the requirement of complex formation for AviCys/AviMeCys macrocyclization. The biosynthesis of lipolanthines and thioamitides suggest that the cooperative function of LanKC- and LanD-like enzymes for AviCys-like macrocyclization might be a general biosynthetic strategy, which requires further investigation to understand the detailed enzymatic mechanisms.

## ASSOCIATED CONTENT

### Supporting Information

Detailed compound characterization and MS spectra are provided. Supporting Information is available free of charge via the Internet at http://pubs.acs.org.

## AUTHOR INFORMATION

### Author Contributions

The manuscript was written through contributions of all authors. All authors have given approval to the final version of the manuscript.

### Notes

The authors declare no competing financial interests.

## ACKNOWLEDGMENT

This work is supported by NSF of China (Grant 21778030 and 21861142005 to H.W.) and the Fundamental Research Funds for the Central Universities (Grant 14380138 and 14380131 to H.W.).

## Methods

### General methods

Primers, genes and peptide were synthesized by Genscript Biotech (Nanjing, China). Restriction endonucleases and T4 DNA ligase were purchased from New England Biolabs (Ipswich, MA, USA). Phanta^®^ Max Master Mix were purchased from Vazyme Biotech Co., Ltd (Nanjing, China). Medium components for bacterial cultures were purchased from Thermo Fisher (Waltham, MA, USA). Chemicals were purchased from Aladdin Reagent (Shanghai, China) and Sigma-Aldrich (Schnelldorf, Germany). Endoprotease trypsin was purchased from Roche Biosciences (Basel, Switzerland). E. coli DH5α was used as a host for cloning and plasmid propagation, and E. coli BL21 (DE3) was used as a host for expression of proteins and peptides. S. sparsogenes ATCC 25498 is purchased from (China General Microbiological Culture Collection Center, CGMCC).

All polymerase chain reactions (PCR) were carried out on a C1000 Touch™ thermal cycler (Bio-Rad). DNA sequencing was performed by the Genscript Biotech. Matrix-assisted laser desorption/ionization time-of-flight mass spectrometry (MALDI-TOF MS) was carried out on Bruker UltraFlextreme. Liquid chromatography electrospray ionization tandem mass spectrometry (LC/ESI-MS/MS) was carried out and processed using a Triple TOF 4600 System (AB SCIEX) equipped with a Prominence Ultra Fast Liquid Chromatography (UFLC) system (Shimadzu). UV-Vis spectrometry was conducted using Cary 300 (Agilent Technologies).

Conditions for all ESI-MS and MS/MS were set as follows: nebulizer gas: 55 psi; heater gas: 55 psi; curtain gas: 35 psi; drying temperature: 550 °C; ion spray voltage: 5500 V; declustering potential: 100 V; collision energy: 35 V (positive); collision energy spread: 10 V. The mass range and accumulation time are 400-4000 m/z, 250 ms for ESI-MS and 100-2000 m/z, 100 ms for MS/MS, respectively. Collision-induced dissociation (CID) was performed for fragmentation of the respective peptide ions. Calibration solutions purchased from AB SCIEX were used for instrument calibration, and high resolution was chosen in the ESI+ mode

### Production and analysis of thiosparsoamide by *S. sparsogenes* ATCC 25498

*S. sparsogenes* ATCC 25498 was spread on PS5 agar plates for sporulation and growth. Approximately 1 cm^2^ of the agar with sporulated *S. sparsogenes* ATCC 25498 was used to inoculate 100 mL of the fermentation medium. After incubation at 28°C and 220 rpm for 96 hours, the whole broth was extracted with methanol, lyophilized, and re-dissolved with methanol. The resulting sample was subjected to LC-HRMS analysis for thiosparsoamide detection. PS5 agars: 20 g of starch, 5 g of pharmamedia, and 20 g of agar per liter (pH 7.0).Fermentation medium: 25 g of glucose, 15 g of soybean meal, 2 g of dry yeast, and 4 g of CaCO3 per liter (pH7.0). LC-MS analysis of thiosparsoamide was carried out with liquid chromatography heated electrospray ionization tandem mass spectrometry (LC/HESI-MS/MS) and processed using a Thermo Scientific™ Q Exactive™ equipped with a Vanquish Duo UHPLC system (Thermo scienctific). UV-Vis spectrometry was conducted using Diode Array Detector FG. Conditions for HESI-MS and MS/MS were set as follows: sheath gas: 45 psi; auxiliary gas: 15 psi; spare gas: 2.0 psi; capillary temperature: 320 °C; spray voltage: 3500 V; normalized collision energy: 30% of the available 5V (positive); The mass range and accumulation time are 200-2000 m/z, 100 ms for HESI-MS and data dependent top 20 ions were select by quadrupole selected and HCD (higher collisional dissociation) was performed for fragmentation of the respective ion.

### Molecular cloning of *spaA, spaF, spaKC, tvaA, tvaF, mutA, mutF, tsdA, tsdF, tsdKC, malA, malF, albA and albF* genes

Plasmids containing target genes were synthesized by Genscript Biotech and PCR-amplified by 30 cycles of denaturing (95 °C for 30 s), annealing (65 °C for 30 s), and extending (72 °C, 1 min/kb) using high fidelity Phanta DNA Polymerase. Amplifications were confirmed by 2% agarose gel electrophoresis, and the PCR products were purified using an Omega Biotech Cycle Pure Kit. Target DNA fragments and selected vectors were digested in separate reactions with selected pair of restriction enzymes for 2 h at 37 °C. The digested products were purified by agarose gel electrophoresis, and the DNA fragments were extracted from the gel using an Omega Biotech Gel Extraction Kit. The resulting DNA products were ligated at 16 °C for 12 h in T4 DNA Ligase buffer with T4 DNA Ligase. *E. coli* DH5α cells were transformed with 2.5 μL of the ligation product by heat shock, and cells were plated on proper antibiotics LB-agar plates and grown for 16 h at 37 °C. For each plasmid transformant, a single colony was picked to inoculate 5 mL culture of LB medium with proper antibiotics. The cultures were grown at 37 °C for 12 h, and plasmids were isolated using an Omega Biotech Plasmid Mini Kit. The sequences of the resulting plasmid products were confirmed by DNA sequencing.

### Construction of plasmids for coexpression of (His_6_-SpaA)-SpaF, (His_6_-TvaA)-TvaF, (His_6_-MutA)-MutF, (His_6_-TsdA)-TsdF, (His_6_-MalA)-MalF and (His_6_-AlbA)-AlbF

Amplification of the two genes was performed using high fidelity Phanta^®^ DNA Polymerase following the protocol described above. Restriction digestions were performed with selected pair of restriction enzymes following standard protocol. The digested products of genes were purified by agarose gel electrophoresis and ligated with pRSFDuet-1 vector. The resulting DNA products were ligated at 16 °C for 12 h in 1X T4 DNA Ligase buffer with T4 DNA Ligase (0.7 U/ μL). *E. coli* DH5α cells were transformed with 2.5 μL of the ligation product by heat shock, and cells were plated on LB-kanamycin agar plates and grown for 16 h at 37 °C. Several colonies were picked and used to inoculate separate 5 mL cultures of LB-kanamycin medium. The cultures were grown at 37 °C for 12 h, and plasmids were isolated using a Omega Biotech Plasmid Mini Kit. The sequences of the resulting plasmid products were confirmed by DNA sequencing.

### Mutagenesis of SpaA, SpaF, SpaKC, TsdA and MalA(136) genes

Mutation of SpaA was carried out by 30 cycles of denaturing (95 °C for 30 s), annealing (65 °C for 30 s), and extending (72 °C, 1 min/kb) using high fidelity Phanta^®^ DNA Polymerase. Amplifications were confirmed by 1% agarose gel electrophoresis, and the PCR products were purified using an Omega Biotech Cycle Pure Kit. The target DNA fragment was digested in reaction containing 1X NEB buffer (New England Biolabs) with DpnI for 3 h at 37 °C. *E. coli* DH5α cells were transformed with 10 μL of the digested product by heat shock, and cells were plated on LB-kanamycin agar plates and grown for 16 h at 37 °C. Several colonies were picked and used to inoculate separate 5 mL cultures of LB-kanamycin medium. The cultures were grown at 37 °C for 12 h, and plasmids were isolated using an Omega Biotech Plasmid Mini Kit. The sequences of the resulting plasmid products were confirmed by DNA sequencing. The other mutations were carried out by the protocol described above.

### Overexpression and purification of SpaF-modified SpaA peptides

*E. coli* BL21(DE3) cells were transformed with pRS-FDuet-1-(His_6_-SpaA)-SpaF, pRSFDuet-1-(His_6_-SpaA_G-1K_)-SpaF and plated on a Luria broth (LB) agar plate containing 50 mg/L of kanamycin. A single colony was used to inoculate a 5 mL culture of LB supplemented with 50 mg/L of kanamycin. The culture was grown at 37 °C for 12 h and was used to inoculate 4 L of LB containing 50 mg/L of kanamycin. Cells were grown at 37 °C to OD_600_=0.6-0.8 before IPTG was added to a final concentration of 0.2 mM and the culture was incubated at 18 °C for additional 16 h. Cells were harvested by centrifugation at 12,000 ×g for 25 min at 4 °C. The resulting cell pellet was resuspended in 30 mL of start buffer (20 mM NaH_2_PO_4_, pH 7.5, 500 mM NaCl, 0.5 mM imidazole, 20% glycerol), and the suspension was sonicated on ice for 20 min to lyse the cells. Cell debris was removed by centrifugation at 23,700 ×g for 30 min at 4 °C. The supernatant was discarded, and the pellet containing peptide products was resuspended in 30 mL of start buffer. The sonication and centrifugation steps were repeated. Again, the supernatant was discarded, and the pellet was re-suspended in 30 mL of buffer 1 (6 M guanidine HCl, 20 mM NaH_2_PO_4_, pH 7.5, 500 mM NaCl, 0.5 mM imidazole). The sample was sonicated and insoluble material was removed by centrifugation at 23,700 ×g for 30 min at 4 °C, followed by filtration of the supernatant through a 0.45 μm filter. The filtered sample was applied to a 5 mL HisTrap HP (GE Healthcare Life Sciences) immobilized metal affinity chromatography (IMAC) column previously charged with NiSO_4_ and equilibrated in buffer 1. The column was washed with two column volumes of buffer 1, followed by two column volumes of buffer 2 (4 M guanidine HCl, 20 mM NaH_2_PO_4_, pH 7.5, 500 mM NaCl, 30 mM imidazole). The peptide was eluted with 1-2 column volumes of elution buffer (4 M guanidine HCl, 20 mM NaH_2_PO_4_, pH 7.5, 500 mM NaCl, 1 M imidazole). The fractions were desalted using a Sep-Pak^®^ C18 Cartridges and analyzed by MALDI-TOF MS and the organic solvents were removed by rotary evaporation, followed by lyophilization. The product was kept at −80 °C for long-term storage. Typical yields from 4 L culture were 1 mg for His_6_-SpaA peptides.

### Overexpression of SpaKC with thioamitide precursor peptides

*E. coli* BL21(DE3) cells were transformed with pRS-FDuet-1-(His_6_-thioamitide precursor peptide) and pACYCDuet-1-SpaKC and plated on a Luria broth (LB) agar plate containing 50 mg/L of kanamycin and 35 mg/L of chloramphenicol. A single colony was used to inoculate a 5 mL culture of LB supplemented with 50 mg/L of kanamycin and 35 mg/L of chloramphenicol at 37 °C for 12 h. The culture was used to inoculate 4 L of LB containing 50 mg/L of kanamycin and 35 mg/L of chloramphenicol. Cells were grown at 37 °C to OD_600_=0.6-0.8 before IPTG was added to a final concentration of 0.2 mM and the culture was incubated at 18 °C for another 16 h before harvesting. The purification procedure was described as above.

### Overexpression and purification of (SpaKC-SpaF)-modified SpaA peptides

*E. coli* BL21(DE3) cells were transformed with pRSFDuet-1-His_6_-SpaA-SpaF and pACYCDuet-1-SpaKC or pACYCDuet-1-SpaKC_mutants_ and plated on a Luria broth (LB) agar plate containing 50 mg/L of kanamycin and 35 mg/L of chloramphenicol. A single colony was used to inoculate a 5 mL culture of LB supplemented with 50 mg/L of kanamycin and 35 mg/L of chloramphenicol at 37 °C for 12 h. The culture was used to inoculate 4 L of LB containing 50 mg/L of kanamycin and 35 mg/L of chloramphenicol. Cells were grown at 37 °C to OD_600_~0.6-0.8 before IPTG was added to a final concentration of 0.2 mM and the culture was incubated at 18 °C for another 16 h before harvesting. The purification procedure was described as above.

### Overexpression and purification of (SpaKC-LanD_t_)-modified thioamitide precursor peptides

The coexpression of SpaKC, TvaF and TvaA peptide is used as a representative example. *E. coli* BL21(DE3) cells were transformed with pRS-FDuet-1-(His_6_-TvaA)-TvaF and pACYCDuet-1-SpaKC plated on a Luria broth (LB) agar plate containing 50 mg/L of kanamycin and 35 mg/L of chloramphenicol. A single colony was used to inoculate a 5 mL culture of LB supplemented with 50 mg/L of kanamycin and 35 mg/L of chloramphenicol at 37 °C for 12 h. The culture was used to inoculate 4 L of LB containing 50 mg/L of kanamycin and 35 mg/L of chloramphenicol. Cells were grown at 37 °C to OD_600_=0.6-0.8 before IPTG was added to a final concentration of 0.2 mM and the culture was incubated at 18 °C for another 16 h before harvesting. The purification procedure was described as above. The other coexpression were carried out by the protocol described above.

### Determination of the flavin cofactor bound to SpaF

To identify the type of flavin cofactor, an aliquot of 200 μL of a 1.2 mM solution of His_6_-SpaF was denatured at 100 °C for 10 min. Precipitated protein was then removed by centrifugation (10,000 ×g, 10 min, 25 °C) and the released flavin was purified by Sep-Pak^®^ C18 Cartridges and analyzed by LC-ESI-MS.

### Modification of SpaA peptides with IAA

To determine the presence of free cysteine, SpaA peptides was digested by trypsin and then incubated with 20 mM IAA at 37 °C for 3h in 50 μL of reaction buffer (50 mM Tris, pH 8.6, 1 mM TCEP) in the dark, after which the reaction was quenched by addition of 2μL of 1 M DTT to eliminate excess IAA and prevent further side reactions. The reactions were further analyzed by MALDI-TOF.

### Reductive desulfurization of modified SpaA peptides

(SpaKC-SpaF)-modified SpaA_G-1K_ (1.0 mg) was suspended in 4.0 mL of CH_3_OH/H_2_O (1:1), to which 20 mg of NiCl_2_ and 20 mg of NaBH_4_ were added. This mixture was stirred under 1 atm of H_2_ at room temperature. After 8.5 h, the mixture was taken off the hydrogenation apparatus, and then centrifuged, followed by the removal of the supernatant. Mixed solvent of CH_3_OH/H_2_O (ration=1:1, 2.0 mL) was added to the black nickel boride pellets, and the suspension was sonicated for 15 minutes. The suspension was centrifuged, and the supernatants were collected. The organic solvent was removed in vacuo, and the desulfurized peptide was treated with trypsin. Digestion conditions: 20 mM Tris-HCl, pH = 8.0, 1 μM trypsin, 100 μM peptide product at 37 °C for 6 h.

### Measurement of protein-protein and protein-peptide interactions by MST assays

As a representative example, the binding affinity between SpaKC and SpaA_leader_ was measured using Monolith NT.115 Pico (Nanotemper Technologies). SpaKC was fluorescently labelled by the labeling kit (Monolith Protein Labeling Kit RED-NHS 2nd Generation (Amine Reactive), MO-L011). A volume of 90 μL SpaKC sample (10 μM) in the labelling buffer (130 mM NaHCO_3_, 50 mM NaCl, pH 8.2) was mixed with 10 μL dye (Dye RED-NHS 2nd Generation (Nanotemper MO-L011)) solution (300 μM) for 30 min at room temperature in the dark. Next, the SpaKC sample was loaded to column B (Nanotemper MO-L011) and eluted with 450 μL of assay buffer (50mM Na_2_CO_3_/NaHCO_3_, 150mM NaCl, 10mM MgCl_2_, 0.07% tween 20, pH 9.0). The labelled SpaKC sample was diluted until pretest fluorescence excitation range from 16000-20000 for the most optimized response (~1 nM). To perform the MST assay, the labelled SpaKC sample (10 μL) was first incubated with SpaA peptide (10 μL) of 16 different serial dilutions in the assay buffer for 5 min to allow binding. The samples were then loaded into Monolith NT.115 capillary (Nanotemper Technologies) and measured by using 20% (Auto-detect) Pico – RED as excitation power and medium MST power. The binding measurement was repeated three times. Data analysis was performed using Nanotemper affinity analysis software.

To measure the binding affinity between SpaKC-SpaF-SpaA_leader_, SpaF was incubated separately with SpaA_leader_ peptide of 16 different serial concentrations in assay buffer. The fluorescent-labelled SpaKC (10 μL) was mixed with these 16 samples and incubated for 5 min before loading into Monolith NT.115 capillary. The measurement condition and data analysis were the same as SpaKC-SpaAleader binding affinity measurement mentioned before.

### *In vitro* modification of SpaA_G-1K_ by SpaKC, SpaF and SpaKC-SpaF combination

Typically, SpaA peptide (25 μM) was incubated with SpaKC, SpaF or SpaKC-SpaF (12.5 μM) in reaction buffer (500 mM Na_2_CO_3_/NaHCO_3_, 200 mM NaCl, 1mM TCEP, pH 9.0). 5mM ATP and 5mM MgCl_2_ were supplied when SpaKC is present. All assays were carried out at 28 °C for 4h before quenched with 1% formic acid. The supernatant was lyophilized, re-dissolved in H_2_O and desalted by a SPE column. The resulting sample was lyophilized, re-dissolved and digested by trypsin before analyzed by LC-MS/MS.

