## Supplementary information for "Lanthipeptide Synthetases Participate the Biosynthesis of 2-Aminovinyl-Cysteine Motifs in Thioamitides"

### General methods and materials

Primers, genes and peptide were synthesized by Genscript Biotech (Nanjing, China). Restriction endonucleases and T4 DNA ligase were purchased from New England Biolabs (Ipswich, MA, USA). Phanta® Max Master Mix were purchased from Vazyme Biotech Co., Ltd (Nanjing, China). Medium components for bacterial cultures were purchased from Thermo Fisher (Waltham, MA, USA). Chemicals were purchased from Aladdin Reagent (Shanghai, China) and Sigma-Aldrich (Schnelldorf, Germany). Endoprotease trypsin was purchased from Roche Biosciences (Basel, Switzerland). *E. coli* DH5α was used as a host for cloning and plasmid propagation, and *E. coli* BL21 (DE3) was used as a host for expression of proteins and peptides. *S. sparsogenes* ATCC 25498 is purchased from (China General Microbiological Culture Collection Center, CGMCC).

All polymerase chain reactions (PCR) were carried out on a C1000 Touch™ thermal cycler (Bio-Rad). DNA sequencing was performed by the Genscript Biotech. Matrix-assisted laser desorption/ionization time-of-flight mass spectrometry (MALDI-TOF MS) was carried out on Bruker UltraFlextreme. Liquid chromatography electrospray ionization tandem mass spectrometry (LC/ESI-MS/MS) was carried out and processed using a Triple TOF 4600 System (AB SCIEX) equipped with a Prominence Ultra Fast Liquid Chromatography (UFLC) system (Shimadzu). UV-Vis spectrometry was conducted using Cary 300 (Agilent Technologies).

Conditions for all ESI-MS and MS/MS were set as follows: nebulizer gas: 55 psi; heater gas: 55 psi; curtain gas: 35 psi; drying temperature: 550 °C; ion spray voltage: 5500 V; declustering potential: 100 V; collision energy: 35 V (positive); collision energy spread: 10 V. The mass range and accumulation time are 400-4000 m/z, 250 ms for ESI-MS and 100-2000 m/z, 100 ms for MS/MS, respectively. Collision-induced dissociation (CID) was performed for fragmentation of the respective peptide ions. Calibration solutions purchased from AB SCIEX were used for instrument calibration, and high resolution was chosen in the ESI+ mode

### Primers used in this study

| Primer name | Sequence (5'-3') |
|---|---|
| SpaA_BamHI_F | GTTAGGATCCAATGTCTGAAGCAATGGTTTCCGCGGTCGAC |
| SpaA_HindIII_R | CATTAAGCTTTCAACAGTGGAAGGCCACCGTCGCGGCAG |
| $SpaA_{G-1K}$_HindIII_R | GTTCAAGCTTTCAACAGTGGAAGGCCACCGTCGCGGCAGCCGCCATCACGCTTTTCTGC |
| SpaF_NdeI_F | GTTACATATGAGCGAACGCGCCGACGGCGCC |
| SpaF_KpnI_R | CATAGGTACCTCACTGTCCTCGAGGGTCGGTGCGGCGG |
| $SpaF_{H85A}$_F | ACAAGGGCGGAGGCGCCGCCGCGGTCGCCCTCGGCGCCTGGGCGGATG |
| $SpaF_{H85A}$ _R | CCGCCCAGGCGCCGAGGGCGACCGCGGCGGCGCCTCCGCCCTTGTGGTC |
| SpaKC_ NdeI_F | GACTCATATGAGGAAACCGTACGGATTCACTGTCGCCGACCCGGACTTCTAC |
| SpaKC_ XhoI_R | GAACCTCGAGTCAGGACCGTGCGGCGATCGCGTCCACGGTGAAGAACGGATC |
| $SpaKC_{R159A}$_F | GGCGAGTCCCGCACGGTCCACTACGCGTACGGGGCCTACCGGAGGCTGGAA |
| $SpaKC_{R159A}$_R | TTCCAGCCTCCGGTAGGCCCCGTACGCGTAGTGGACCGTGCGGGACTCGCC |
| $SpaKC_{D362}$_F | TCGGCTTCGTCTTCGTCGCGGTCAGCCCCGGCAATGTGATGGCCA |
| $SpaKC_{R159A}$_R | TGGCCATCACATTGCCGGGGCTGACCGCGACGAAGACGAAGCCGA |
| $SpaKC_{600aa}$_ XhoI_R | GAATCTCGAGTTACGTGGTGAGGGGATGGAGGTCGGCCGTG |
| TvaA_BamHI_F | AGCTGGATCCAATGACCGAGAAGACCCAGATCACCGAC |
| TvaA_HindIII_R | GATTAAGCTTTTAGCAGTGCAGCGCAATGCTCGCC |
| TvaF_NdeI_F | AGCTCATATGGCGGAGCATGATGCGGCGGCGGAAG |
| TvaF_XhoI_R | AGCCCTCGAGTTAATCGCTCGCGCTTTTACGTTCCGCC |
| MutA_BamHI_F | GTTAGGATCCAATGTCTGAAACCACGACAGCAGCTCAGGTG |
| MutA_HindIII_R | GTAGAAGCTTTCAGCAGTGGTAAGCCACGGTGGCGATG |

| | |
|---|---|
| MutF_NdeI_F | AGCTCATATGGGCGGCATCCGACCAGCCAG |
| MutF_XhoI_R | TAGCCTCGAGTCACTGTCCTTCTGCTCGCGCGGCGTG |
| TsdA_BamHI_F | GTTAGGATCCGATGAGCGAAGCGATGGCGAGCGCGGTGGATGAAGCGGCGTTTG |
| TsdA_HindIII_R | CAAGCTTTTAGCAGTGAAACGCAACGGTCGCCGCCGCCGCCATCAC |
| TsdA$_{G-1K}$_F | ATCAGCCCGGACGAGGAAGCGCAGAAAAGCGTGATGGCGGCGGCGGCGACC |
| TsdA$_{G-1K}$ _R | GGTCGCCGCCGCCGCCATCACGCTTTTCTGCGCTTCCTCGTCCGGGCTGAT |
| TsdF_MfeI_F | GTTTACAATTGATGGGCACCGACCACGCCACCGGCCCGCAGCAGGCAC |
| TsdF_XhoI_R | CTTACTCGAGTTACTGTCCTTGTGGGTGGGTGCGTCGTGCAGCTGCCTGGACC |
| TsdKC_NdeI_F | GTTACATATGATGAGCCTGCGTCATATTGCGTACTGCCCGCC |
| TsdKC_XhoI_R | CATACTCGAGTTAACGACCCGCCGCAACCGGCAGACGCAG |
| MalA_BamHI_F | GATAGGATCCGATGGTTAGCGCGGTTGACGAGACCGCGTTCGCGG |
| MalA_HindIII_R | CAACAAGCTTTTAGCAGTGAAACGCAACGGTCGCCGCCGCCGCC |
| MalA$_{G-1K}$_F | CCGGATGAGGAAGCGCAAAAAAGCGTGATGGCGGCGGCGG |
| MalA$_{G-1K}$ _R | GTCGCCGCCGCCGCCATCACGCTTTTTTGCGCTTCCTCATC |
| MalF_NdeI_F | GTTACATATGAGCGAGCGTGCGAGCGGTCCGCAGGAAG |
| MalF_XhoI_R | GTTACTCGAGTTATTGACCACGCGGGTGGGTCAGAC |
| AlbA_BamHI_F | CTAGGGATCCGATGAACAGCCACGACGATGCGGTGGCGCCGAG |
| AlbA_HindIII_R | GCATAAGCTTTTAGCAGTGAACCGCAATGGTCACC |
| AlbF_NdeI_F | GTTACATATGAGCACCGGTCGTGCGCGTCACACCAG |
| AlbF_XhoI_R | GTCACTCGAGTTACAGACGATCACCGTGACGGCTG |

**Production and analysis of thiosparsoamide by *S. sparsogenes* ATCC 25498**

*S. sparsogenes* ATCC 25498 was spread on PS5 agar plates for sporulation and growth. Approximately 1 $cm^2$ of the agar with sporulated *S. sparsogenes* ATCC 25498 was used to inoculate 100 mL of the fermentation medium. After incubation at 28°C and 220 rpm for 96 hours, the whole broth was extracted with methanol, lyophilized, and re-dissolved with methanol. The resulting sample was subjected to LC-HRMS analysis for thiosparsoamide detection.

PS5 agars: 20 g of starch, 5 g of pharmamedia, and 20 g of agar per liter (pH 7.0).

Fermentation medium: 25 g of glucose, 15 g of soybean meal, 2 g of dry yeast, and 4 g of CaCO3 per liter (pH7.0)。

LC-MS analysis of thiosparsoamide was carried out with liquid chromatography heated electrospray ionization tandem mass spectrometry (LC/HESI-MS/MS) and processed using a Thermo Scientific™ Q Exactive™ equipped with a Vanquish Duo UHPLC system (Thermo sciienctific). UV-Vis spectrometry was conducted using Diode Array Detector FG.

Conditions for HESI-MS and MS/MS were set as follows: sheath gas: 45 psi; auxiliary gas: 15 psi; spare gas: 2.0 psi; capillary temperature: 320 °C; spray voltage: 3500 V; normalized collision energy: 30% of the available 5V (positive); The mass range and accumulation time are 200-2000 m/z, 100 ms for HESI-MS. And data dependent top 20 ion were select by quadrupole selected and HCD (higher collisional dissociation) was performed for fragmentation of the respective ion.

**Molecular cloning of *spaA, spaF, spaKC, tvaA, tvaF, mutA, mutF, tsdA, tsdF, tsdKC, malA, malF, albA and albF* genes.**

Plasmids containing target genes were synthesized by Genscript Biotech and PCR-amplified by 30 cycles of denaturing (95 °C for 30 s), annealing (65 °C for 30 s), and extending (72 °C, 1 min/kb) using high fidelity Phanta DNA Polymerase. Amplifications were confirmed by 2% agarose gel electrophoresis, and the PCR products were purified using an Omega Biotech Cycle Pure Kit. Target DNA fragments and selected vectors were digested in separate reactions with selected pair of restriction enzymes for 2 h at 37 °C. The digested products were purified by agarose gel electrophoresis, and the DNA fragments were extracted from the gel using an Omega Biotech Gel Extraction Kit. The resulting DNA products were ligated at 16 °C for 12 h in T4 DNA Ligase buffer with T4 DNA Ligase. *E. coli* DH5α cells were transformed with 2.5 μL of the ligation product by heat shock, and cells were plated on proper antibiotics LB-agar plates and grown for 16 h at 37 °C. For each plasmid transformant, a single colony was picked to inoculate 5 mL culture of LB medium with proper antibiotics. The cultures were grown at 37 °C for 12 h, and plasmids were isolated using an Omega Biotech Plasmid Mini Kit. The sequences of the resulting plasmid products were confirmed by DNA sequencing.

**Construction of plasmids for coexpression of (His$_6$-SpaA)-SpaF, (His$_6$-TvaA)-TvaF, (His$_6$-MutA)-MutF, (His$_6$-TsdA)-TsdF, (His$_6$-MalA)-MalF and (His$_6$-AlbA)-AlbF**

Amplification of the two genes was performed using high fidelity Phanta® DNA Polymerase following the protocol described above. Restriction digestions were performed with selected pair of restriction enzymes following standard protocol. The digested products of genes were purified by agarose gel electrophoresis and ligated with pRSFDuet-1 vector. The resulting DNA products were ligated at 16 °C for 12 h in 1X T4 DNA Ligase buffer with T4 DNA Ligase (0.7 U/ μL). *E. coli* DH5α cells were transformed with 2.5 μL of the ligation product by heat shock, and cells were plated on LB-kanamycin agar plates and grown for 16 h at 37 °C. Several colonies were picked and used to inoculate separate 5 mL cultures of LB-kanamycin medium. The cultures were grown at 37 °C for 12 h, and plasmids were isolated using a Omega Biotech Plasmid Mini Kit. The sequences of the resulting plasmid products were confirmed by DNA sequencing.

**Mutagenesis of SpaA, SpaF, SpaKC, TsdA and MalA(136) genes**

Mutation of SpaA was carried out by 30 cycles of denaturing (95 °C for 30 s), annealing (65 °C for 30 s), and extending (72 °C, 1 min/kb) using high fidelity Phanta® DNA Polymerase. Amplifications were confirmed by 1% agarose gel electrophoresis, and the PCR products were purified using an Omega Biotech Cycle Pure Kit. The target DNA fragment was digested in reaction containing 1X NEB buffer (New England Biolabs) with DpnI for 3 h at 37 °C. *E. coli* DH5α cells were transformed with 10 μL of the digested product by heat shock, and cells were plated on LB- kanamycin agar plates and grown for 16 h at 37 °C. Several colonies were picked and used to inoculate separate 5 mL cultures of LB-kanamycin medium. The cultures were grown at 37 °C for 12 h, and plasmids were isolated using an Omega Biotech Plasmid Mini Kit. The sequences of the resulting plasmid products were confirmed by DNA sequencing. The other mutations were carried out by the protocol described above.

**Overexpression and purification of SpaF-modified SpaA peptides**

*E. coli* BL21(DE3) cells were transformed with pRSFDuet-1-(His$_6$-SpaA)-SpaF, pRSFDuet-1-(His$_6$-SpaA$_{G-1K}$)-SpaF and plated on a Luria broth (LB) agar plate containing 50 mg/L of kanamycin. A single colony was used to inoculate a 5 mL culture of LB supplemented with 50 mg/L of kanamycin. The culture was grown at 37 °C for 12 h and was used to inoculate 4 L of LB containing 50 mg/L of kanamycin. Cells were grown at 37 °C to $OD_{600}$=0.6-0.8 before IPTG was added to a final concentration of 0.2 mM and the culture was incubated at 18 °C for additional 16 h. Cells were harvested by centrifugation at 12,000 ×g for 25 min at 4 °C. The resulting cell pellet was resuspended in 30 mL of start buffer (20 mM $NaH_2PO_4$, pH 7.5, 500 mM NaCl, 0.5 mM imidazole, 20% glycerol), and the suspension was sonicated on ice for 20 min to lyse the cells. Cell debris was removed by centrifugation at 23,700 ×g for 30 min at 4 °C. The supernatant was discarded, and the pellet containing peptide products was resuspended in 30 mL of start buffer. The sonication and centrifugation steps were repeated. Again, the supernatant was discarded, and the pellet was resuspended in 30 mL of buffer 1 (6 M guanidine HCl, 20 mM $NaH_2PO_4$, pH 7.5, 500 mM NaCl, 0.5 mM imidazole). The sample was sonicated and insoluble material was removed by centrifugation at 23,700 ×g for 30 min at 4 °C, followed by filtration of the supernatant through a 0.45 μm filter. The filtered sample was applied to a 5 mL HisTrap HP (GE Healthcare Life Sciences) immobilized metal affinity chromatography (IMAC) column previously charged with $NiSO_4$ and equilibrated in buffer 1. The column was washed with two column volumes of buffer 1, followed by two column volumes of buffer 2 (4 M guanidine HCl, 20 mM $NaH_2PO_4$ , pH 7.5, 500 mM NaCl, 30 mM imidazole). The peptide was eluted with 1-2 column volumes of elution buffer (4 M guanidine HCl, 20 mM $NaH_2PO_4$, pH 7.5, 500 mM NaCl, 1 M imidazole). The fractions were desalted using a Sep-Pak® C18 Cartridges and analyzed by MALDI-TOF MS and the organic solvents were removed by rotary evaporation, followed by lyophilization. The product was kept at −80 °C for long-term storage. Typical yields from 4 L culture were 1 mg for $His_6$-SpaA peptides.

**Overexpression of SpaKC with thioamitide precursor peptides.**

*E. coli* BL21(DE3) cells were transformed with pRSFDuet-1-($His_6$-thioamitide precursor peptide) and pACYCDuet-1-SpaKC and plated on a Luria broth (LB) agar plate containing 50 mg/L of kanamycin and 35 mg/L of chloramphenicol. A single colony was used to inoculate a 5 mL culture of LB supplemented with 50 mg/L of kanamycin and 35 mg/L of chloramphenicol at 37 °C for 12 h. The culture was used to inoculate 4 L of LB containing 50 mg/L of kanamycin and 35 mg/L of chloramphenicol. Cells were grown at 37 °C to $OD_{600}$=0.6-0.8 before IPTG was added to a final concentration of 0.2 mM and the culture was incubated at 18 °C for another 16 h before harvesting. The purification procedure was described as above.

**Overexpression and purification of (SpaKC-SpaF)-modified SpaA peptides.**

*E. coli* BL21(DE3) cells were transformed with pRSFDuet-1-$His_6$-SpaA-SpaF and pACYCDuet-1-SpaKC or pACYCDuet-1-$SpaKC_{mutants}$ and plated on a Luria broth (LB) agar plate containing 50 mg/L of kanamycin and 35 mg/L of chloramphenicol. A single colony was used to inoculate a 5 mL culture of LB supplemented with 50 mg/L of kanamycin and 35 mg/L of chloramphenicol at 37 °C for 12 h. The culture was used to inoculate 4 L of LB containing 50 mg/L of kanamycin and 35 mg/L of chloramphenicol. Cells were grown at 37 °C to $OD_{600}$~0.6-0.8 before IPTG was added to a final concentration of 0.2 mM and the culture was incubated at 18 °C for another 16 h before harvesting. The purification procedure was described as above.

**Overexpression and purification of (SpaKC-$LanD_t$)-modified thioamitide precursor peptides**

The coexpression of SpaKC, TvaF and TvaA peptide is used as a representative example.

*E. coli* BL21(DE3) cells were transformed with pRSFDuet-1-($His_6$-TvaA)-TvaF and pACYCDuet-1-SpaKC plated on a Luria broth (LB) agar plate containing 50 mg/L of kanamycin and 35 mg/L of chloramphenicol. A single colony was used to inoculate a 5 mL culture of LB supplemented with 50 mg/L of kanamycin and 35 mg/L of chloramphenicol at 37 °C for 12 h. The culture was used to inoculate 4 L of LB containing 50 mg/L of kanamycin and 35 mg/L of chloramphenicol. Cells were grown at 37 °C to $OD_{600}$=0.6-0.8 before IPTG was added to a final concentration of 0.2 mM and the culture was incubated at 18 °C for another 16 h before harvesting. The purification procedure was described as above. The other coexpression were carried out by the protocol described above.

**Determination of the flavin cofactor bound to SpaF.**

To identify the type of flavin cofactor, an aliquot of 200 µL of a 1.2 mM solution of $His_6$-SpaF was denatured at 100 °C for 10 min. Precipitated protein was then removed by centrifugation (10,000 ×g, 10 min, 25 °C) and the released flavin was purified by Sep-Pak® C18 Cartridges and analyzed by LC-ESI-MS.

**Modification of SpaA peptides with IAA.**

To determine the presence of free cysteine, SpaA peptides was digested by trypsin and then incubated with 20 mM IAA at 37 °C for 3h in 50 µL of reaction buffer (50 mM Tris, pH 8.6, 1 mM TCEP) in the dark, after which the reaction was quenched by addition of 2 µL of 1 M DTT to eliminate excess IAA and prevent further side reactions. The reactions were further analyzed by MALDI-TOF.

**Reductive desulfurization of modified SpaA peptides**

(SpaKC-SpaF)-modified $SpaA_{G\text{-}1K}$ (1.0 mg) was suspended in 4.0 mL of $CH_3OH/H_2O$ (1:1), to which 20 mg of $NiCl_2$ and 20 mg of $NaBH_4$ were added. This mixture was stirred under 1 atm of $H_2$ at room temperature. After 8.5 h, the mixture was taken off the hydrogenation apparatus, and then centrifuged, followed by the removal of the supernatant. Mixed solvent of $CH_3OH/H_2O$ (ration=1:1, 2.0 mL) was added to the black nickel boride pellets, and the suspension was sonicated for 15 minutes. The suspension was centrifuged, and the supernatants were collected. The organic solvent was removed in vacuo, and the desulfurized peptide was treated with trypsin.
Digestion conditions: 20 mM Tris-HCl, pH = 8.0, 1 µM trypsin, 100 µM peptide product at 37 °C for 6 h.

**Measurement of protein-protein and protein-peptide interactions by MST assays**

As a representative example, the binding affinity between SpaKC and $SpaA_{leader}$ was measured using Monolith NT.115 Pico (Nanotemper Technologies). SpaKC was fluorescently labelled by the labeling kit (Monolith Protein Labeling Kit RED-NHS 2nd Generation (Amine Reactive), MO-L011). A volume of 90 µL SpaKC sample (10 µM) in the labelling buffer (130 mM $NaHCO_3$, 50 mM NaCl, pH 8.2) was mixed with 10 µL dye (Dye RED-NHS 2nd Generation (Nanotemper MO-L011)) solution (300 µM) for 30 min at room temperature in the dark. Next, the SpaKC sample was loaded to column B (Nanotemper MO-L011) and eluted with 450 µL of assay buffer (50mM $Na_2CO_3/NaHCO_3$, 150mM NaCl, 10mM $MgCl_2$, 0.07% tween 20, pH 9.0). The labelled SpaKC sample was diluted until pretest fluorescence excitation range from 16000-20000 for the most optimized response (~1nM). To perform the MST assay, the labelled SpaKC sample (10 µL) was first incubated with SpaA peptide (10 µL) of 16 different serial dilutions in the assay buffer for 5 min to allow binding. The samples were then loaded into Monolith NT.115 capillary (Nanotemper Technologies) and measured by using 20% (Auto-detect) Pico – RED as excitation power and medium MST power. The binding measurement was repeated three times. Data analysis was performed using Nanotemper affinity analysis software.

To measure the binding affinity between SpaKC-SpaF-$SpaA_{leader}$, SpaF was incubated separately with $SpaA_{leader}$ peptide of 16 different serial concentrations in assay buffer. The fluorescent-labelled SpaKC (10 µL) was mixed with these 16 samples and incubated for 5 min before loading into Monolith NT.115 capillary. The measurement condition and data analysis were the same as SpaKC-SpaAleader binding affinity measurement mentioned before.

***In vitro* modification of $SpaA_{G\text{-}1K}$ by SpaKC, SpaF and SpaKC-SpaF combination**

Typically, SpaA peptide (25 µM) was incubated with SpaKC, SpaF or SpaKC-SpaF (12.5 µM) in reaction buffer (500 mM $Na_2CO_3/NaHCO_3$, 200 mM NaCl, 1mM TCEP, pH 9.0). 5mM ATP and 5mM $MgCl_2$ were supplied when SpaKC is present. All assays were carried out at 28 °C for 4h before quenched with 1% formic acid. The supernatant was lyophilized, re-dissolved in $H_2O$ and desalted by a SPE column. The resulting sample was lyophilized, re-dissolved and digested by trypsin before analyzed by LC-MS/MS.

| Strain | LanKC homologs | Accession number | NCBI Reference Sequence |
|---|---|---|---|
| *Streptomyces sparsogenes* ATCC 25498 | SpaKC | WP_065963996.1 | NZ_MAXF01000094.1 |
| *Streptomyces sp.* MUSC 125 | SmuKC(125) | WP_039650135.1 | NZ_JUIG01000010.1 |
| *Streptomyces sp.* MUSC 14 | SmuKC(14) | WP_071371419.1 | NZ_MLYN01000014.1 |
| *Streptomyces malaysiense* MUSC 136 | MalKC(136)-1 | OIK27458.1 | LBDA02000024.1 |
| | MalKC(136)-2 | OIK27459.1 | LBDA02000024.1 |
| *Streptomyces malaysiense* | MalKC | WP_071381943.1 | NZ_MLYO01000031.1 |
| *Streptomyces sp.* NRRL S-87 | TsdKC-1 | WP_037887254.1 | NZ_JOGB01000003.1 |
| | TsdKC-2 | WP_107048477.1 | NZ_JOGB01000016.1 |
| *Streptomyces sp.* CNB091 | ScnKC-1 | WP_026290248.1 | NZ_KB898991.1 |
| | ScnKC-2 | WP_018959331.1 | NZ_KB899001.1 |
| *Streptomyces sp.* NRRL S-4 | SnrKC(S-4) | WP_053928640.1 | NZ_LGKJ01000001.1 |
| *Streptomyces mutomycini strain* NRRL B-65393 | MutKC | WP_065848356.1 | NZ_MAPV01000093.1 |
| *Amycolatopsis alba* DSM 44262 | AmyKC | WP_020633044.1 | NZ_NMQU01000026.1 |
| *Nocardiopsis potens* DSM 45234 | NocKC | WP_017595403.1 | NZ_ANBB01000057.1 |

**Table S1.** $LanKC_t$ genes discovered in the genomes of thioamitide-producing strains. All genes are located outside corresponding thioamitide BGCs.

| Strain | LanKC homologs | Accession number | NCBI Reference Sequence |
|---|---|---|---|
| *Streptomyces lividans TK24* | LivKC | AIJ12038.1 | CP009124.1 |
| *Streptomyces coelicolor M1146(S-4)* | CoeKC | TDY69100.1 | SORM01000032.1 |
| *Streptomyces avermitilis SUKA24(17)* | SukaKC | WP_010988898.1 | NZ_MLYN01000014.1 |

**Table S2.** LanKC homologs from the genomes of streptomyces strains as heterologous hosts for thioviridamides.

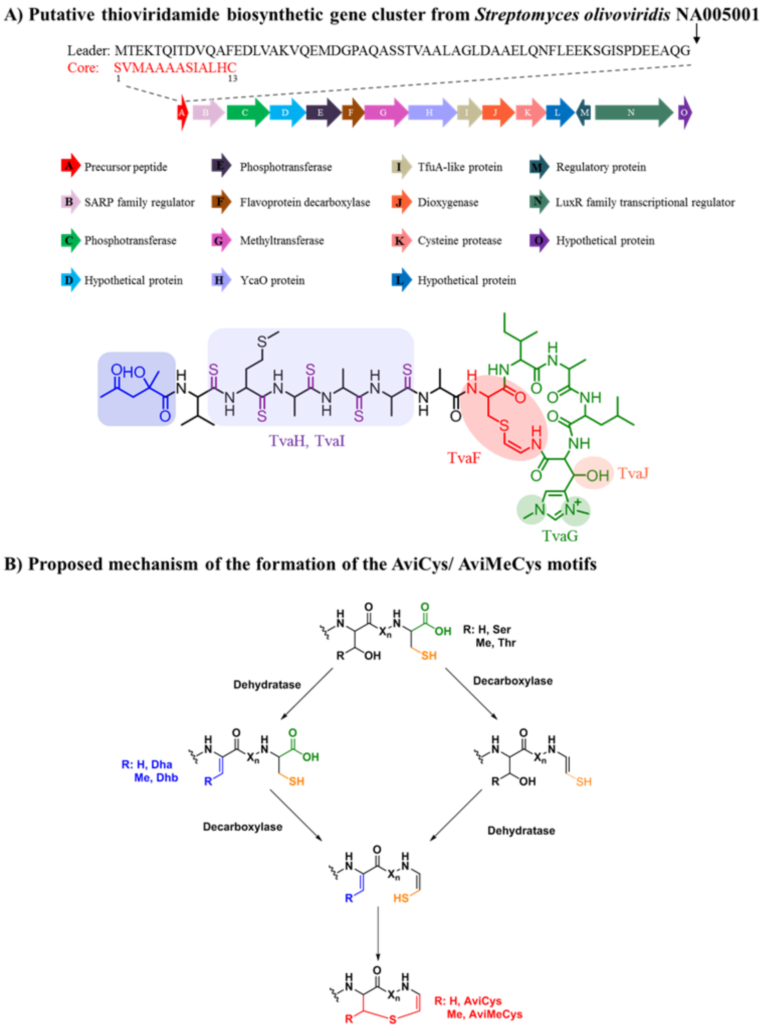


**Figure S1.** Putative thioviridamide biosynthetic gene cluster and the proposed formation of AviCys/AviMeCys motif. (A) Putative thioviridamide biosynthetic gene cluster from *Streptomyces olivoviridis* NA005001. (B) Proposed mechanism of the formation of AviCys/ AviMeCys motifs.

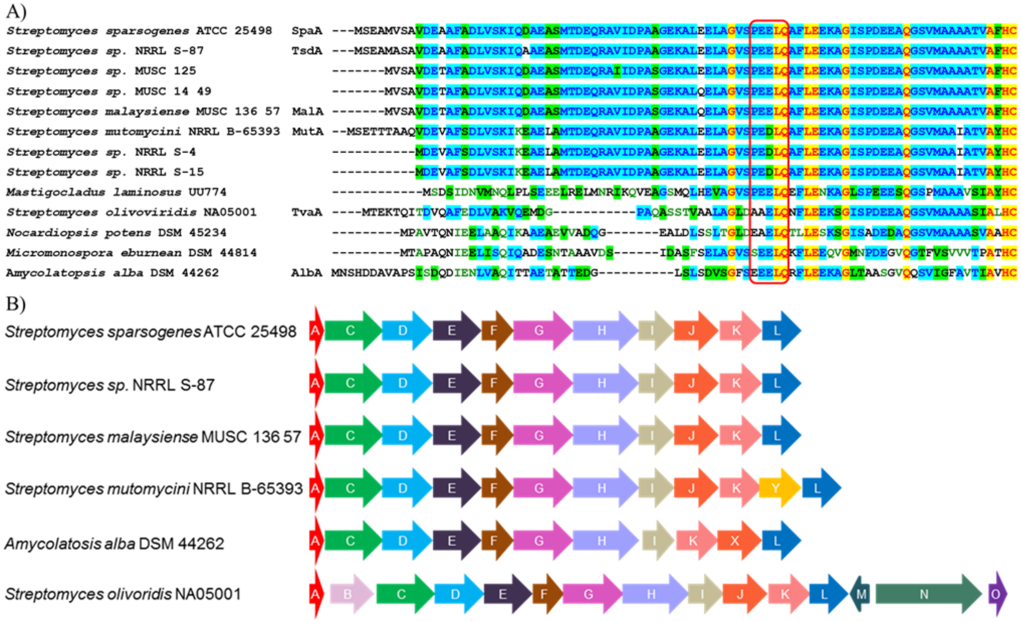


**Figure S2.** Bioinformatic analysis of thioamitide BGCs. (A) Sequence alignment of thioamitide precursor peptides. (B) Putative thiosparsoamide biosynthetic gene cluster (top) from *Streptomyces sparsogenes* ATCC 25498 compared to homologous gene clusters from other organisms. The putative function of each gene is listed as follows:A, precursor peptide; B, putative SARP family regulator; C, phosphotransferase; D, hypothetical protein; E, phosphotransferase; F, putative flavoprotein decarboxylase; G, putative methyltransferase; H, YcaO protein; I, TfuA-like protein; J, putative dioxygenase; K, Cysteine proteinase; L, hypothetical protein; M, putative regulatory protein; N, putative LuxR family transcriptional regulator; O, hypothetical protein; Y, putative methyltransferase.

The accession numbers of thioamitide precursor peptides in GenBank are listed as follows: SpaA (WP_104531249.1), TsdA (WP_107048385.1), precursor peptide form *Streptomyces sp.* MUSC 125 (KIE26472.1), precursor peptide form *Streptomyces sp.* MUSC 14 49 (OIJ91358.1), MalA (ATJ00792.1), MutA (WP_107418454.1), precursor peptide form *Streptomyces sp.* NRRL S-4 (KPC82025.1), precursor peptide form *Streptomyces sp.* NRRL S-15 (WP_031090922.1), precursor peptide from *Mastigocladus laminosus* UU774 (WP_044449650.1), TvaA (BAN83916.1), precursor peptide form *Nocardiopsis potens* DSM 45234 (WP_017595621.1), precursor peptide form *Micromonospora eburnean* DSM 44814 (SCL51081.1), AblA (WP_020634202.1).

**A) Detection of thiosparsoamide by LC-MS.**

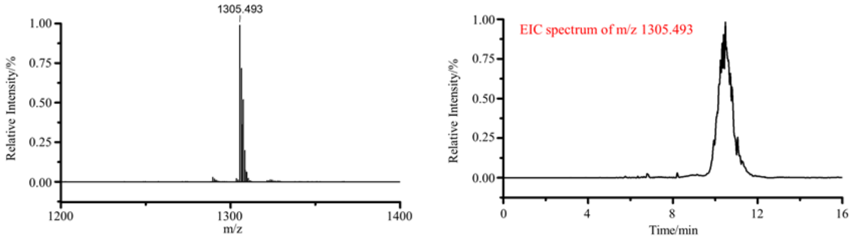


**B) MS/MS analysis of thiosparoamide**

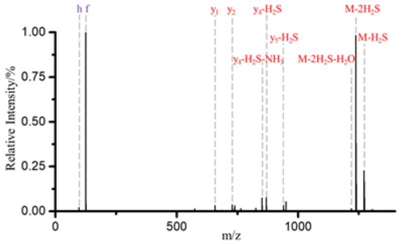


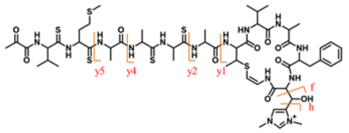


| | theoretical m/z | Observed m/z | relative deviation (ppm) |
|---|---|---|---|
| h | 97.0760 | 97.0761 | 1.0301 |
| f(retro aldol) | 125.0709 | 125.0707 | -1.5991 |
| y1 | 657.3177 | 657.3116 | -9.2801 |
| y2 | 728.3548 | 728.3613 | 8.9242 |
| y4-$H_2S$-$NH_3$ | 851.3691 | 851.3707 | 1.8793 |
| y4-$H_2S$ | 868.3957 | 868.3996 | 4.4910 |
| y5-$H_2S$ | 939.4292 | 939.4314 | 2.3418 |
| M-2$H_2S$-$H_2O$ | 1219.5032 | 1219.4880 | -12.4641 |
| M-2$H_2S$ | 1237.5137 | 1237.4956 | -14.6261 |
| M-$H_2S$ | 1271.5015 | 1271.4984 | -2.4381 |

**Figure S3.** Production of thiosparsoamide by *S. sparsogenes* ATCC 25498 and its structural characterization. (A) Detection of thiosparsoamide by LC-MS. (B) MS/MS analysis of thiosparsoamide.

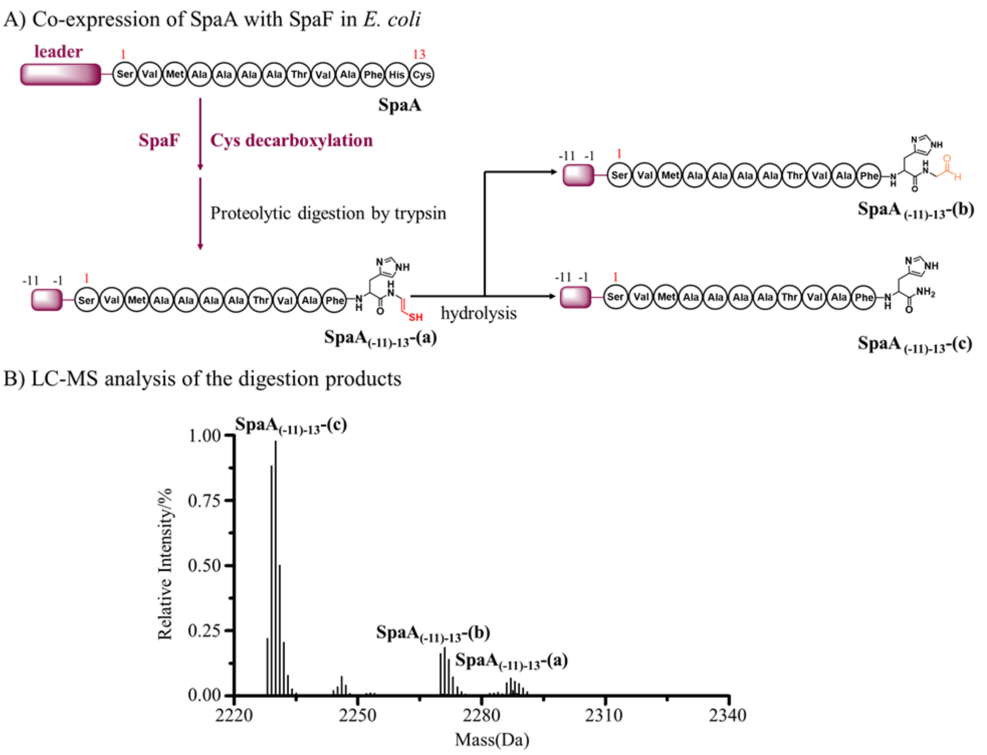


**Figure S4.** SpaA was oxidatively decarboxylated by SpaF by co-expression in *E. coli*. (A). SpaA was modified by SpaF by generating unstable decarboxylation product **SpaA-(a)**, which was further hydrolyzed to **SpaA-(b)** and **SpaA-(c)**. (B) LC-MS analysis of digestion product **SpaA$_{(-11)-13}$-(a)**, **SpaA$_{(-11)-13}$-(b)** and **SpaA$_{(-11)-13}$-(c)**. Digestion conditions: 20 mM Tris-HCl, pH = 8.0, 1 µM trypsin, 100 µM peptide product at 37 °C for 6 h.

Monoisotopic mass of decarboxylation products:

**SpaA$_{(-11)-13}$-(a)**: calc. 2286.041, obs. 2228.069;

**SpaA$_{(-11)-13}$-(b)**: calc. 2270.064, obs. 2270.067;

**SpaA$_{(-11)-13}$-(c)**: calc. 2228.053, obs. 2228.058.

A) LC-MS/MS analysis of the **SpaA$_{(-11)-13}$-(b)**

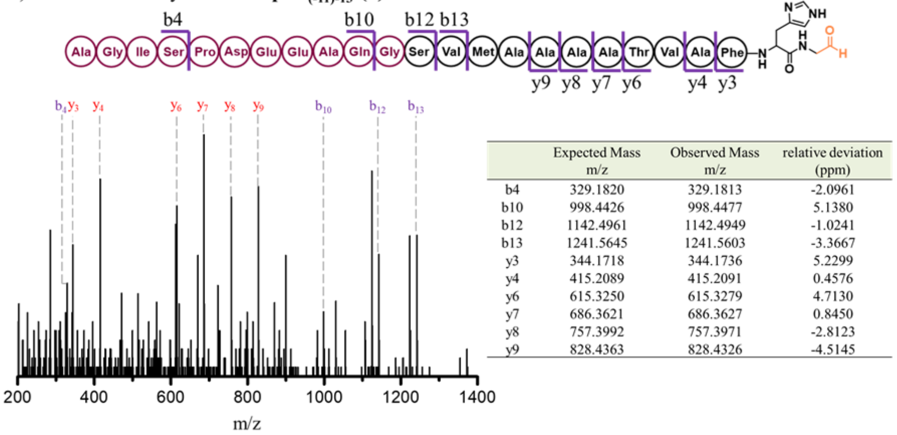


| | Expected Mass m/z | Observed Mass m/z | relative deviation (ppm) |
|---|---|---|---|
| b4 | 329.1820 | 329.1813 | -2.0961 |
| b10 | 998.4426 | 998.4477 | 5.1380 |
| b12 | 1142.4961 | 1142.4949 | -1.0241 |
| b13 | 1241.5645 | 1241.5603 | -3.3667 |
| y3 | 344.1718 | 344.1736 | 5.2299 |
| y4 | 415.2089 | 415.2091 | 0.4576 |
| y6 | 615.3250 | 615.3279 | 4.7130 |
| y7 | 686.3621 | 686.3627 | 0.8450 |
| y8 | 757.3992 | 757.3971 | -2.8123 |
| y9 | 828.4363 | 828.4326 | -4.5145 |

B) LC-MS/MS analysis of the **SpaA$_{(-11)-13}$-(c)**

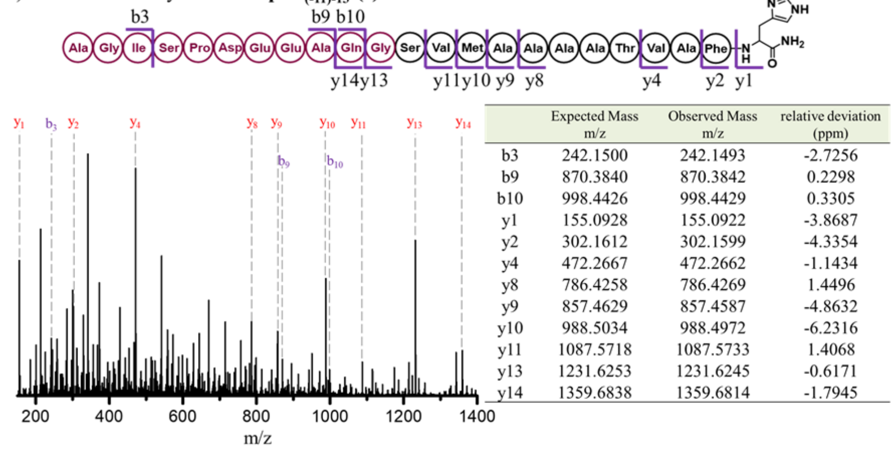


| | Expected Mass m/z | Observed Mass m/z | relative deviation (ppm) |
|---|---|---|---|
| b3 | 242.1500 | 242.1493 | -2.7256 |
| b9 | 870.3840 | 870.3842 | 0.2298 |
| b10 | 998.4426 | 998.4429 | 0.3305 |
| y1 | 155.0928 | 155.0922 | -3.8687 |
| y2 | 302.1612 | 302.1599 | -4.3354 |
| y4 | 472.2667 | 472.2662 | -1.1434 |
| y8 | 786.4258 | 786.4269 | 1.4496 |
| y9 | 857.4629 | 857.4587 | -4.8632 |
| y10 | 988.5034 | 988.4972 | -6.2316 |
| y11 | 1087.5718 | 1087.5733 | 1.4068 |
| y13 | 1231.6253 | 1231.6245 | -0.6171 |
| y14 | 1359.6838 | 1359.6814 | -1.7945 |

**Figure S5.** MS/MS analysis of **SpaA$_{(-11)-13}$-(b)** and **SpaA$_{(-11)-13}$-(c)** after trypsin digestion. (A) LC-MS/MS analysis of **SpaA$_{(-11)-13}$-(b)**. The *b* and *y* ions are listed in the table and marked in the spectrum. (B) LC-MS/MS analysis of **SpaA$_{(-11)-13}$-(c)**. The *b* and *y* ions are listed in the table and marked in the spectrum.

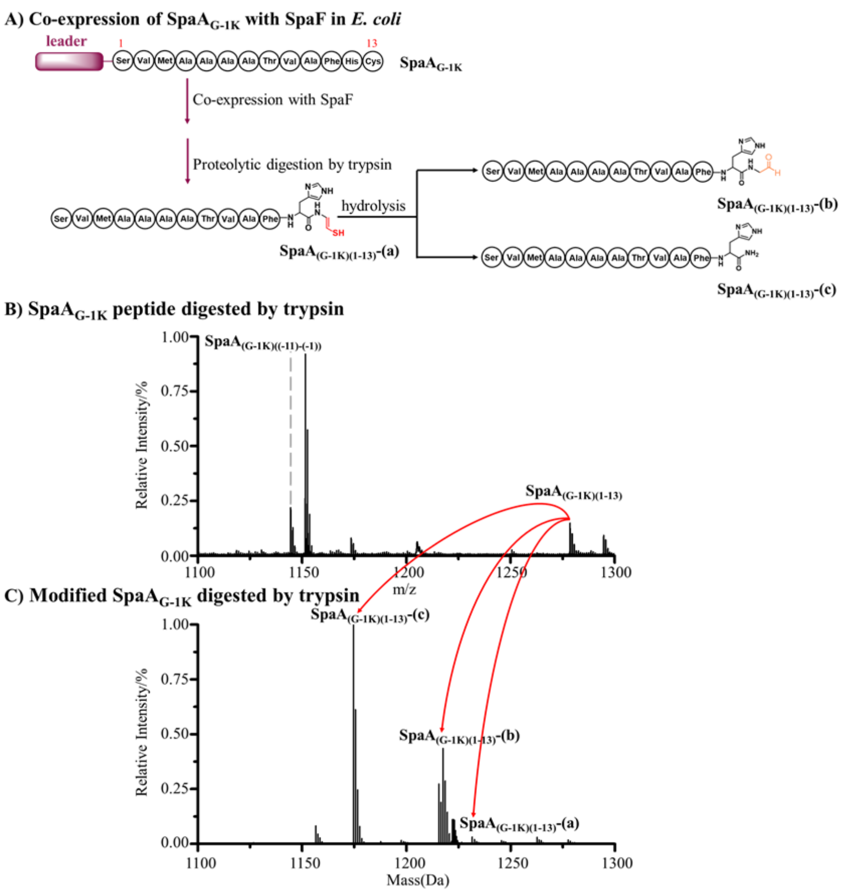


**Figure S6.** His$_6$-tagged SpaA$_{G\text{-}1K}$ was co-expressed with SpaF in *E. coli*, purified and digested by trypsin. (A). Co-expression of SpaF and SpaA$_{G\text{-}1K}$ generated decarboxylation products. **(**B) MALDI-TOF analysis of digestion products of SpaF modified SpaA$_{G\text{-}1K}$. (C) LC-MS analysis of product **SpaA$_{(G\text{-}1K)(1\text{-}13)}$-(a)**, **SpaA$_{(G\text{-}1K)(1\text{-}13)}$-(b)** and **SpaA$_{(G\text{-}1K)(1\text{-}13)}$-(c)**. Digestion conditions: 20 mM Tris-HCl, pH = 8.0, 1.0 µM trypsin, 100 µM peptide product at 37 °C for 6 h.

Monoisotopic mass of products:

**SpaA$_{(G\text{-}1K)(1\text{-}13)}$**: calc. 1278.59, obs. 1278.58;

**SpaA$_{(G\text{-}1K)((-11)\text{-}(-1))}$**: calc. 1144.55, obs. 1144.54.

**SpaA$_{(G\text{-}1K)(1\text{-}13)}$-(a)**: calc. 1231.584, obs. 1231.590;

**SpaA$_{(G\text{-}1K)(1\text{-}13)}$-(b)**: calc. 1215.607, obs. 1215.601;

**SpaA$_{(G\text{-}1K)(1\text{-}13)}$-(c)**: calc. 1173.596, obs. 1173.588.

A) LC-MS/MS analysis of the **SpaA$_{(G-1K)(1-13)}$-(b)**

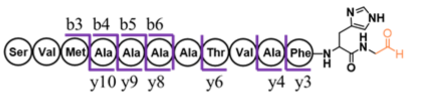


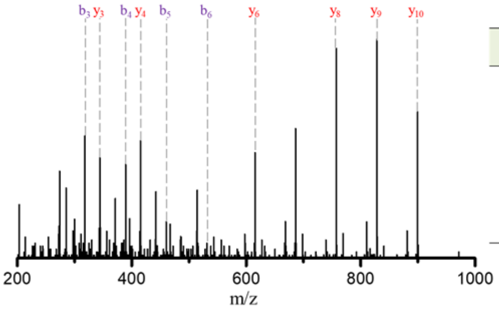


| | Expected Mass m/z | Observed Mass m/z | relative deviation (ppm) |
|---|---|---|---|
| b3 | 318.1483 | 318.1462 | -6.4435 |
| b4 | 389.1854 | 389.1834 | -5.0362 |
| b5 | 460.2225 | 460.2205 | -4.3023 |
| b6 | 531.2596 | 531.2594 | -0.3576 |
| y3 | 344.1718 | 344.1707 | -3.1961 |
| y4 | 415.2089 | 415.2085 | -0.9875 |
| y6 | 615.3250 | 615.3219 | -5.0380 |
| y8 | 757.3992 | 757.4003 | 1.4127 |
| y9 | 828.4363 | 828.4295 | -8.2565 |
| y10 | 899.4735 | 899.4659 | -8.4049 |

B) LC-MS/MS analysis of the **SpaA$_{(G-1K)(1-13)}$-(c)**

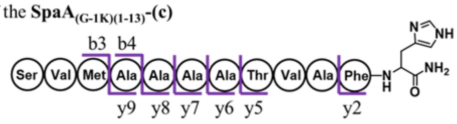


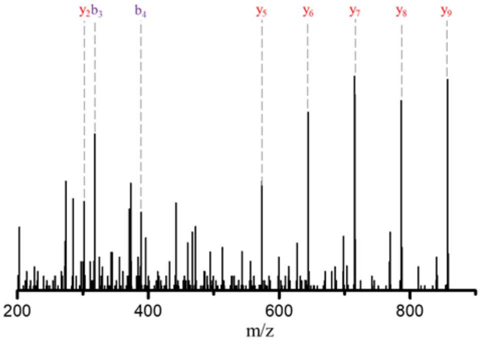


| | Expected Mass m/z | Observed Mass m/z | relative deviation (ppm) |
|---|---|---|---|
| b3 | 318.1483 | 318.1493 | 3.3003 |
| b4 | 389.1854 | 389.1845 | -2.2097 |
| y2 | 302.1612 | 302.1605 | -2.3497 |
| y5 | 573.3144 | 573.3112 | -5.6165 |
| y6 | 644.3515 | 644.3481 | -5.3232 |
| y7 | 715.3886 | 715.3854 | -4.5290 |
| y8 | 786.4258 | 786.4227 | -3.8910 |
| y9 | 857.4629 | 857.4579 | -5.7962 |

**Figure S7.** LC-MS/MS analysis of **SpaA$_{(G-1K)(1-13)}$-(b)** and **SpaA$_{(G-1K)(1-13)}$-(c)**. A) LC-MS/MS analysis of **SpaA$_{(G-1K)(1-13)}$-(b)**. (B) LC-MS/MS analysis of **SpaA$_{(G-1K)(1-13)}$-(c)**. The *b* and *y* ions are listed in the table and marked in the spectrum.

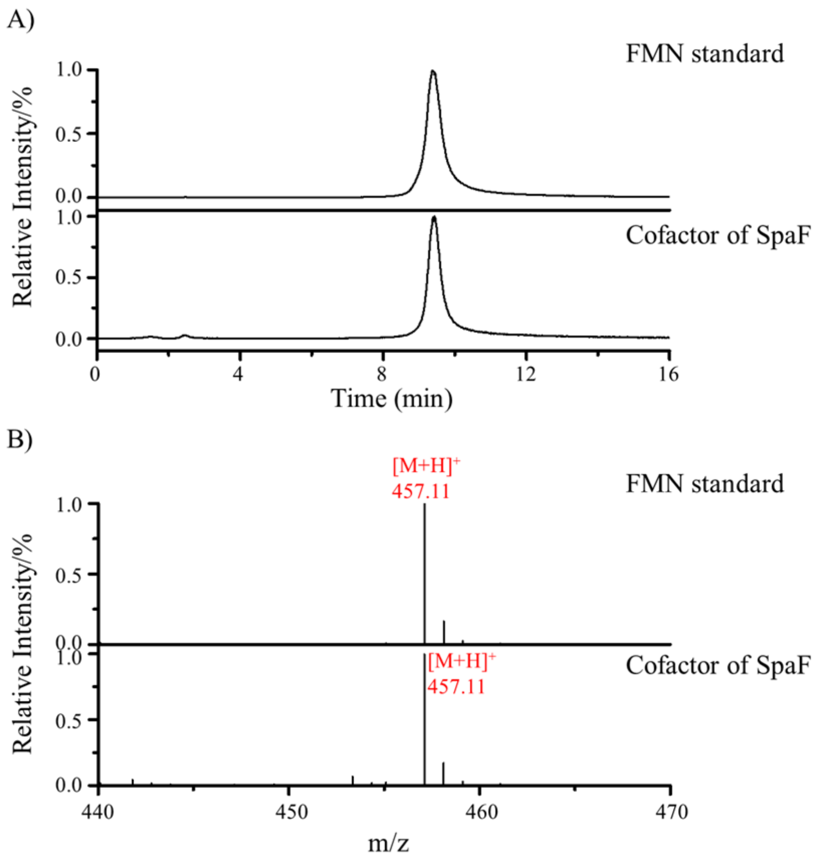


**Figure S8.** Characterization of the cofactor bound with SpaF. (A) EIC trace of FMN standard and the cofactor extracted from denatured SpaF. (B) The cofactor isolated from SpaF has the same mass as FMN, as determined by LC-MS analysis. Expected monoisotopic mass of FMN 457.11 Da, observed monoisotopic mass 457.11 Da.

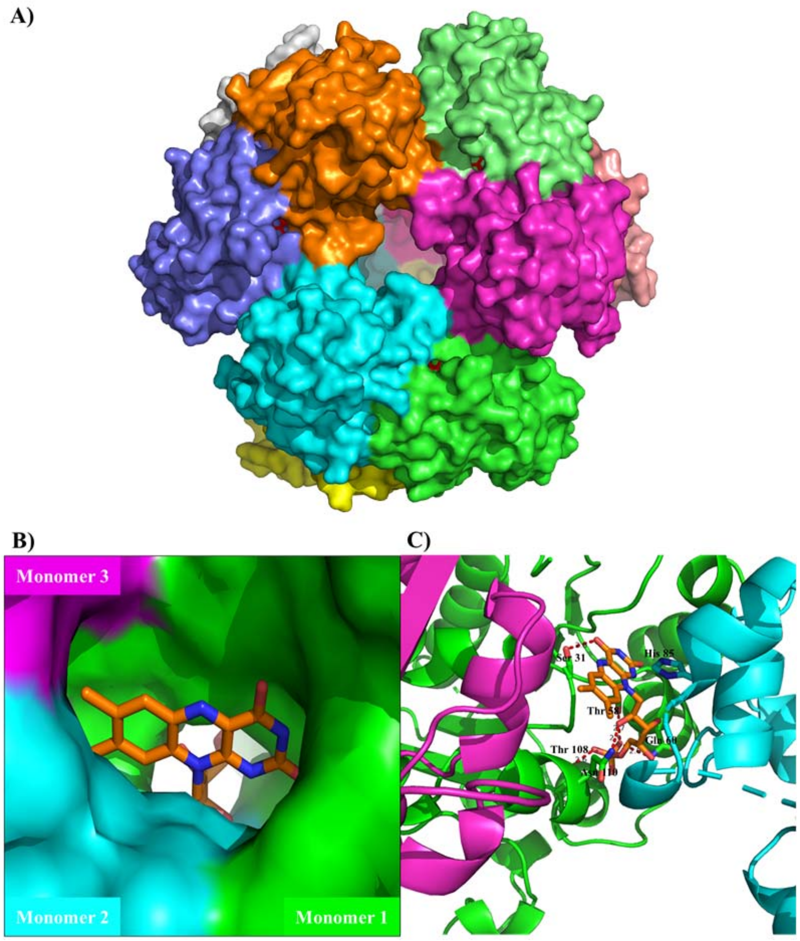


**Figure S9.** Crystal structure of SpaF dodecamer bound with FMN molecules. (A) Overall structure of SpaF dodecamer. (B) Close-up view of FMN bound in SpaF. (C) FMN is positioned in SpaF by a network of hydrogen bonds.

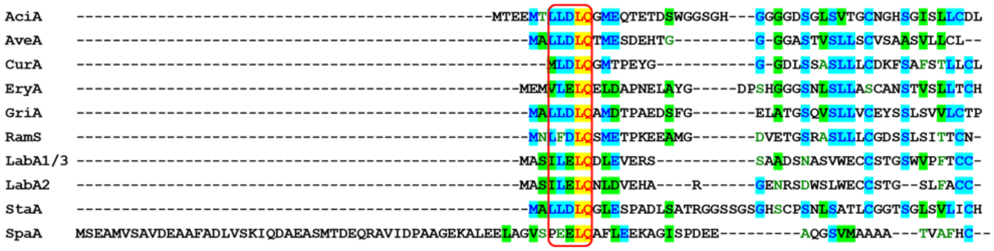


**Figure S10.** Sequence alignment of class III lanthipeptide precursor peptides and SpaA. The accession numbers of class III precursor peptides in GenBank are listed as follows: AciA (ACU73104.1), AveA (WP_010988897.1), CurA (WP_086014640.1), EryA (WP_009949109.1), GriA (WP_003966507.1), RamS (NP_630757.1), LabA1/3 (CAX48972.1), LabA2 (CAX48973.1), StaA (WP_013015818.1), SpaA (WP_104531249.1).

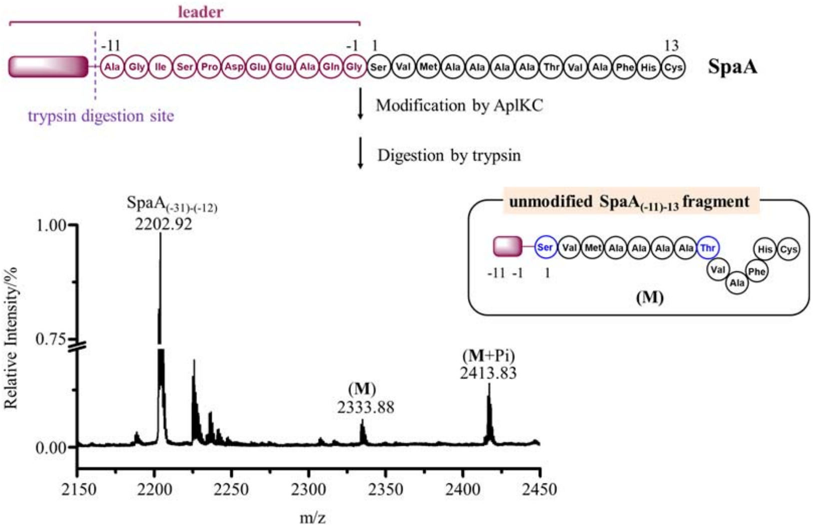


**Figure S11.** SpaA is modified AplKC, the class III lanthipeptide synthetase from of NAI-112 biosynthesis, with one phosphorylation in *E. coli*. The resulting peptide products were purified, digested by trypsin and analyzed by MALDI-TOF MS. Digestion conditions: 20 mM Tris-HCl, pH = 8.0, 1.0 μM trypsin, 100 μM peptides at 37 °C for 6 h.

Monoisotopic mass of SpaA and its derivative:

**(M)**, calc. 2333.05, obs. 2333.88;

**(M+Pi)**, calc. 2413.02, obs. 2413.83;

**SpaA$_{\mathbf{(-31)-(-12)}}$**, calc. 2202.12, obs. 2202.92.

**Pi**: phosphorylation

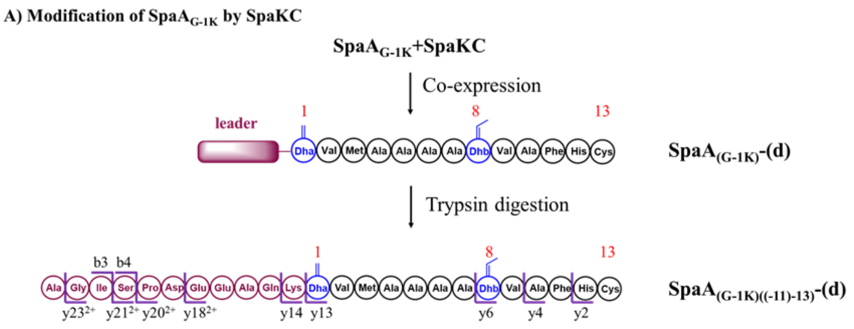


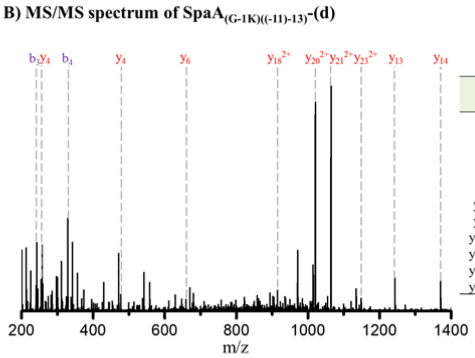


| | Expected Mass m/z | Observed Mass m/z | relative deviation (ppm) |
|---|---|---|---|
| b3 | 242.1500 | 242.1507 | 3.0560 |
| b4 | 329.1820 | 329.1808 | -3.6150 |
| y2 | 259.0860 | 259.0863 | 1.2351 |
| y4 | 477.1915 | 477.1903 | -2.5357 |
| y6 | 659.2971 | 659.2926 | -6.8255 |
| y13 | 1242.5760 | 1242.5770 | 0.8209 |
| y14 | 1370.6710 | 1370.6720 | 0.7660 |
| y18$^{2+}$ | 914.4296 | 914.4263 | -3.5760 |
| y20$^{2+}$ | 1020.4694 | 1020.4713 | 1.8325 |
| y21$^{2+}$ | 1063.9854 | 1063.9853 | -0.1316 |
| y23$^{2+}$ | 1149.0382 | 1149.0357 | -2.1757 |

**Figure S12.** SpaKC dehydrates SpaA$_{G-1K}$ peptide by 2-fold. (A) Modification of SpaA$_{G-1K}$ by SpaKC. (B) LC-MS/MS analysis of **SpaA$_{(G-1K)((-11)-13)}$-(d)** generated from trypsin digestion of **SpaA$_{(G-1K)}$-(d)**. The *b* and *y* ions are listed in the table and indicated in the spectrum.

**A) Modification of SpaA$_{G\text{-}1K}$ by SpaKC-SpaF combination**

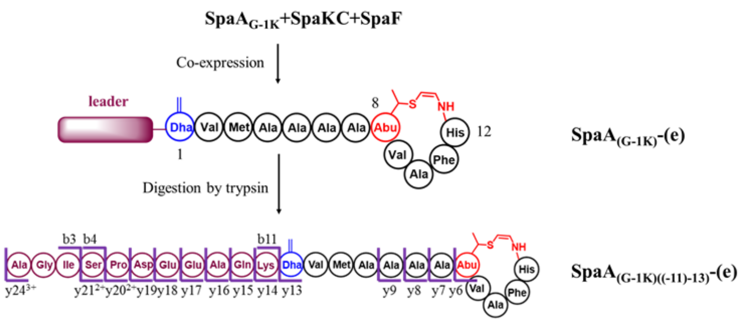


**B) MS/MS spectrum of the SpaA$_{(G\text{-}1K)((-11)\text{-}13)}$-(e)**

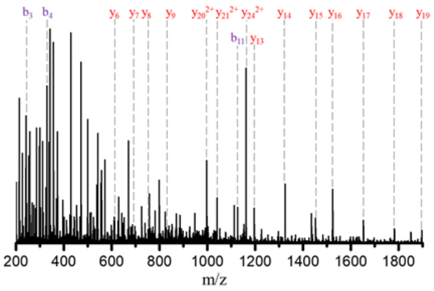


| | Expected Mass m/z | Observed Mass m/z | relative deviation (ppm) |
|---|---|---|---|
| b3 | 242.1500 | 242.1499 | -0.2478 |
| b4 | 329.1820 | 329.1829 | 2.7644 |
| b11 | 1126.5375 | 1126.5402 | 2.3612 |
| y6 | 613.2917 | 613.2936 | 0.3098 |
| y7 | 684.3288 | 684.3253 | -0.5144 |
| y8 | 755.3659 | 755.3703 | 0.5785 |
| y9 | 826.4030 | 826.4080 | 0.6002 |
| y13 | 1196.5706 | 1196.5759 | 0.4446 |
| y14 | 1324.6656 | 1324.6755 | 0.7511 |
| y15 | 1452.7241 | 1452.7291 | 0.3428 |
| y16 | 1523.7612 | 1523.7737 | 0.8177 |
| y17 | 1652.8038 | 1652.8157 | 0.7182 |
| y18 | 1781.8464 | 1781.8591 | 0.7116 |
| y19 | 1896.8734 | 1896.8874 | 0.7396 |
| $y20^{2+}$ | 997.4667 | 997.4715 | 0.4782 |
| $y21^{2+}$ | 1040.9827 | 1040.9911 | 0.8031 |
| $y24^{2+}$ | 1161.5541 | 1161.5589 | 0.4167 |

**Figure S13.** SpaKC-SpaF combination installs an AviMeCys motif in SpaA$_{\mathbf{G\text{-}1K}}$ peptide. (A) Modification of SpaA$_{G\text{-}1K}$ by SpaKC-SpaF combination. (B) MS/MS analysis of **SpaA$_{(G\text{-}1K)((-11)\text{-}13)}$-(e)** generated from trypsin digestion of **SpaA$_{(G\text{-}1K)}$-(e)**. The *b* and *y* ions are listed in the table and indicated in the spectrum.

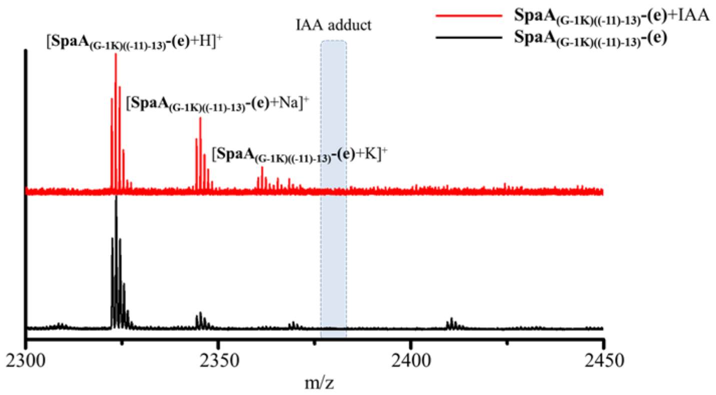


**Figure S14.** MALDI-TOF MS analysis of iodoacetamide (IAA)-treated **SpaA$_{(G-1K)((-11)-13)}$-(e)**. No IAA addition was detected, indicating the absence of free thiols.

Monoisotopic mass of **SpaA$_{(G-1K)((-11)-13)}$-(e)**, calc. 2322.10 Da, obs. 2322.36 Da. The theoretical mass of IAA addition product is 2379.12 Da, which is not observed by MALDI-TOF MS (denoted with the blue box).

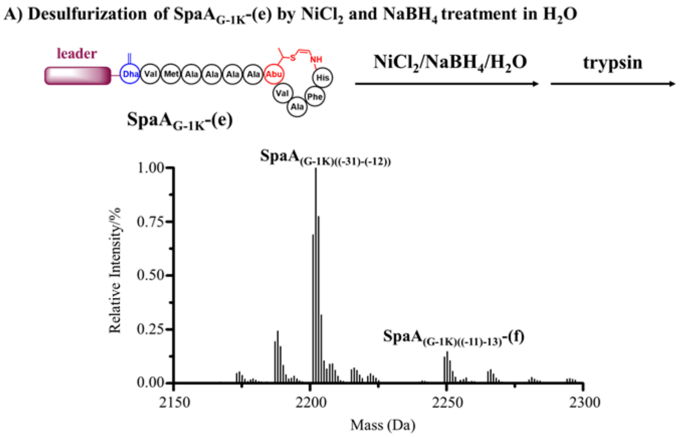


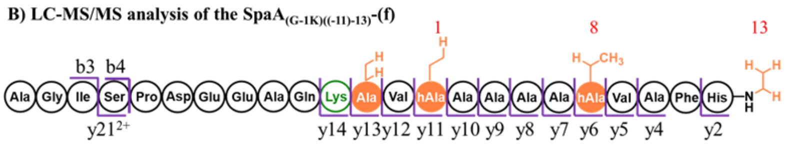


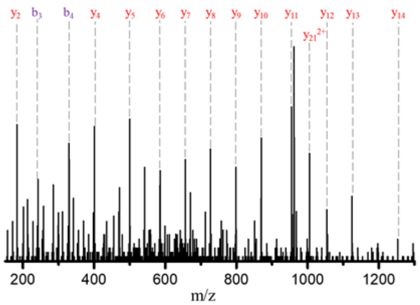


| | Expected Mass m/z | Observed Mass m/z | relative deviation (ppm) |
|---|---|---|---|
| b3 | 242.1500 | 242.1490 | -3.9645 |
| b4 | 329.1820 | 329.1830 | 3.0682 |
| y2 | 183.1241 | 183.1254 | 7.2082 |
| y4 | 401.2296 | 401.2299 | 0.7228 |
| y5 | 500.2980 | 500.2994 | 2.7584 |
| y6 | 585.3508 | 585.3566 | 9.9086 |
| y7 | 656.3879 | 656.3872 | -1.0969 |
| y8 | 727.4250 | 727.4267 | 2.2958 |
| y9 | 798.4621 | 798.4614 | -0.9268 |
| y10 | 869.4993 | 869.5020 | 3.1512 |
| y11 | 954.5520 | 954.5570 | 5.1962 |
| y12 | 1053.6205 | 1053.6232 | 2.6006 |
| y13 | 1124.6576 | 1124.6529 | -4.1613 |
| y14 | 1252.7526 | 1252.7506 | -1.5566 |
| y21$^{2+}$ | 1005.0262 | 1005.0275 | 1.2537 |

**Figure S15.** LC-MS/MS analysis of **SpaA$_{(G-1K)((-11)-13)}$-(f)** generated from reductive desulfurization of **SpaA$_{(G-1K)}$-(e)** by NiCl$_2$ and NaBH$_4$ in H$_2$O. (A) **SpaA$_{(G-1K)}$-(e)** was treated with NaBH$_4$ and NiCl$_2$ and digested by trypsin. The resulting mixture was analyzed by LC-MS. Deconvoluted mass of SpaA$_{G-1K}$ derivative: **SpaA$_{(G-1K)((-11)-13)}$-(f)**, calc. 2249.180 Da, obs. 2249.200 Da. **SpaA$_{(G-1K)((-31)-(-12))}$**, calc. 2201.110 Da, obs. 2201.110 Da. Digestion conditions: 20 mM Tris-HCl, pH = 8.0, 1.0 µM trypsin, 100 µM peptide product at 37 °C for 6 h. (B) MS/MS analysis of **SpaA$_{(G-1K)((-11)-13)}$-(f)**. The *b* and *y* ions are listed in the table and marked in the spectrum.

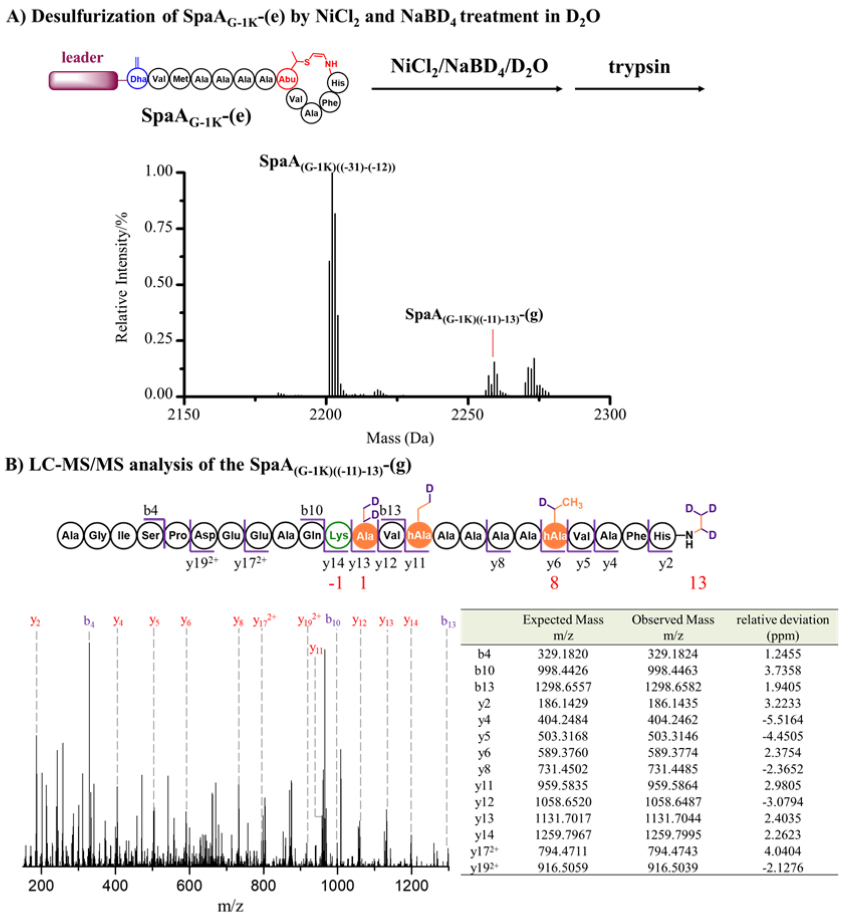


|  | Expected Mass m/z | Observed Mass m/z | relative deviation (ppm) |
|---|---|---|---|
| b4 | 329.1820 | 329.1824 | 1.2455 |
| b10 | 998.4426 | 998.4463 | 3.7358 |
| b13 | 1298.6557 | 1298.6582 | 1.9405 |
| y2 | 186.1429 | 186.1435 | 3.2233 |
| y4 | 404.2484 | 404.2462 | -5.5164 |
| y5 | 503.3168 | 503.3146 | -4.4505 |
| y6 | 589.3760 | 589.3774 | 2.3754 |
| y8 | 731.4502 | 731.4485 | -2.3652 |
| y11 | 959.5835 | 959.5864 | 2.9805 |
| y12 | 1058.6520 | 1058.6487 | -3.0794 |
| y13 | 1131.7017 | 1131.7044 | 2.4035 |
| y14 | 1259.7967 | 1259.7995 | 2.2623 |
| $y17^{2+}$ | 794.4711 | 794.4743 | 4.0404 |
| $y19^{2+}$ | 916.5059 | 916.5039 | -2.1276 |

**Figure S16.** LC-MS/MS analysis of **SpaA$_{(G-1K)((-11)-13)}$-(g)** generated from reductive desulfurization by $NiCl_2$ and $NaBD_4$ in $D_2O$. (A) **SpaA$_{(G-1K)}$-(e)** was treated with $NaBD_4$ and $NiCl_2$ and digested by trypsin. The resulting mixture was analyzed by LC-MS. Deconvoluted mass of SpaA$_{G-1K}$ derivative: **SpaA$_{(G-1K)((-11)-13)}$-(g)**, calc. 2256.224 Da, obs. 2256.229 Da. **SpaA$_{(G-1K)((-31)-(-12))}$**, calc. 2201.110 Da, obs. 2201.120 Da. Digestion conditions: 20 mM Tris-HCl, pH = 8.0, 1.0 µM trypsin, 100 µM peptide product at 37 °C for 6 h. (B) LC-MS/MS analysis of **SpaA$_{(G-1K)((-11)-13)}$-(g)**. The *b* and *y* ions are listed in the table and marked in the spectrum.

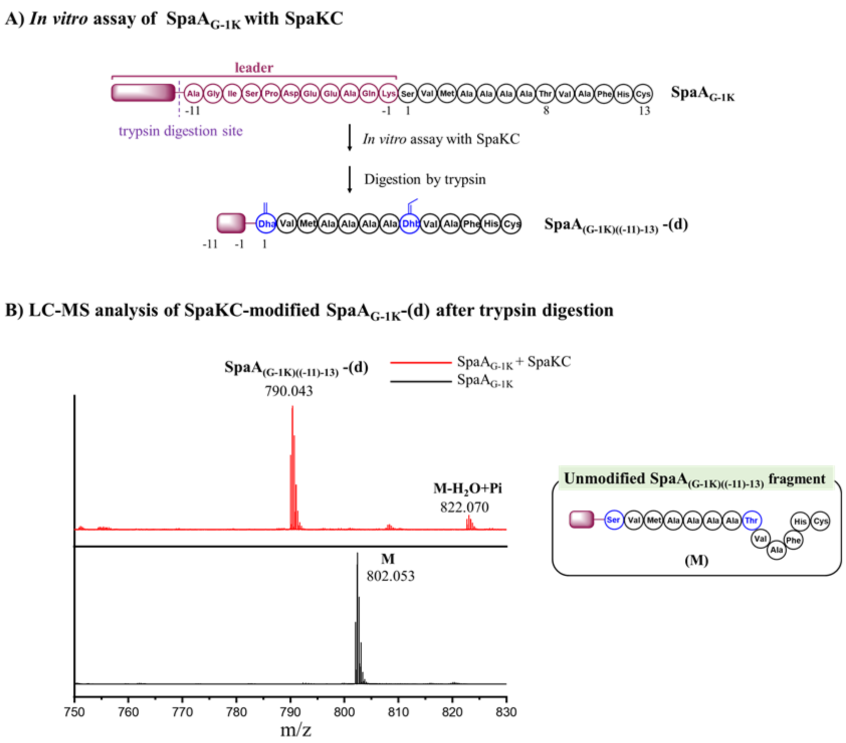


**Figure S17.** SpaKC dehydrates SpaA$_{G-1K}$ by 2-fold *in vitro*. (A). SpaA$_{G-1K}$ peptide is incubated with SpaKC in the presence of 5mM ATP and 5mM $MgCl_2$ and digested by trypsin before subjected to LC-MS analysis. (B) MS spectrum of SpaA$_{G-1K}$ peptide treated with and without SpaKC after trypsin digestion.

Digestion conditions: 20 mM Tris-HCl, pH = 8.0, 1.0 µM trypsin, 100 µM peptide product at 37 °C for 6 h.

Monoisotopic mass of SpaA$_{G-1K}$ and its derivative:

**SpaA$_{(G-1K)((-11)-13)}$-(d)**, calc. 790.040, obs. 790.043;

(**M-$H_2O$+Pi**), calc. 822.699, obs. 822.070.

**(M)**, calc. 802.047, obs. 802.053.

**Pi**: phosphorylation

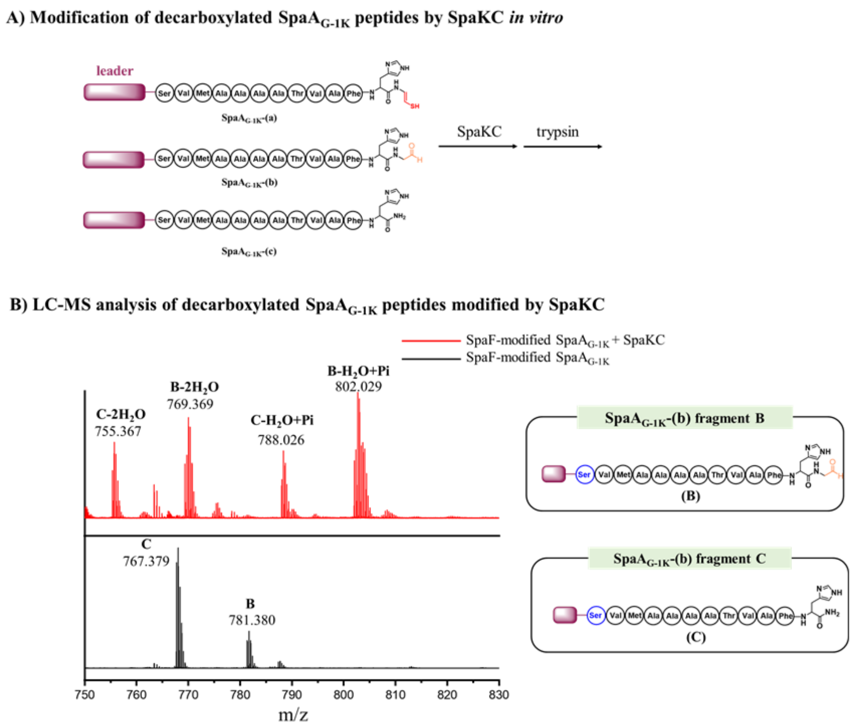


**Figure S18.** SpaKC dehydrates decarboxylated SpaA peptides. (A). Decarboxylated SpaA$_{G\text{-}1K}$ peptides, which are prepared by SpaF modification, are incubated with SpaKC and digested by trypsin before subjected to LC-MS analysis. (B) MS spectrum of decarboxylated SpaA peptides treated with and without SpaKC after trypsin digestion.

Digestion conditions: 20 mM Tris-HCl, pH = 8.0, 1.0 µM trypsin, 100 µM peptide product at 37 °C for 6 h.

Monoisotopic mass of decarboxylated products and dehydrated products:

**(B)**: calc. 781.386, obs. 781.380;

**(C)**: calc. 767.383, obs. 767.379;

**(C-2$H_2$O)**: calc. 755.376, obs. 755.367;

**(B-2$H_2$O)**: calc. 769.379, obs. 769.369;

**(C-$H_2$O+Pi)**: calc. 788.035, obs. 788.026;

**(B-$H_2$O+Pi)**: calc. 802.047, obs. 802.029.

**Pi**: phosphorylation

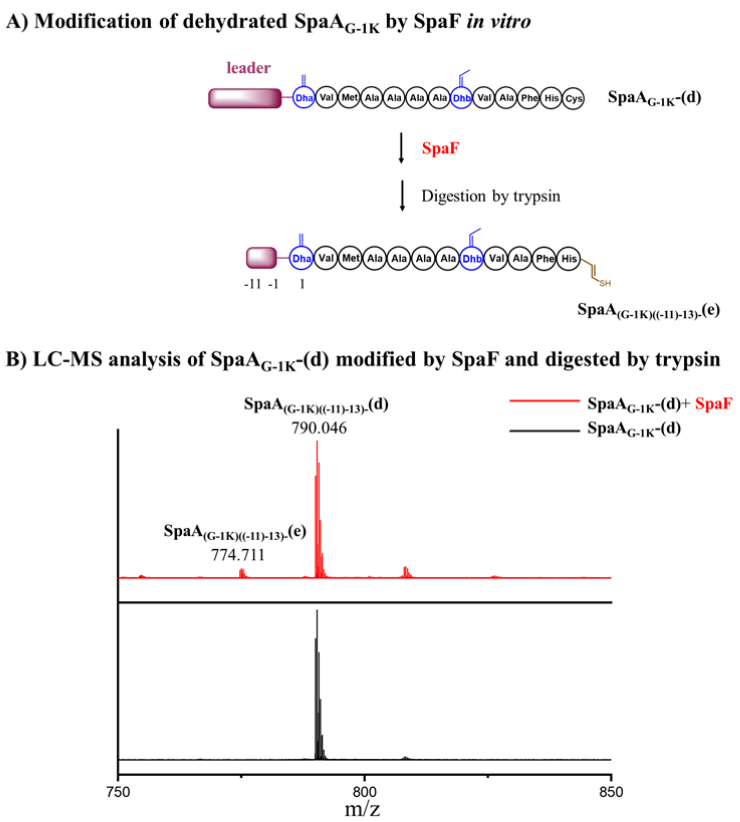


**Figure S19.** Dehydrated peptide **SpaA$_{(G-1K)}$-(d)** was modified by SpaF. (A) **SpaA$_{(G-1K)}$-(d)** was incubated with SpaF *in vitro* and digested by trypsin before LC-MS analysis. (B) MS spectrum of **SpaA$_{(G-1K)}$-(d)** treated with and without SpaF.

Digestion conditions: 20 mM Tris-HCl, pH = 8.0, 1.0 µM trypsin, 100 µM peptide product at 37 °C for 6 h.

**SpaA$_{(G-1K)((-11)-13)}$-(d):** calc. 790.040, obs. 790.036.

**SpaA$_{(G-1K)((-11)-13)}$-(e):** calc. 774.705, obs. 774.711.

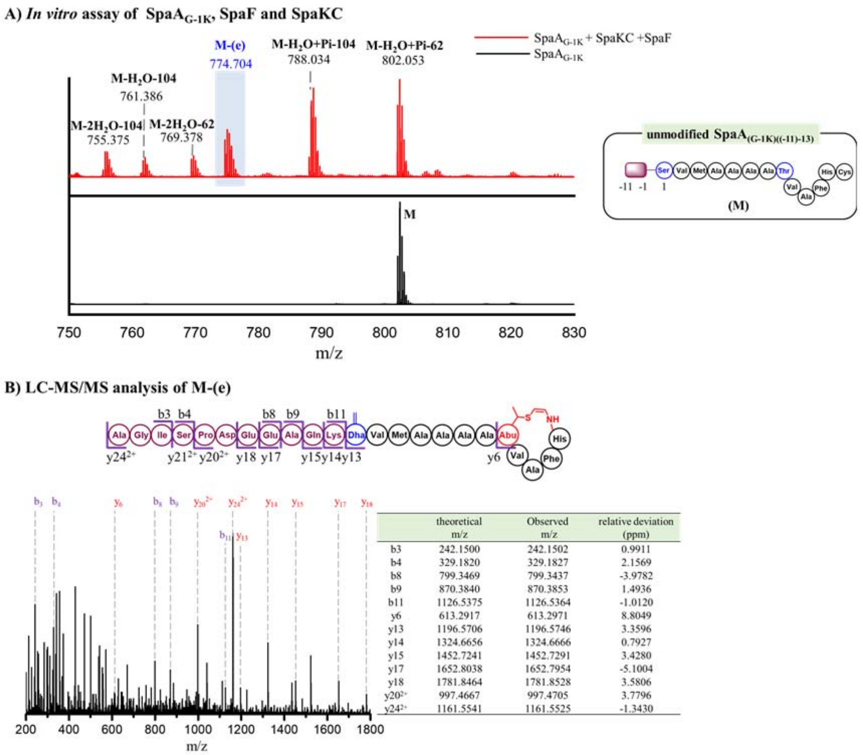


| | theoretical m/z | Observed m/z | relative deviation (ppm) |
|---|---|---|---|
| b3 | 242.1500 | 242.1502 | 0.9911 |
| b4 | 329.1820 | 329.1827 | 2.1569 |
| b8 | 799.3469 | 799.3437 | -3.9782 |
| b9 | 870.3840 | 870.3853 | 1.4936 |
| b11 | 1126.5375 | 1126.5364 | -1.0120 |
| y6 | 613.2917 | 613.2971 | 8.8049 |
| y13 | 1196.5706 | 1196.5746 | 3.3596 |
| y14 | 1324.6656 | 1324.6666 | 0.7927 |
| y15 | 1452.7241 | 1452.7291 | 3.4280 |
| y17 | 1652.8038 | 1652.7954 | -5.1004 |
| y18 | 1781.8464 | 1781.8528 | 3.5806 |
| $y20^{2+}$ | 997.4667 | 997.4705 | 3.7796 |
| $y24^{2+}$ | 1161.5541 | 1161.5525 | -1.3430 |

**Figure S20.** Cyclization of $SpaA_{G\text{-}1K}$ by SpaKC-SpaF combination *in vitro*. (A) $SpaA_{G\text{-}1K}$ was modified by SpaKC-SpaF combination *in vitro*, digested by trypsin and subjected to LC-MS analysis. Digestion conditions: 20 mM Tris-HCl, pH = 8.0, 1 µM trypsin, 100 µM peptide product at 37 °C for 6 h. (B) LC-MS/MS analysis of the segment **(M-(e))** generated from trypsin digestion. Results show that **(M-(e))** is identical to **$SpaA_{(G\text{-}1K)((-11)\text{-}13)}$-(e)** as a cyclized peptide with a C-terminal AviMeCys motif. The *b* and *y* ions are listed in the table and marked in the spectrum.

Monoisotopic mass of $SpaA_{G\text{-}1K}$ and its derivative:

**(M-2$H_2O$-104)**: calc. 755.376, obs. 755.375;

**(M-$H_2O$-104)**: calc. 761.379, obs. 761.386;

**(M-2$H_2O$-62)**: calc. 769.379, obs. 769.378;

**(M-(e)):** calc. 774.705, obs. 774.704;

**(M-$H_2O$+Pi-104)**: calc. 788.035, obs. 788.034;

**(M-$H_2O$+Pi-62)**: calc. 802.047, obs. 802.053.

**(M)**: calc. 802.047, obs. 802.053.

**Pi**: phosphorylation

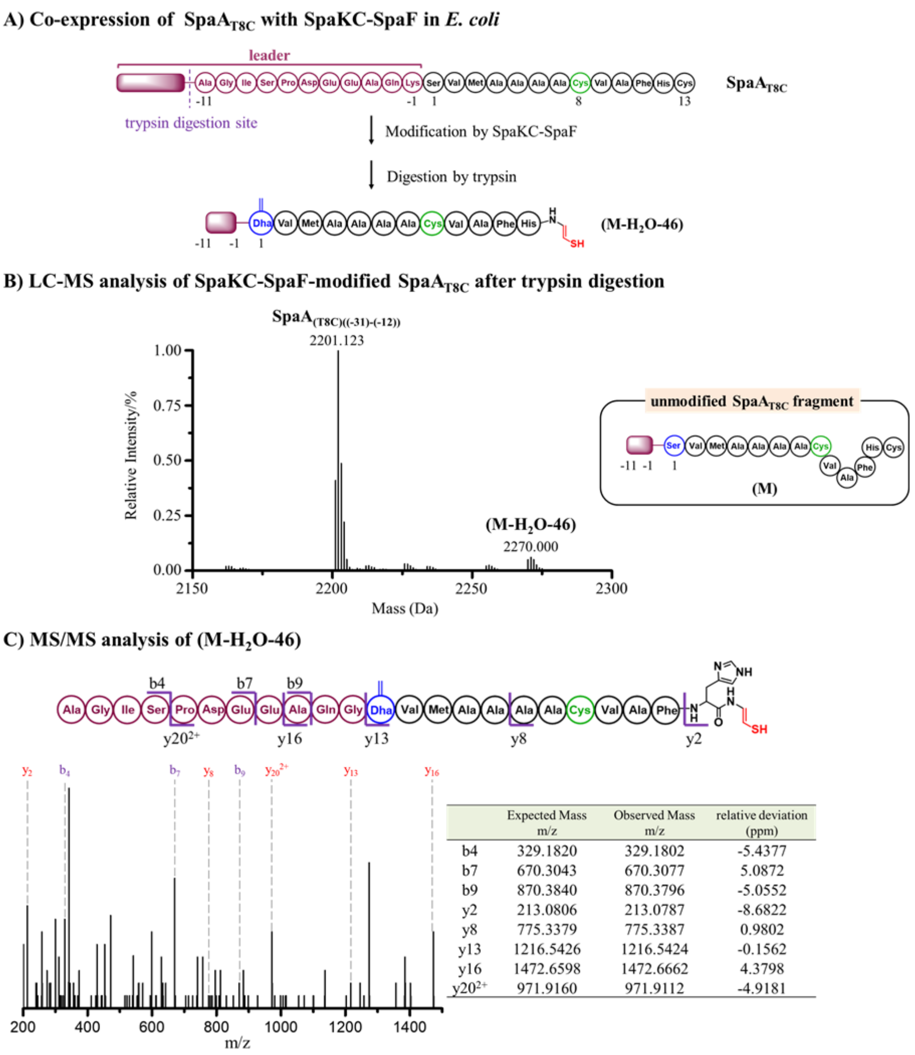


| | Expected Mass m/z | Observed Mass m/z | relative deviation (ppm) |
|---|---|---|---|
| b4 | 329.1820 | 329.1802 | -5.4377 |
| b7 | 670.3043 | 670.3077 | 5.0872 |
| b9 | 870.3840 | 870.3796 | -5.0552 |
| y2 | 213.0806 | 213.0787 | -8.6822 |
| y8 | 775.3379 | 775.3387 | 0.9802 |
| y13 | 1216.5426 | 1216.5424 | -0.1562 |
| y16 | 1472.6598 | 1472.6662 | 4.3798 |
| y20$^{2+}$ | 971.9160 | 971.9112 | -4.9181 |

**Figure S21.** Modification of SpaA$_{T8C}$ peptide by SpaKC-SpaF combination. (A) His$_6$-tagged SpaA$_{T8C}$ was co-expressed with SpaF and SpaKC in *E. coli*, purified and digested by trypsin before subjected to LC-MS analysis. Digestion conditions: 20 mM Tris-HCl, pH = 8.0, 1.0 µM trypsin, 100 µM peptide product at 37 °C for 6 h. (B) MS spectrum of modified SpaA$_{T8C}$ after trypsin digestion. (C) MS/MS spectrum of **(M-H$_2$O-46)**. The *b* and *y* ions are listed in the table and marked in the spectrum.

Monoisotopic mass of SpaA$_{T8C}$ derivatives:

**(M-H$_2$O-46)**, calc. 2269.991 Da, obs. 2270.000 Da.

**SpaA$_{(T8C)((-31)-(-12))}$**, calc. 2201.110, obs. 2201.123.

**-46 Da**: mass loss by oxidative decarboxylation of Cys

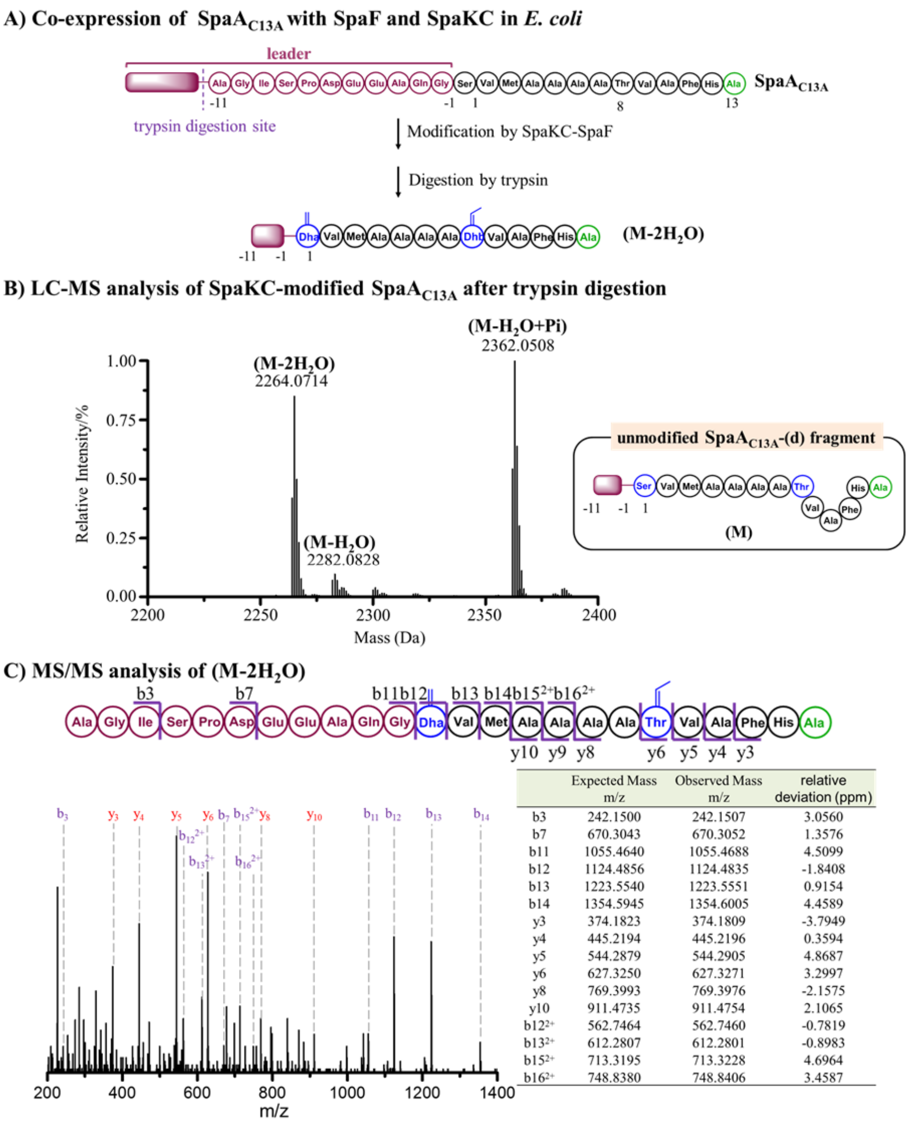


| | Expected Mass m/z | Observed Mass m/z | relative deviation (ppm) |
|---|---|---|---|
| b3 | 242.1500 | 242.1507 | 3.0560 |
| b7 | 670.3043 | 670.3052 | 1.3576 |
| b11 | 1055.4640 | 1055.4688 | 4.5099 |
| b12 | 1124.4856 | 1124.4835 | -1.8408 |
| b13 | 1223.5540 | 1223.5551 | 0.9154 |
| b14 | 1354.5945 | 1354.6005 | 4.4589 |
| y3 | 374.1823 | 374.1809 | -3.7949 |
| y4 | 445.2194 | 445.2196 | 0.3594 |
| y5 | 544.2879 | 544.2905 | 4.8687 |
| y6 | 627.3250 | 627.3271 | 3.2997 |
| y8 | 769.3993 | 769.3976 | -2.1575 |
| y10 | 911.4735 | 911.4754 | 2.1065 |
| $b12^{2+}$ | 562.7464 | 562.7460 | -0.7819 |
| $b13^{2+}$ | 612.2807 | 612.2801 | -0.8983 |
| $b15^{2+}$ | 713.3195 | 713.3228 | 4.6964 |
| $b16^{2+}$ | 748.8380 | 748.8406 | 3.4587 |

**Figure S22.** Modification of SpaA$_{C13A}$ by SpaKC-SpaF. (A) His$_6$-tagged SpaA$_{C13A}$ was co-expressed with SpaF and SpaKC in *E. coli*, purified and digested by trypsin before subjected to LC-MS analysis. Digestion conditions: 20 mM Tris-HCl, pH = 8.0, 1 µM trypsin, 100 µM peptide product at 37 °C for 6 h. (B) MS spectrum of modified SpaA$_{C13A}$ after trypsin digestion. (C) MS/MS spectrum of **(M-2H$_2$O)**. The *b* and *y* ions are listed in the table and marked in the spectrum.

Monoisotopic mass of SpaA$_{C13A}$ derivative:

**(M-2H$_2$O)**, calc. 2264.053 Da, obs. 2264.071 Da;

**(M-H$_2$O)**, calc. 2282.064 Da, obs. 2282.083 Da;

**(M-H$_2$O+Pi)**, calc. 2362.030 Da, obs. 2362.051 Da.

**Pi**: phosphorylation

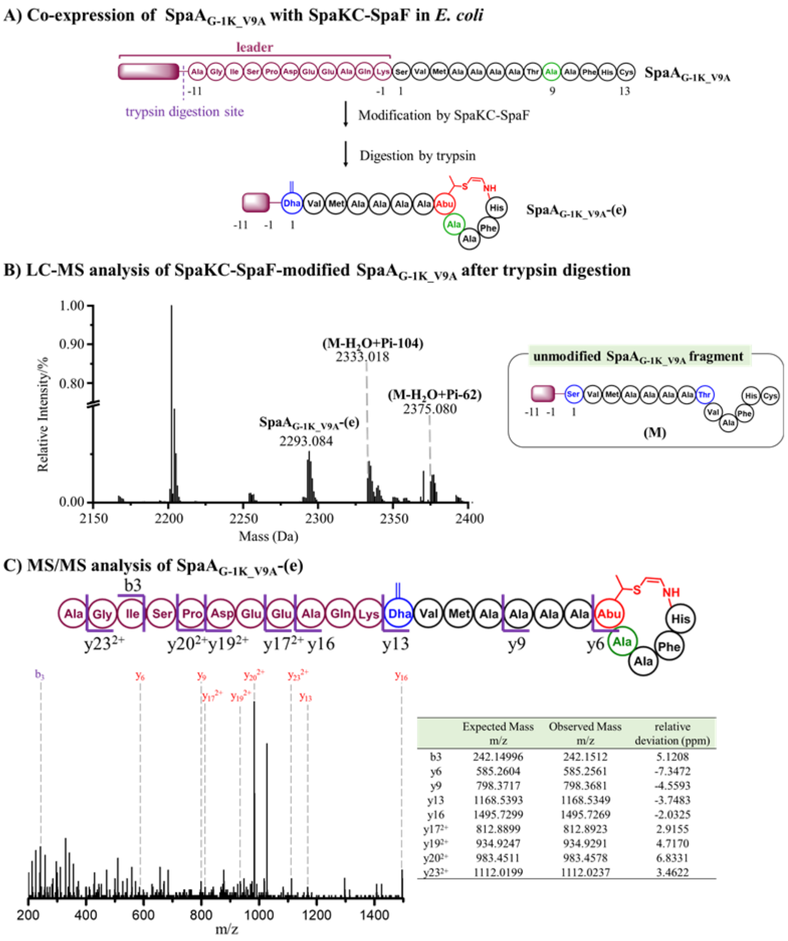


| | Expected Mass m/z | Observed Mass m/z | relative deviation (ppm) |
|---|---|---|---|
| b3 | 242.14996 | 242.1512 | 5.1208 |
| y6 | 585.2604 | 585.2561 | -7.3472 |
| y9 | 798.3717 | 798.3681 | -4.5593 |
| y13 | 1168.5393 | 1168.5349 | -3.7483 |
| y16 | 1495.7299 | 1495.7269 | -2.0325 |
| $y17^{2+}$ | 812.8899 | 812.8923 | 2.9155 |
| $y19^{2+}$ | 934.9247 | 934.9291 | 4.7170 |
| $y20^{2+}$ | 983.4511 | 983.4578 | 6.8331 |
| $y23^{2+}$ | 1112.0199 | 1112.0237 | 3.4622 |

**Figure S23.** Modification of SpaA$_{G\text{-}1K_V9A}$ peptide by SpaKC-SpaF combination. (A) His$_6$-tagged SpaA$_{G\text{-}1K_V9A}$ was co-expressed with SpaF and SpaKC in *E. coli*, and then digested by trypsin. The digestion mixture was subjected to LC-MS analysis. Digestion conditions: 20 mM Tris-HCl, pH = 8.0, 1 µM trypsin, 100 µM peptide product at 37 °C for 6 h. (B) MS spectrum of modified SpaA$_{G\text{-}1K_V9A}$ after trypsin digestion. (C) MS/MS spectrum of **SpaA$_{G\text{-}1K_V9A}$-(e)**. The *b* and *y* ions are listed in the table and marked in the spectrum.

Monoisotopic mass of SpaA$_{G\text{-}1K_V9A}$ derivatives:

**SpaA$_{G\text{-}1K_V9A}$-(e)**, calc. 2293.061 Da, obs. 2293.084 Da;

**(M-$H_2O$+Pi-104)**, calc. 2333.051 Da, obs. 2333.018 Da;

**(M-$H_2O$+Pi-62)**, calc. 2375.062 Da, obs. 2375.080 Da.

**-62** & **-104**: hydrolysis of decarboxylated Cys.

**Pi**: phosphorylation

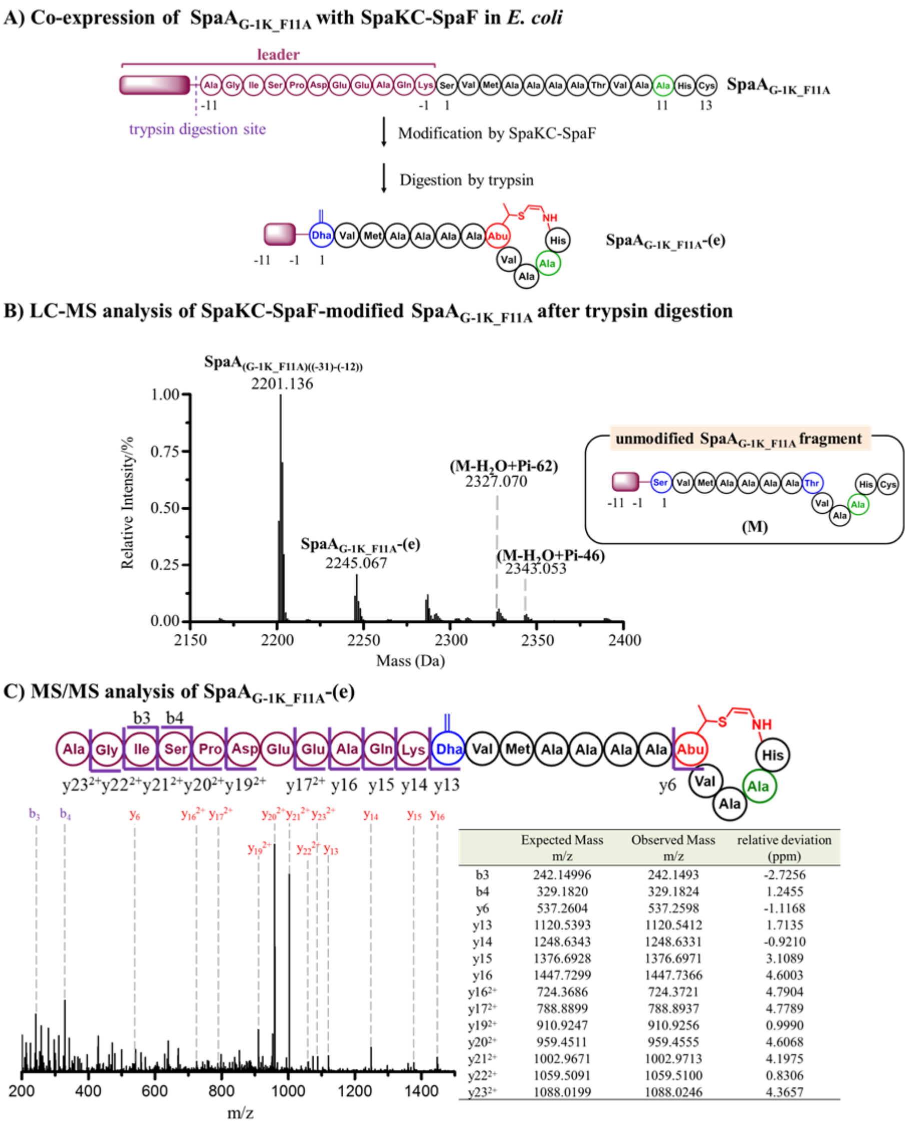


| | Expected Mass m/z | Observed Mass m/z | relative deviation (ppm) |
|---|---|---|---|
| b3 | 242.14996 | 242.1493 | -2.7256 |
| b4 | 329.1820 | 329.1824 | 1.2455 |
| y6 | 537.2604 | 537.2598 | -1.1168 |
| y13 | 1120.5393 | 1120.5412 | 1.7135 |
| y14 | 1248.6343 | 1248.6331 | -0.9210 |
| y15 | 1376.6928 | 1376.6971 | 3.1089 |
| y16 | 1447.7299 | 1447.7366 | 4.6003 |
| $y16^{2+}$ | 724.3686 | 724.3721 | 4.7904 |
| $y17^{2+}$ | 788.8899 | 788.8937 | 4.7789 |
| $y19^{2+}$ | 910.9247 | 910.9256 | 0.9990 |
| $y20^{2+}$ | 959.4511 | 959.4555 | 4.6068 |
| $y21^{2+}$ | 1002.9671 | 1002.9713 | 4.1975 |
| $y22^{2+}$ | 1059.5091 | 1059.5100 | 0.8306 |
| $y23^{2+}$ | 1088.0199 | 1088.0246 | 4.3657 |

**Figure S24.** Modification of SpaA$_{G-1K_F11A}$ peptide by SpaKC-SpaF combination. (A) His$_6$-tagged SpaA$_{G-1K_F11A}$ was co-expressed with SpaF and SpaKC in *E. coli*, and then digested by trypsin. The digestion mixture was subjected to LC-MS analysis. Digestion conditions: 20 mM Tris-HCl, pH = 8.0, 1 µM trypsin, 100 µM peptide product at 37 °C for 6 h. (B) MS spectrum of modified SpaA$_{G-1K_F11A}$ after trypsin digestion. (C) MS/MS spectrum of **SpaA$_{G-1K_F11A}$-(e)**. The b and y ions are listed in the table and marked in the spectrum.

Monoisotopic mass of SpaA$_{G-1K_F11A}$ derivatives:

**SpaA$_{G-1K_F11A}$-(e)**, calc. 2245.062 Da, obs. 2245.067 Da;

**(M-$H_2O$+Pi-62)**, calc. 2327.062 Da, obs. 2327.070 Da;

**(M-$H_2O$+Pi-46)**, calc. 2343.039 Da, obs. 2343.053 Da.

**SpaA$_{(G-1K_F11A)((-31)-(-12))}$**, calc. 2201.110, obs. 2201.136.

**-46**: mass loss by oxidative decarboxylation of Cys.

**-62**: hydrolysis of the decarboxylated Cys.

**Pi**: phosphorylation

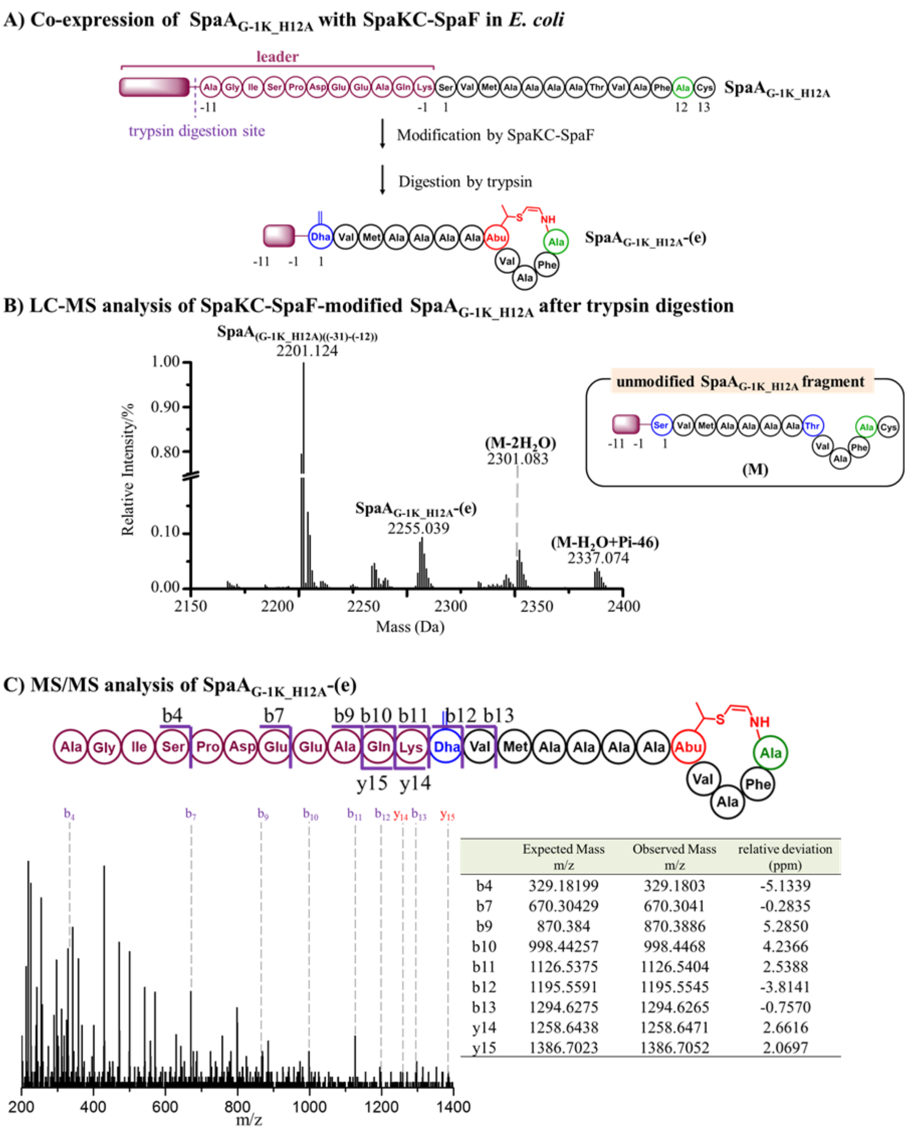


| | Expected Mass m/z | Observed Mass m/z | relative deviation (ppm) |
|---|---|---|---|
| b4 | 329.18199 | 329.1803 | -5.1339 |
| b7 | 670.30429 | 670.3041 | -0.2835 |
| b9 | 870.384 | 870.3886 | 5.2850 |
| b10 | 998.44257 | 998.4468 | 4.2366 |
| b11 | 1126.5375 | 1126.5404 | 2.5388 |
| b12 | 1195.5591 | 1195.5545 | -3.8141 |
| b13 | 1294.6275 | 1294.6265 | -0.7570 |
| y14 | 1258.6438 | 1258.6471 | 2.6616 |
| y15 | 1386.7023 | 1386.7052 | 2.0697 |

**Figure S25.** Modification of SpaA$_{G\text{-}1K_H12A}$ peptide by SpaKC-SpaF combination. (A) His$_6$-tagged SpaA$_{G\text{-}1K_H12A}$ was co-expressed with SpaF and SpaKC in *E. coli*, and then digested by trypsin. The digestion mixture was subjected to LC-MS analysis. Digestion conditions: 20 mM Tris-HCl, pH = 8.0, 1 µM trypsin, 100 µM peptide product at 37 °C for 6 h. (B) MS spectrum of modified SpaA$_{G\text{-}1K_H12A}$ after trypsin digestion. (C) MS/MS spectrum of **SpaA$_{G\text{-}1K_H12A}$-(e)**. The *b* and *y* ions are listed in the table and marked in the spectrum.

Monoisotopic mass of SpaA$_{G\text{-}1K_H12A}$ derivatives:

**SpaA$_{G\text{-}1K_H12A}$-(e)**, calc. 2255.071 Da, obs. 2255.039 Da;

**(M-2H$_2$O)**, calc. 2301.077 Da, obs. 2301.083 Da;

**(M-H$_2$O+Pi-46)**, calc. 2337.071 Da, obs. 2337.074 Da.

**SpaA$_{(G\text{-}1K_H12A)((-31)\text{-}(-12))}$**, calc. 2201.110, obs. 2201.124.

**-46**: mass loss by oxidative decarboxylation of Cys.

**Pi**: phosphorylation

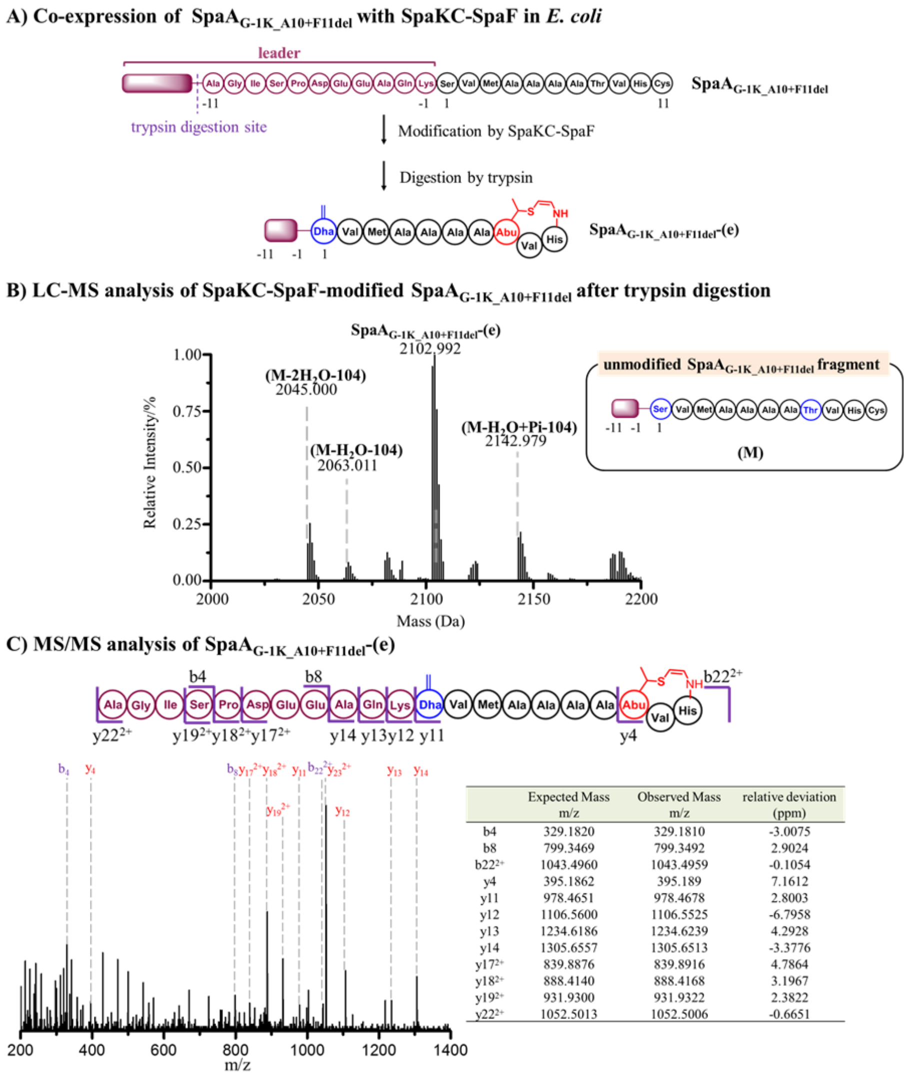


|  | Expected Mass m/z | Observed Mass m/z | relative deviation (ppm) |
|---|---|---|---|
| b4 | 329.1820 | 329.1810 | -3.0075 |
| b8 | 799.3469 | 799.3492 | 2.9024 |
| $b22^{2+}$ | 1043.4960 | 1043.4959 | -0.1054 |
| y4 | 395.1862 | 395.189 | 7.1612 |
| y11 | 978.4651 | 978.4678 | 2.8003 |
| y12 | 1106.5600 | 1106.5525 | -6.7958 |
| y13 | 1234.6186 | 1234.6239 | 4.2928 |
| y14 | 1305.6557 | 1305.6513 | -3.3776 |
| $y17^{2+}$ | 839.8876 | 839.8916 | 4.7864 |
| $y18^{2+}$ | 888.4140 | 888.4168 | 3.1967 |
| $y19^{2+}$ | 931.9300 | 931.9322 | 2.3822 |
| $y22^{2+}$ | 1052.5013 | 1052.5006 | -0.6651 |

**Figure S26.** Modification of SpaA$_{G\text{-}1K_A10+F11del}$ peptide by SpaKC-SpaF combination. (A) His$_6$-tagged SpaA$_{G\text{-}1K_A10+F11del}$ was co-expressed with SpaF and SpaKC in *E. coli*, and then digested by trypsin. The digestion mixture was subjected to LC-MS analysis. Digestion conditions: 20 mM Tris-HCl, pH = 8.0, 1.0 µM trypsin, 100 µM peptide product at 37 °C for 6 h. (B) MS spectrum of modified SpaA$_{G\text{-}1K_A10+F11del}$ after trypsin digestion. (C) MS/MS spectrum of **SpaA$_{G\text{-}1K_A10+F11del}$ -(e)**. The *b* and *y* ions are listed in the table and marked in the spectrum.

Monoisotopic mass of SpaA$_{G\text{-}1K_A10+F11del}$ derivatives:

**SpaA$_{G\text{-}1K_A10+F11del}$-(e)**, calc. 2250.056 Da, obs. 2250.668 Da;

**(M-2$H_2O$-104)**, calc. 2045.000 Da, obs. 2045.000 Da;

**(M-$H_2O$-104)**, calc. 2063.011 Da, obs. 2063.011 Da;

**(M-$H_2O$+Pi-104)**, calc. 2142.977 Da, obs. 2142.979 Da.

**-104**: hydrolysis of decarboxylated Cys.

**Pi**: phosphorylation

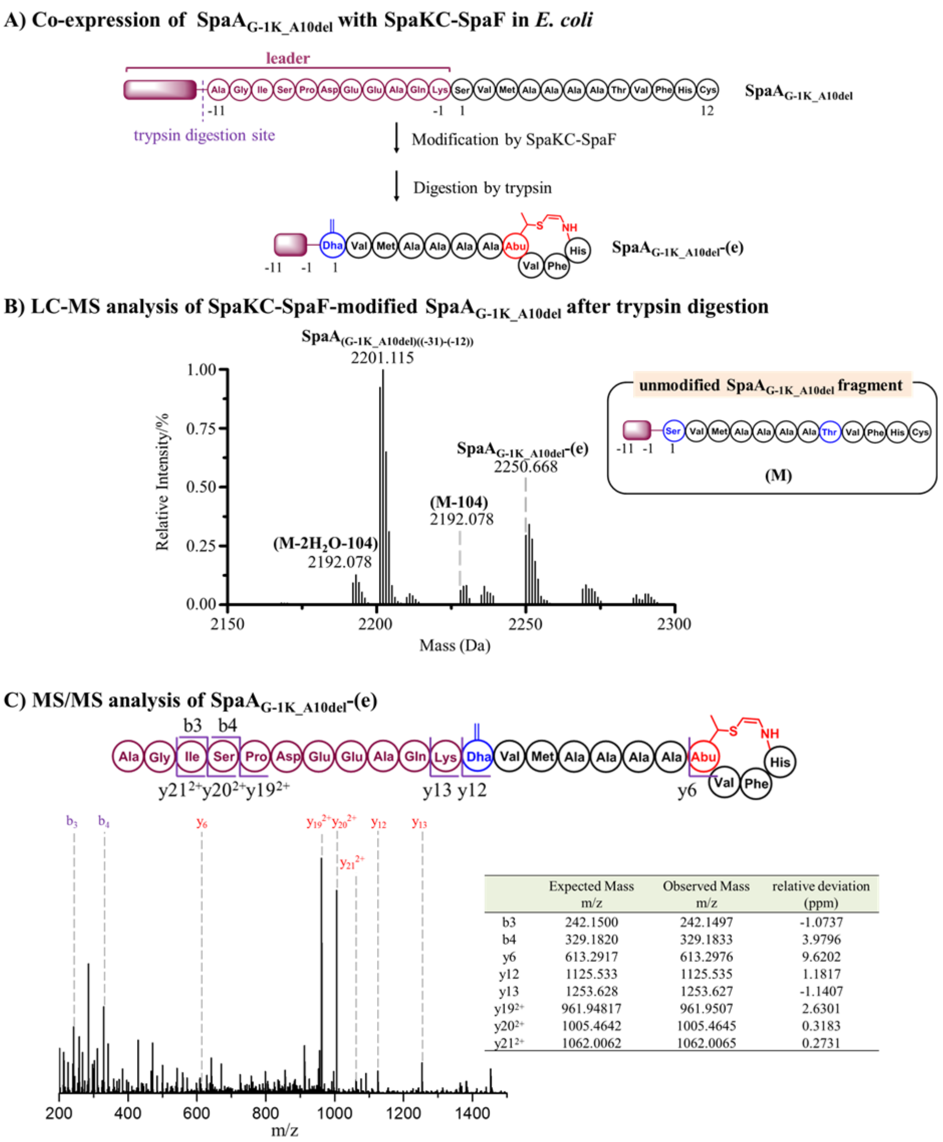


| | Expected Mass m/z | Observed Mass m/z | relative deviation (ppm) |
|---|---|---|---|
| b3 | 242.1500 | 242.1497 | -1.0737 |
| b4 | 329.1820 | 329.1833 | 3.9796 |
| y6 | 613.2917 | 613.2976 | 9.6202 |
| y12 | 1125.533 | 1125.535 | 1.1817 |
| y13 | 1253.628 | 1253.627 | -1.1407 |
| y19$^{2+}$ | 961.94817 | 961.9507 | 2.6301 |
| y20$^{2+}$ | 1005.4642 | 1005.4645 | 0.3183 |
| y21$^{2+}$ | 1062.0062 | 1062.0065 | 0.2731 |

**Figure S27.** Modification of SpaA$_{G-1K_A10del}$ peptide by SpaKC-SpaF combination. (A) $His_6$-tagged SpaA$_{G-1K_A10del}$ was co-expressed with SpaF and SpaKC in *E. coli*, and then digested by trypsin. The digestion mixture was subjected to LC-MS analysis. Digestion conditions: 20 mM Tris-HCl, pH = 8.0, 1.0 µM trypsin, 100 µM peptide product at 37 °C for 6 h. (B) MS spectrum of modified SpaA$_{G-1K_A10del}$ after trypsin digestion. (C) MS/MS spectrum of **SpaA$_{G-1K_A10del}$-(e)**. The *b* and *y* ions are listed in the table and marked in the spectrum.

Monoisotopic mass of SpaA$_{G-1K_A10del}$ derivatives:

**SpaA$_{G-1K_A10del}$-(e)**, calc. 2250.056 Da, obs. 2250.668 Da;

**(M-2$H_2$O-104)**, calc. 2192.069 Da, obs. 2192.078 Da;

**(M-104)**, calc. 2228.290 Da, obs. 2228.078 Da.

**SpaA$_{(G-1K_A10del)((-31)-(-12))}$**, calc. 2201.110, obs. 2201.115.

**-104**: hydrolysis of decarboxylated Cys.

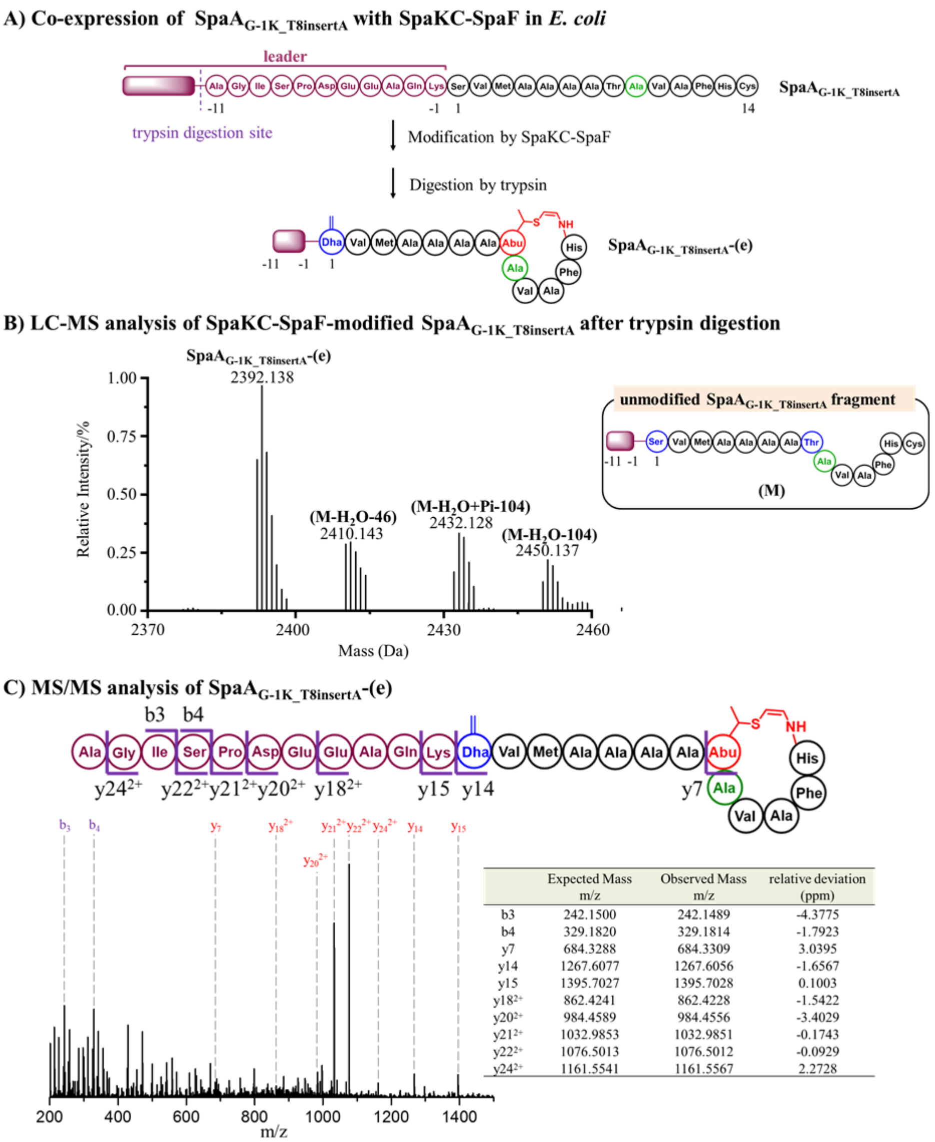


| | Expected Mass m/z | Observed Mass m/z | relative deviation (ppm) |
|---|---|---|---|
| b3 | 242.1500 | 242.1489 | -4.3775 |
| b4 | 329.1820 | 329.1814 | -1.7923 |
| y7 | 684.3288 | 684.3309 | 3.0395 |
| y14 | 1267.6077 | 1267.6056 | -1.6567 |
| y15 | 1395.7027 | 1395.7028 | 0.1003 |
| $y18^{2+}$ | 862.4241 | 862.4228 | -1.5422 |
| $y20^{2+}$ | 984.4589 | 984.4556 | -3.4029 |
| $y21^{2+}$ | 1032.9853 | 1032.9851 | -0.1743 |
| $y22^{2+}$ | 1076.5013 | 1076.5012 | -0.0929 |
| $y24^{2+}$ | 1161.5541 | 1161.5567 | 2.2728 |

**Figure S28.** Modification of $\mathrm{SpaA}_{\mathrm{G\text{-}1K\_T8insertA}}$ peptide by SpaKC-SpaF combination. (A) $\mathrm{His}_6$-tagged $\mathrm{SpaA}_{\mathrm{G\text{-}1K\_T8insertA}}$ was co-expressed with SpaF and SpaKC in *E. coli*, and then digested by trypsin. The digestion mixture was subjected to LC-MS analysis. Digestion conditions: 20 mM Tris-HCl, pH = 8.0, 1.0 μM trypsin, 100 μM peptide product at 37 °C for 6 h. (B) MS spectrum of modified $\mathrm{SpaA}_{\mathrm{G\text{-}1K\_V9A}}$ after trypsin digestion. (C) MS/MS spectrum of **$\mathbf{SpaA}_{\mathbf{G\text{-}1K\_T8insertA}}$-(e)**. The *b* and *y* ions are listed in the table and marked in the spectrum.

Monoisotopic mass of $\mathrm{SpaA}_{\mathrm{G\text{-}1K\_T8insertA}}$ derivatives:

**$\mathbf{SpaA}_{\mathbf{G\text{-}1K\_T8insertA}}$-(e)**, calc. 2392.131 Da, obs. 2392.138 Da;

**(M-$H_2O$-46)**, calc. 2410.141 Da, obs. 2410.143 Da;

**(M-$H_2O$+Pi-104)**, calc. 2432.120 Da, obs. 2432.128 Da;

**(M-$H_2O$-104)**, calc. 2450.130 Da, obs. 2450.137 Da.

**-46**: mass loss by oxidative decarboxylation of Cys.

**-104**: hydrolysis of decarboxylated Cys.

**Pi**: phosphorylation

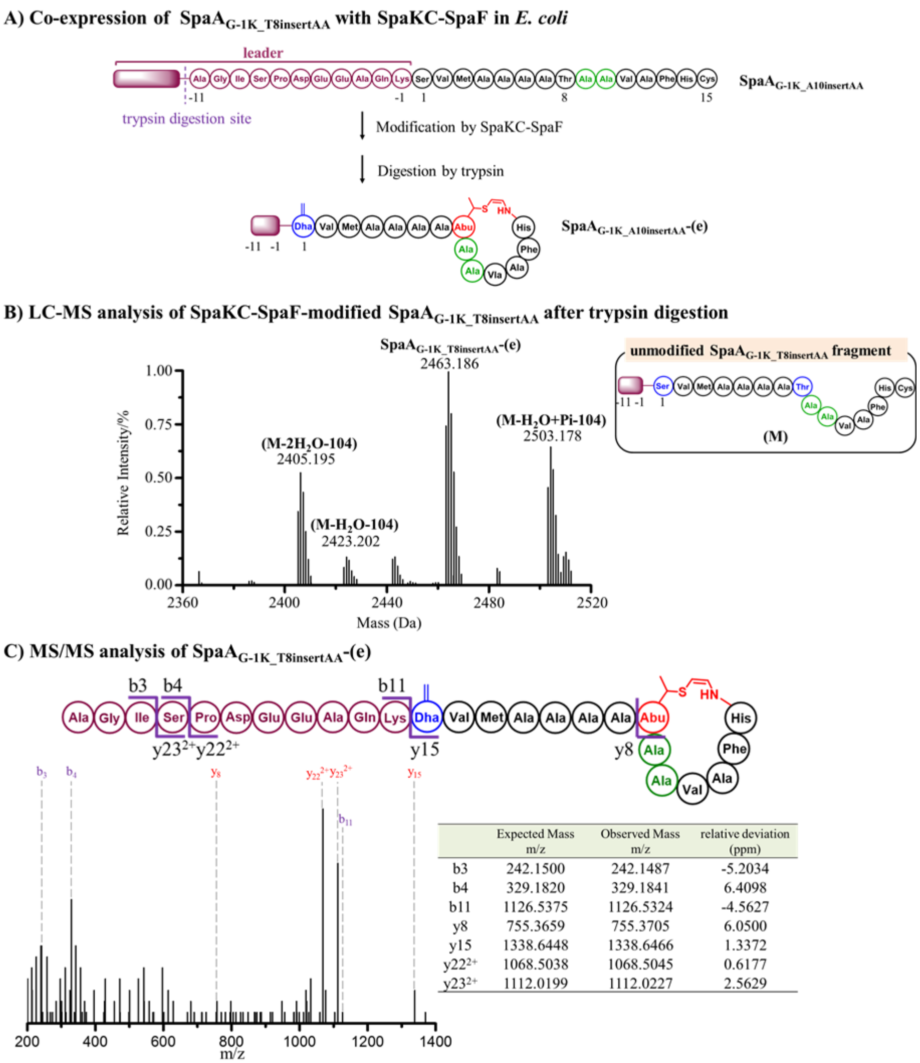


| | Expected Mass m/z | Observed Mass m/z | relative deviation (ppm) |
|---|---|---|---|
| b3 | 242.1500 | 242.1487 | -5.2034 |
| b4 | 329.1820 | 329.1841 | 6.4098 |
| b11 | 1126.5375 | 1126.5324 | -4.5627 |
| y8 | 755.3659 | 755.3705 | 6.0500 |
| y15 | 1338.6448 | 1338.6466 | 1.3372 |
| y22$^{2+}$ | 1068.5038 | 1068.5045 | 0.6177 |
| y23$^{2+}$ | 1112.0199 | 1112.0227 | 2.5629 |

**Figure S29.** Modification of SpaA$_{G\text{-}1K_T8insertAA}$ peptide by SpaKC-SpaF combination. (A) His$_6$-tagged SpaA$_{G\text{-}1K_T8insertAA}$ was co-expressed with SpaF and SpaKC in *E. coli*, and then digested by trypsin. The digestion mixture was subjected to LC-MS analysis. Digestion conditions: 20 mM Tris-HCl, pH = 8.0, 1.0 μM trypsin, 100 μM peptide product at 37 °C for 6 h. (B) MS spectrum of modified SpaA$_{G\text{-}1K_T8insertAA}$ after trypsin digestion. (C) MS/MS spectrum of **SpaA$_{G\text{-}1K_T8insertAA}$-(e)**. The *b* and *y* ions are listed in the table and marked in the spectrum.

Monoisotopic mass of SpaA$_{G\text{-}1K_T8insertAA}$ derivatives:

**SpaA$_{G\text{-}1K_T8insertAA}$-(e)**, calc. 2463.168 Da, obs. 2463.186 Da;

**(M-2H$_2$O-104)**, calc. 2405.180 Da, obs. 2405.195 Da;

**(M-H$_2$O-104)**, calc. 2423.190 Da, obs. 2423.202 Da;

**(M-H$_2$O+Pi-104)**, calc. 2503.157 Da, obs. 2503.178 Da.

**-104**: hydrolysis of decarboxylated Cys.

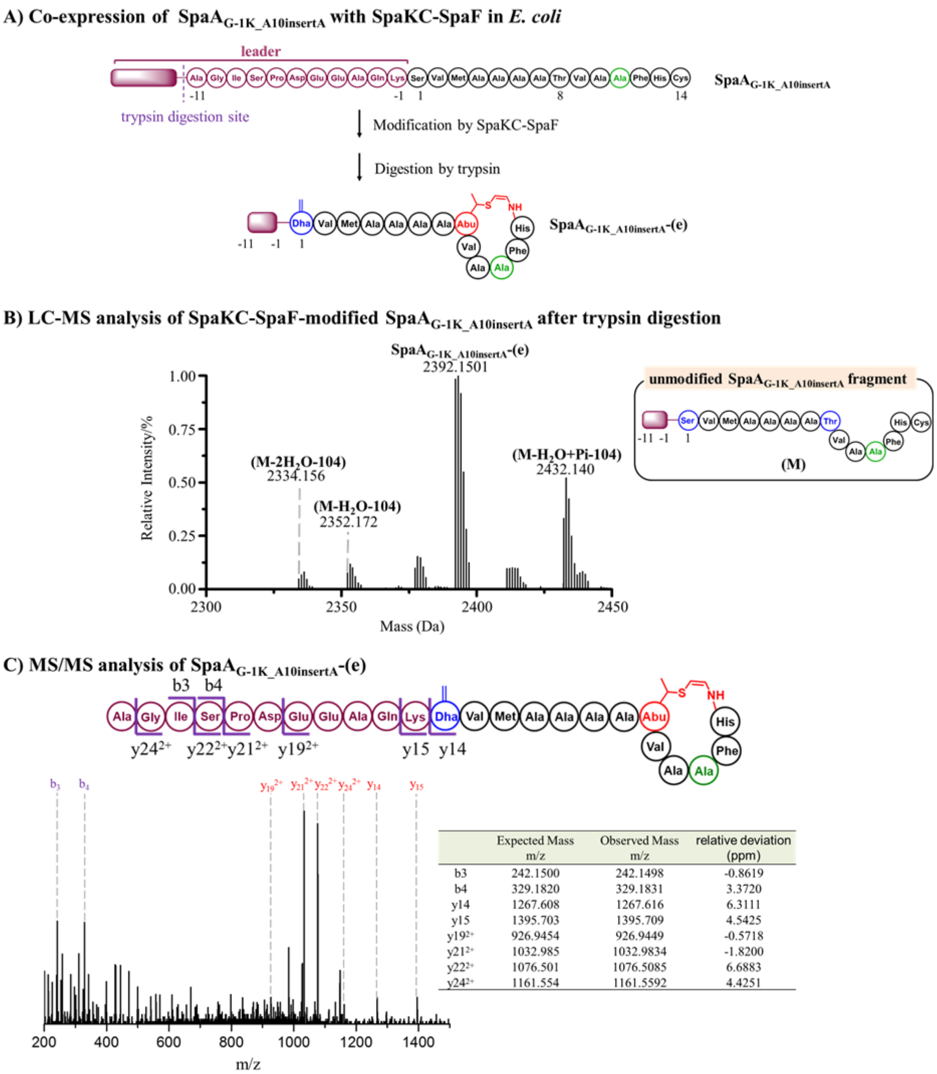


| | Expected Mass m/z | Observed Mass m/z | relative deviation (ppm) |
|---|---|---|---|
| b3 | 242.1500 | 242.1498 | -0.8619 |
| b4 | 329.1820 | 329.1831 | 3.3720 |
| y14 | 1267.608 | 1267.616 | 6.3111 |
| y15 | 1395.703 | 1395.709 | 4.5425 |
| $y19^{2+}$ | 926.9454 | 926.9449 | -0.5718 |
| $y21^{2+}$ | 1032.985 | 1032.9834 | -1.8200 |
| $y22^{2+}$ | 1076.501 | 1076.5085 | 6.6883 |
| $y24^{2+}$ | 1161.554 | 1161.5592 | 4.4251 |

**Figure S30.** Modification of SpaA$_{G\text{-}1K_A10insertA}$ peptide by SpaKC-SpaF combination. (A) His$_6$-tagged SpaA$_{G\text{-}1K_A10insertA}$ was co-expressed with SpaF and SpaKC in *E. coli*, and then digested by trypsin. The digestion mixture was subjected to LC-MS analysis. Digestion conditions: 20 mM Tris-HCl, pH = 8.0, 1 µM trypsin, 100 µM peptide product at 37 °C for 6 h. (B) LC-MS analysis of the segment of co-expression after trypsin digestion. (C) MS/MS spectrum of **SpaA$_{G\text{-}1K_A10insertA}$-(e)**. The *b* and *y* ions are listed in the table and marked in the spectrum.

Monoisotopic mass of SpaA$_{G\text{-}1K_A10insertA}$ derivatives:

**SpaA$_{G\text{-}1K_A10insertA}$-(e)**, calc. 2392.131 Da, obs. 2392.151 Da.

**(M-2$H_2O$-104)**, calc. 2334.143 Da, obs. 2334.156 Da;

**(M-$H_2O$-104)**, calc. 2352.153 Da, obs. 2352.172 Da;

**(M-$H_2O$+Pi-104)**, calc. 2432.120 Da, obs. 2432.140 Da.

**Pi**: phosphorylation

**-104**: hydrolysis of the decarboxylated Cys.

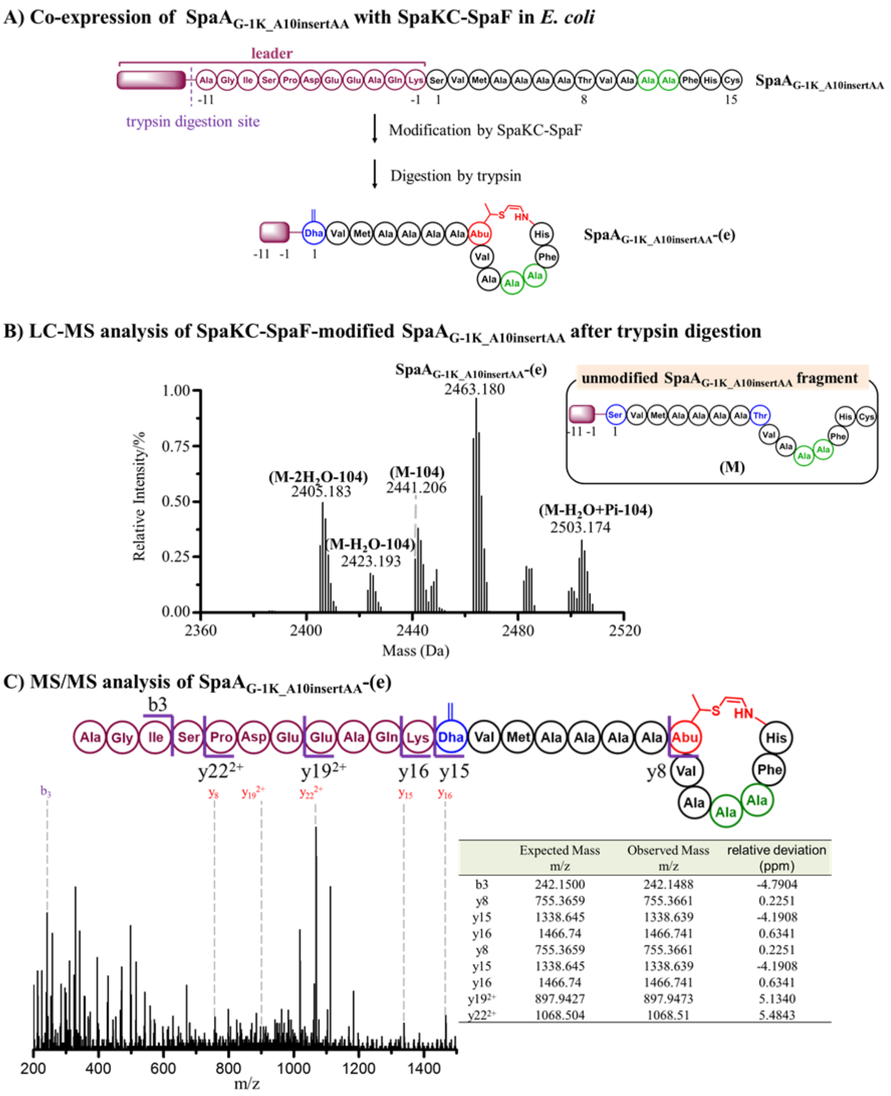


| | Expected Mass m/z | Observed Mass m/z | relative deviation (ppm) |
|---|---|---|---|
| b3 | 242.1500 | 242.1488 | -4.7904 |
| y8 | 755.3659 | 755.3661 | 0.2251 |
| y15 | 1338.645 | 1338.639 | -4.1908 |
| y16 | 1466.74 | 1466.741 | 0.6341 |
| y8 | 755.3659 | 755.3661 | 0.2251 |
| y15 | 1338.645 | 1338.639 | -4.1908 |
| y16 | 1466.74 | 1466.741 | 0.6341 |
| y19$^{2+}$ | 897.9427 | 897.9473 | 5.1340 |
| y22$^{2+}$ | 1068.504 | 1068.51 | 5.4843 |

**Figure S31.** Modification of SpaA$_{G\text{-}1K_A10insertAA}$ peptide by SpaKC-SpaF combination. (A) His$_6$-tagged SpaA$_{G\text{-}1K_A10insertAA}$ was co-expressed with SpaF and SpaKC in *E. coli*, and then digested by trypsin. The digestion mixture was subjected to LC-MS analysis. Digestion conditions: 20 mM Tris-HCl, pH = 8.0, 1.0 µM trypsin, 100 µM peptide product at 37 °C for 6 h. (B) MS spectrum of modified SpaA$_{G\text{-}1K_A10insertAA}$ after trypsin digestion. (C) MS/MS spectrum of **SpaA$_{G\text{-}1K_A10insertAA}$-(e)**. The *b* and *y* ions are listed in the table and marked in the spectrum.

Monoisotopic mass of SpaA$_{G\text{-}1K_A10insertAA}$ derivatives:

**SpaA$_{G\text{-}1K_A10insertAA}$-(e)**, calc. 2463.168 Da, obs. 2463.108 Da.

**(M-2H$_2$O-104)**, calc. 2405.180 Da, obs. 2405.183 Da;

**(M-H$_2$O-104)**, calc. 2423.190 Da, obs. 2423.193 Da;

**(M-104)**, calc. 2441.201 Da, obs. 2441.206 Da;

**(M-H$_2$O+Pi-104)**, calc. 2503.157 Da, obs. 2503.173 Da.

**Pi**: phosphorylation

**-104**: hydrolysis of the decarboxylated Cys.

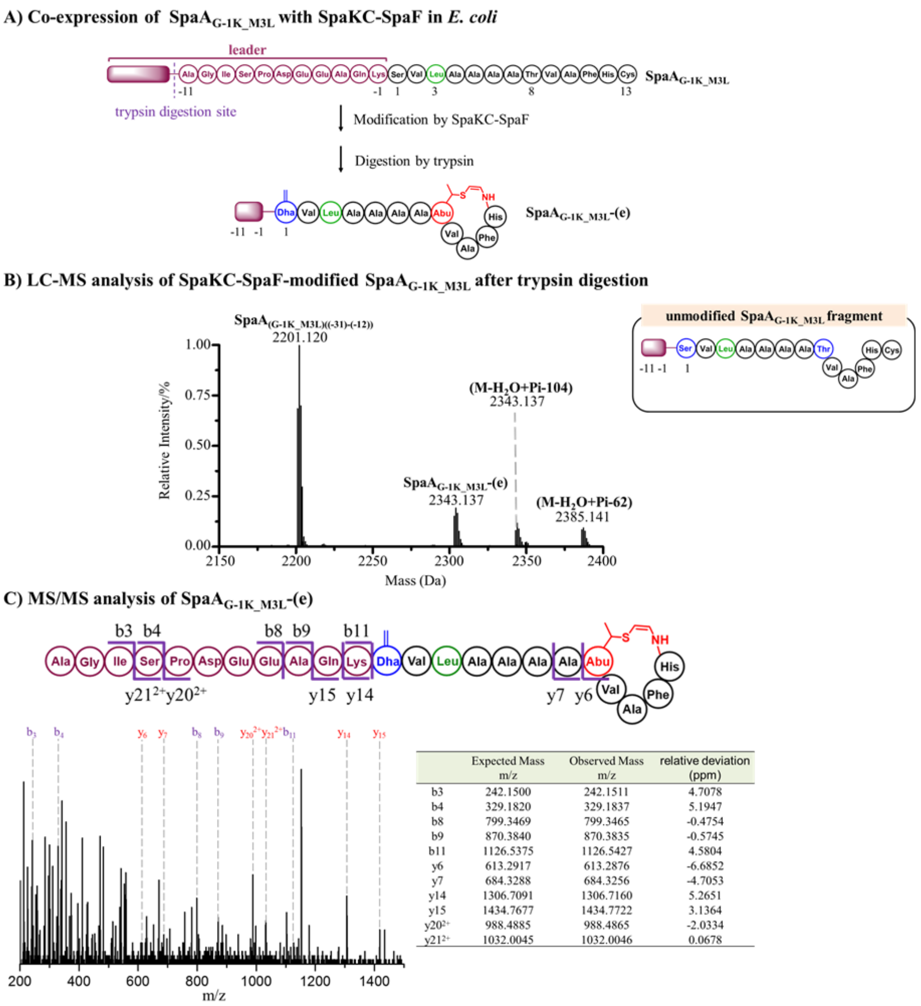


| | Expected Mass m/z | Observed Mass m/z | relative deviation (ppm) |
|---|---|---|---|
| b3 | 242.1500 | 242.1511 | 4.7078 |
| b4 | 329.1820 | 329.1837 | 5.1947 |
| b8 | 799.3469 | 799.3465 | -0.4754 |
| b9 | 870.3840 | 870.3835 | -0.5745 |
| b11 | 1126.5375 | 1126.5427 | 4.5804 |
| y6 | 613.2917 | 613.2876 | -6.6852 |
| y7 | 684.3288 | 684.3256 | -4.7053 |
| y14 | 1306.7091 | 1306.7160 | 5.2651 |
| y15 | 1434.7677 | 1434.7722 | 3.1364 |
| $y20^{2+}$ | 988.4885 | 988.4865 | -2.0334 |
| $y21^{2+}$ | 1032.0045 | 1032.0046 | 0.0678 |

**Figure S32.** Modification of SpaA$_{G\text{-}1K_M3L}$ peptide by SpaKC-SpaF combination. (A) His$_6$-tagged SpaA$_{G\text{-}1K_M3L}$ was co-expressed with SpaF and SpaKC in *E. coli*. The product was digested by trypsin and subjected to LC-MS analysis. Digestion conditions: 20 mM Tris-HCl, pH = 8.0, 1.0 µM trypsin, 100 µM peptide product at 37 °C for 6 h. (B) MS spectrum of modified SpaA$_{G\text{-}1K_M3L}$ after trypsin digestion. (C) MS/MS spectrum of **SpaA$_{G\text{-}1K_M3L}$-(e)**. The *b* and *y* ions are listed in the table and marked in the spectrum.

Monoisotopic mass of SpaA$_{G\text{-}1K_M3L}$ derivatives:

**SpaA$_{G\text{-}1K_M3L}$-(e)**, calc. 2303.137 Da, obs. 2303.149 Da;

**(M-$H_2O$+Pi-104)**, calc. 2343.126 Da, obs. 2343.137;

**(M-$H_2O$+Pi-62)**, calc. 2385.137 Da, obs. 2385.141.

**Pi**: phosphorylation

**-62 & -104**: hydrolysis of decarboxylated Cys.

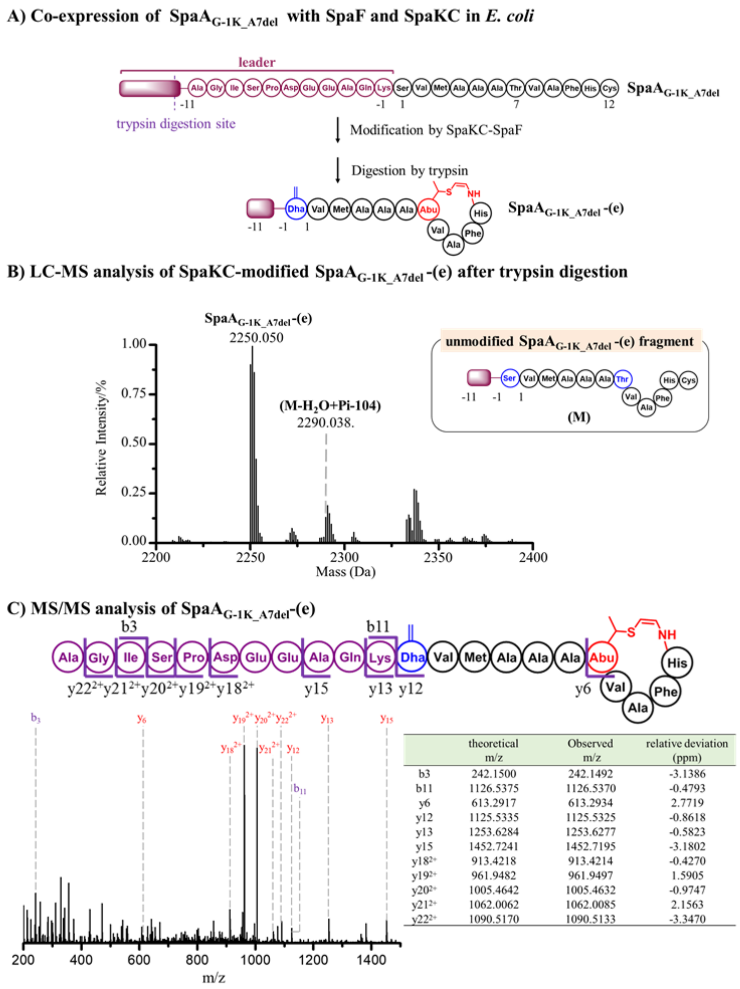


|  | theoretical m/z | Observed m/z | relative deviation (ppm) |
|---|---|---|---|
| b3 | 242.1500 | 242.1492 | -3.1386 |
| b11 | 1126.5375 | 1126.5370 | -0.4793 |
| y6 | 613.2917 | 613.2934 | 2.7719 |
| y12 | 1125.5335 | 1125.5325 | -0.8618 |
| y13 | 1253.6284 | 1253.6277 | -0.5823 |
| y15 | 1452.7241 | 1452.7195 | -3.1802 |
| $y18^{2+}$ | 913.4218 | 913.4214 | -0.4270 |
| $y19^{2+}$ | 961.9482 | 961.9497 | 1.5905 |
| $y20^{2+}$ | 1005.4642 | 1005.4632 | -0.9747 |
| $y21^{2+}$ | 1062.0062 | 1062.0085 | 2.1563 |
| $y22^{2+}$ | 1090.5170 | 1090.5133 | -3.3470 |

**Figure S33.** Modification of SpaA$_{G-1K_A7del}$ peptide by SpaKC-SpaF combination. (A) His$_6$-tagged SpaA$_{G-1K_A7del}$ was co-expressed with SpaF and SpaKC in *E. coli*, purified and then digested by trypsin. The digestion mixture was subjected to LC-MS analysis. Digestion conditions: 20 mM Tris-HCl, pH = 8.0, 1.0 μM trypsin, 100 μM peptide product at 37 °C for 6 h. (B) MS spectrum of modified SpaA$_{G-1K_A7del}$ after trypsin digestion. (C) MS/MS spectrum of **SpaA$_{G-1K_A7del}$-(e)**. The *b* and *y* ions are listed in the table and marked in the spectrum.

Monoisotopic mass of SpaA$_{G-1K_A7del}$ derivatives:

**SpaA$_{G-1K_A7del}$-(e)**, calc. 2250.052 Da, obs. 2250.050 Da;

**(M-$H_2O$+Pi-104)**, calc. 2290.045 Da, obs. 2290.038 Da.

**-104**: hydrolysis of decarboxylated Cys.

**Pi**: phosphorylation

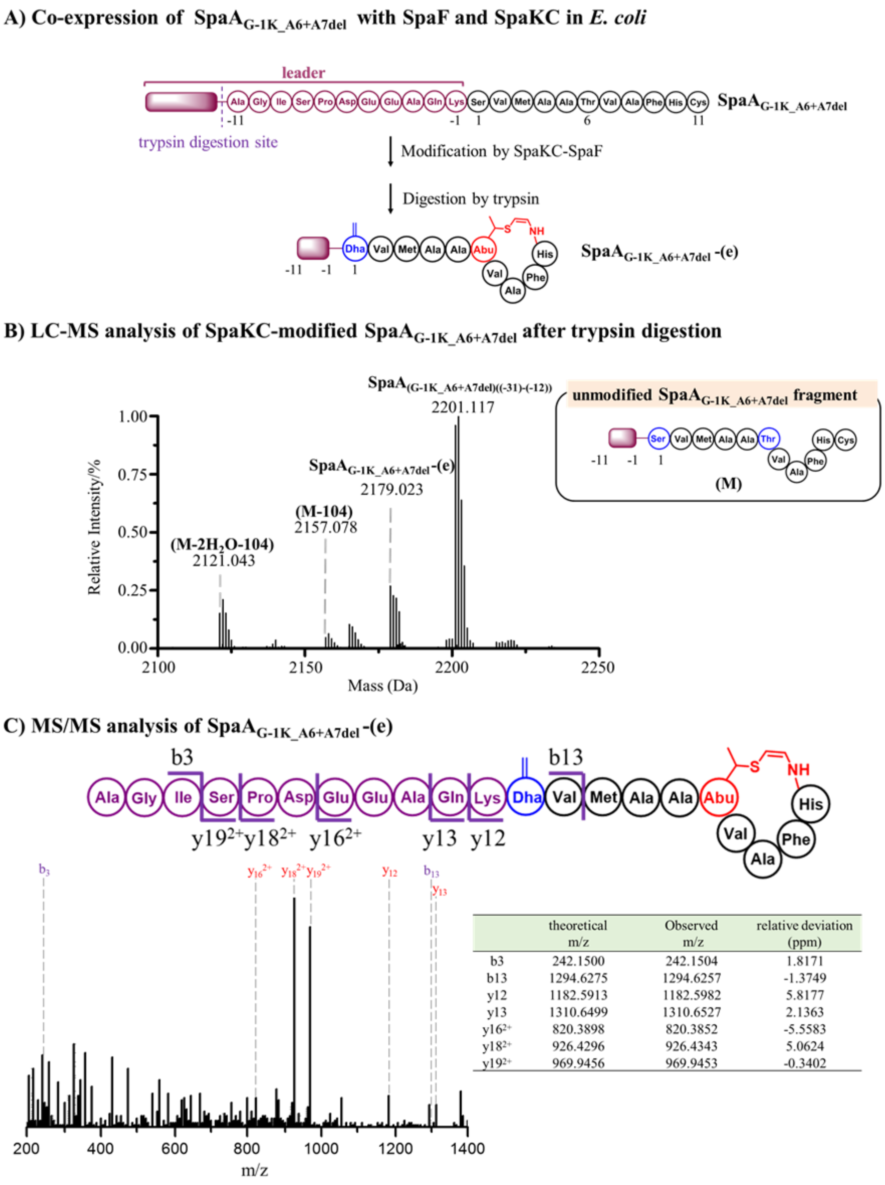


|  | theoretical m/z | Observed m/z | relative deviation (ppm) |
|---|---|---|---|
| b3 | 242.1500 | 242.1504 | 1.8171 |
| b13 | 1294.6275 | 1294.6257 | -1.3749 |
| y12 | 1182.5913 | 1182.5982 | 5.8177 |
| y13 | 1310.6499 | 1310.6527 | 2.1363 |
| $y16^{2+}$ | 820.3898 | 820.3852 | -5.5583 |
| $y18^{2+}$ | 926.4296 | 926.4343 | 5.0624 |
| $y19^{2+}$ | 969.9456 | 969.9453 | -0.3402 |

**Figure S34.** Modification of SpaA $_{G\text{-}1K\_A6+A7del}$ peptide by SpaKC-SpaF combination. (A) $His_6$-tagged SpaA$_{G\text{-}1K_A6+A7del}$ was co-expressed with SpaF and SpaKC in *E. coli*, purified and then digested by trypsin. The digestion mixture was subjected to LC-MS analysis. Digestion conditions: 20 mM Tris-HCl, pH = 8.0, 1.0 µM trypsin, 100 µM peptide product at 37 °C for 6 h. (B) MS spectrum of modified SpaA$_{G\text{-}1K_A6+A7del}$ after trypsin digestion. (C) MS/MS spectrum of **SpaA$_{G\text{-}1K_A6+A7del}$-(e)**. The *b* and *y* ions are listed in the table and marked in the spectrum.

Monoisotopic mass of SpaA$_{G\text{-}1K_A6+A7del}$ derivatives:

**SpaA$_{G\text{-}1K_A6+A7del}$-(e)**, calc. 2179.019 Da, obs. 2179.023 Da;

**(M-2$H_2$O-104)**, calc. 2121.031 Da, obs. 2121.043 Da;

**(M-104)**, calc. 2157.053 Da, obs. 2157.078 Da.

**-104**: hydrolysis of decarboxylated Cys.

**Pi**: phosphorylation

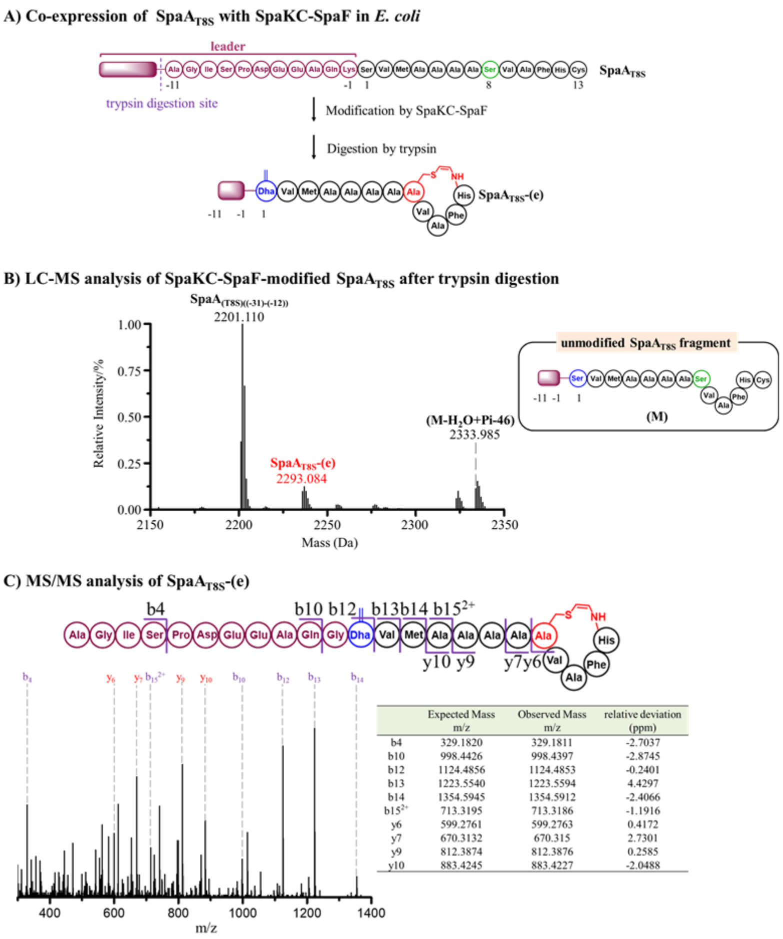


| | Expected Mass m/z | Observed Mass m/z | relative deviation (ppm) |
|---|---|---|---|
| b4 | 329.1820 | 329.1811 | -2.7037 |
| b10 | 998.4426 | 998.4397 | -2.8745 |
| b12 | 1124.4856 | 1124.4853 | -0.2401 |
| b13 | 1223.5540 | 1223.5594 | 4.4297 |
| b14 | 1354.5945 | 1354.5912 | -2.4066 |
| $b15^{2+}$ | 713.3195 | 713.3186 | -1.1916 |
| y6 | 599.2761 | 599.2763 | 0.4172 |
| y7 | 670.3132 | 670.315 | 2.7301 |
| y9 | 812.3874 | 812.3876 | 0.2585 |
| y10 | 883.4245 | 883.4227 | -2.0488 |

**Figure S35.** Modification of SpaA$_{T8S}$ peptide by SpaKC-SpaF combination. (A) His$_6$-tagged SpaA$_{T8S}$ was co-expressed with SpaF and SpaKC in *E. coli*. The product was purified, digested by trypsin and subjected to LC-MS analysis. Digestion conditions: 20 mM Tris-HCl, pH = 8.0, 1.0 μM trypsin, 100 μM peptide product at 37 °C for 6 h. (B) MS spectrum of modified SpaA$_{G\text{-}1K_V9A}$ after trypsin digestion. (C) MS/MS spectrum of **SpaA$_{T8S}$-(e)**. The *b* and *y* ions are listed in the table and marked in the spectrum.

Monoisotopic mass of SpaA$_{T8S}$ derivatives:

**SpaA$_{T8S}$-(e)**, calc. 2236.004 Da, obs. 2236.008 Da;

**(M-$H_2O$+Pi-46)**, calc. 2333.981 Da, obs. 2333.985.

**-46**: oxidative decarboxylation of Cys.

**Pi**: phosphorylation

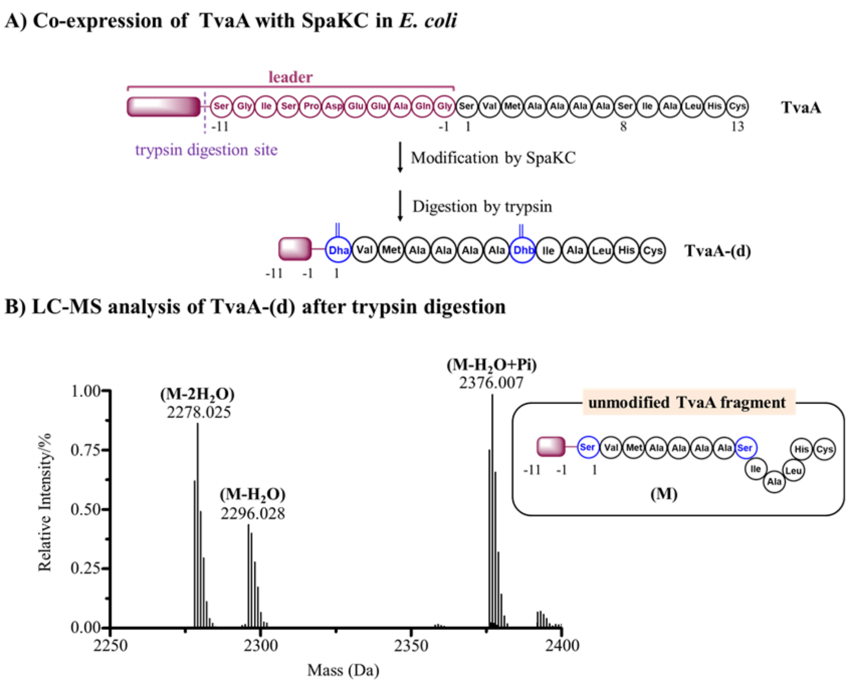


**Figure S36.** SpaKC dehydrates TvaA by up to 2-fold. (A) $His_6$-tagged TvaA was co-expressed with SpaKC in *E. coli*, purified and digested by trypsin. Digestion conditions: 20 mM Tris-HCl, pH = 8.0, 1.0 µM trypsin, 100 µM peptide product at 37 °C for 6 h. (B) MS spectrum of digested **TvaA-(d)** peptide.

Monoisotopic mass of TvaA derivatives:

**(M-$H_2O$)**: calc. 2296.046; obs. 2296.028;

**(M-2$H_2O$)**: calc. 2278.036; obs., 2278.025;

**(M-$H_2O$+Pi)**: calc. 2376.012; obs., 2376.007.

**Pi**: phosphorylation

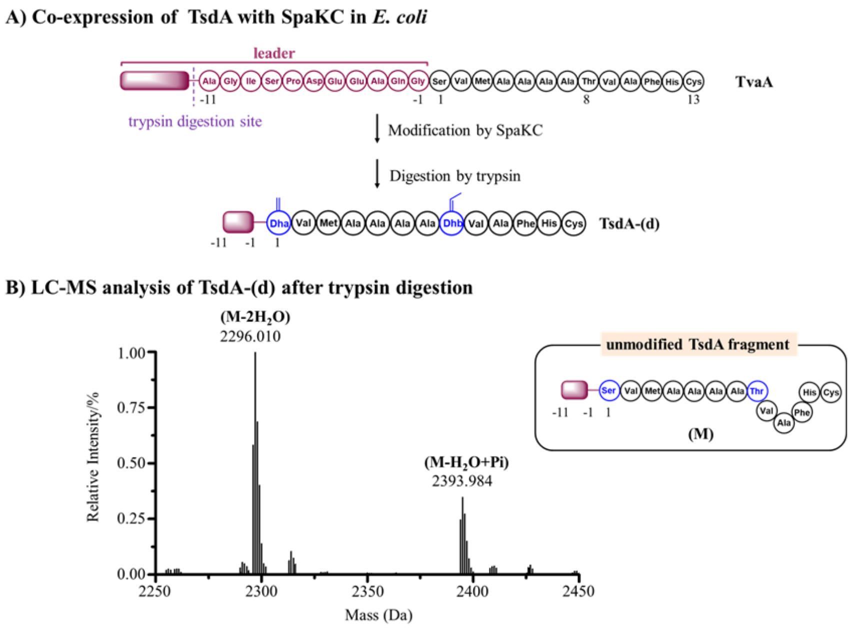


**Figure S37.** SpaKC dehydrates TsdA by up to 2-fold. (A) His$_6$-tagged TsdA was co-expressed with SpaKC in *E. coli*, purified and digested by trypsin. Digestion conditions: 20 mM Tris-HCl, pH = 8.0, 1.0 µM trypsin, 100 µM peptide product at 37 °C for 6 h.(B) MS spectrum of digested **TsdA-(d)** peptide.

Monoisotopic mass of TsdA derivative:

**(M-2$H_2$O)**:calcd. 2296.0252; obsd.,2296.0096

**(M-$H_2$O+Pi)**: calc. 2394.002 Da, obs. 2393.983 Da.

**Pi**: phosphorylation

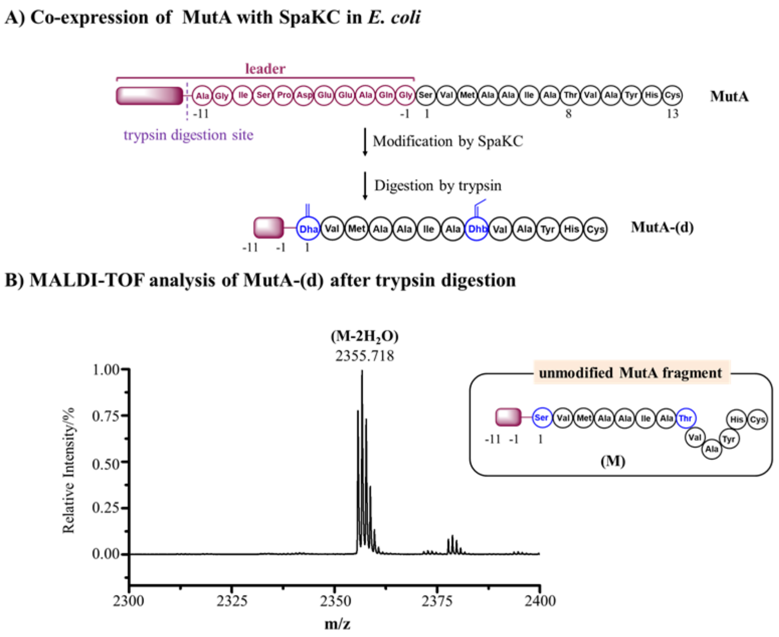


**Figure S38.** SpaKC dehydrates MutA by 2-fold. (A) $His_6$-tagged MutA was co-expressed with SpaKC in *E. coli*, purified and digested by trypsin. Digestion conditions: 20 mM Tris-HCl, pH = 8.0, 1.0 μM trypsin, 100 μM peptide product at 37 °C for 6 h. (B) MS spectrum of digested **MutA-(d)** peptide.

Monoisotopic mass of **MutA-(d)** fragment:

**(M-2$H_2$O)**: calcd. 2355.0749, obs. 2355.718.

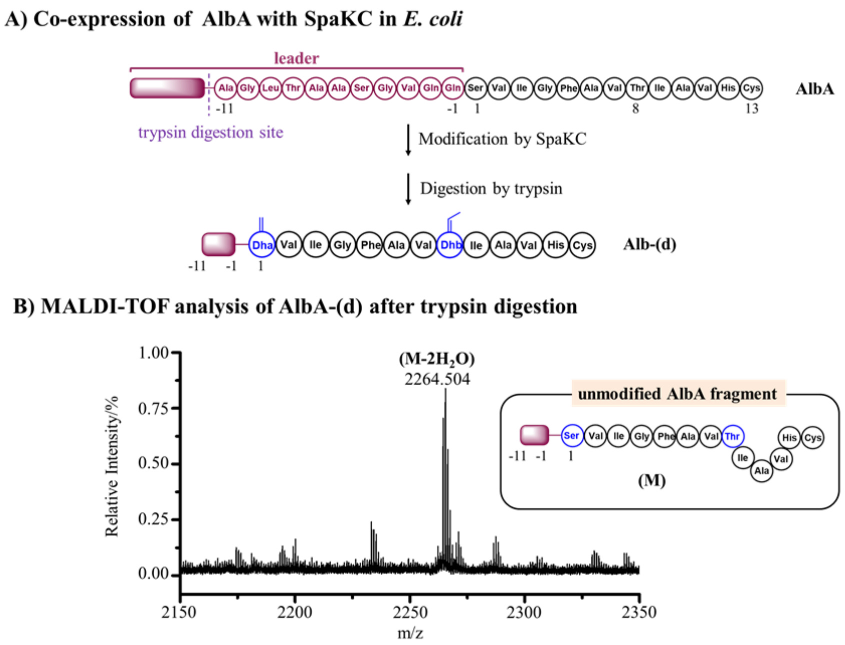


**Figure S39.** SpaKC dehydrates AlbA by 2-fold. (A) $His_6$-tagged AlbA was co-expressed with SpaKC in *E. coli*, purified and digested by trypsin. Digestion conditions: 20 mM Tris-HCl, pH = 8.0, 1.0 µM trypsin, 100 µM peptide product at 37 °C for 6 h. (B) MS spectrum of digested **AlbA-(d)** peptide.

Monoisotopic mass of AlbA derivative:

**(M-2$H_2$O)**: calcd. 2264.186, obs. 2264.504.

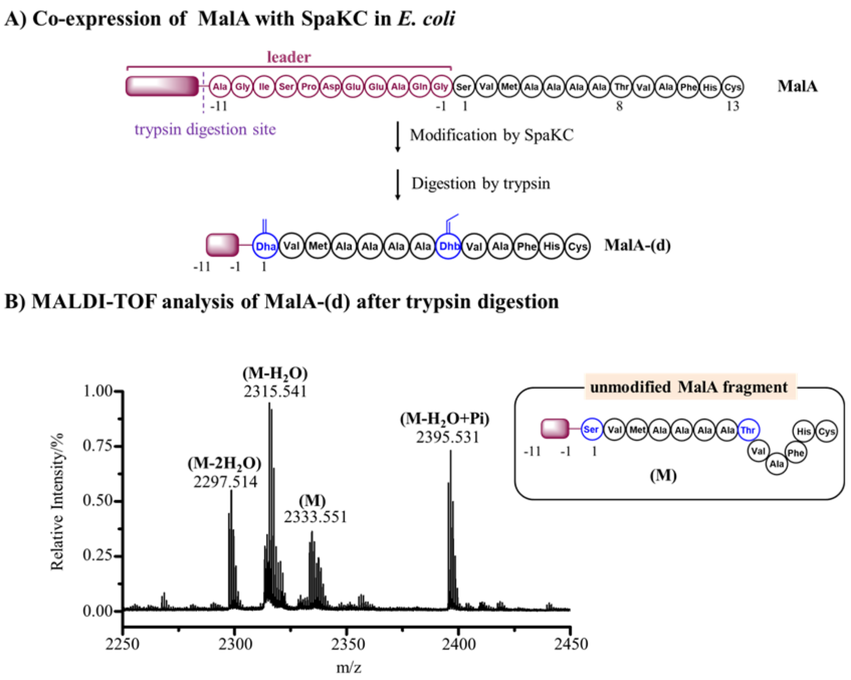


**Figure S40.** SpaKC dehydrates MalA by up to 2-fold. (A) $His_6$-tagged MalA was co-expressed with SpaKC in *E. coli*, purified and digested by trypsin. Digestion conditions: 20 mM Tris-HCl, pH = 8.0, 1.0 µM trypsin, 100 µM peptide product at 37 °C for 6 h.(B) MS spectrum of digested **MalA-(d)** peptide.

Monoisotopic mass of MalA derivative:

**(M-2$H_2$O)**: calc. 2297.0330, obs. 2297.514.

**(M-$H_2$O)**: calc. 2315.0436; obs. 2315.541.

**(M)**: calc. 2333.0542; obs. 2333.551.

**(M-$H_2$O+Pi)**: calc. 2395.0099; obs. 2395.531.

**Pi**: phosphorylation

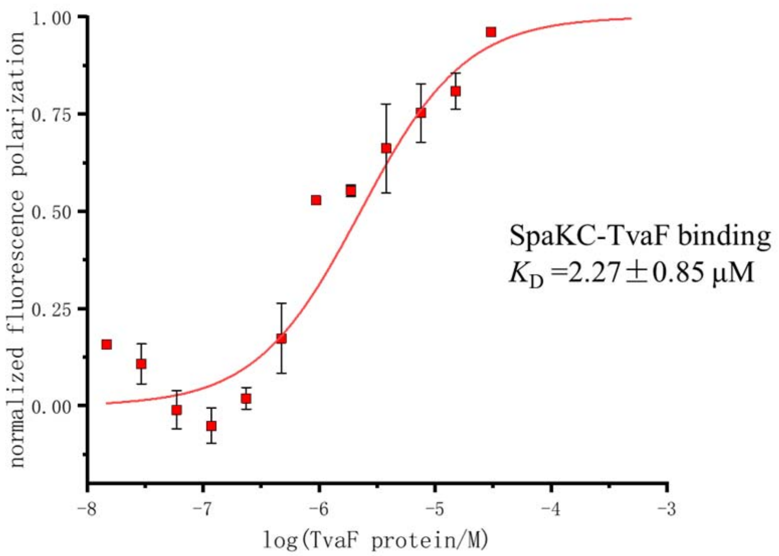


**Figure S41.** SpaKC binds to TvaF with a $K_D$ of 2.27±0.85 μM, as determined by MST assays. Error bars, s.e.m. of 3 independent measurements.

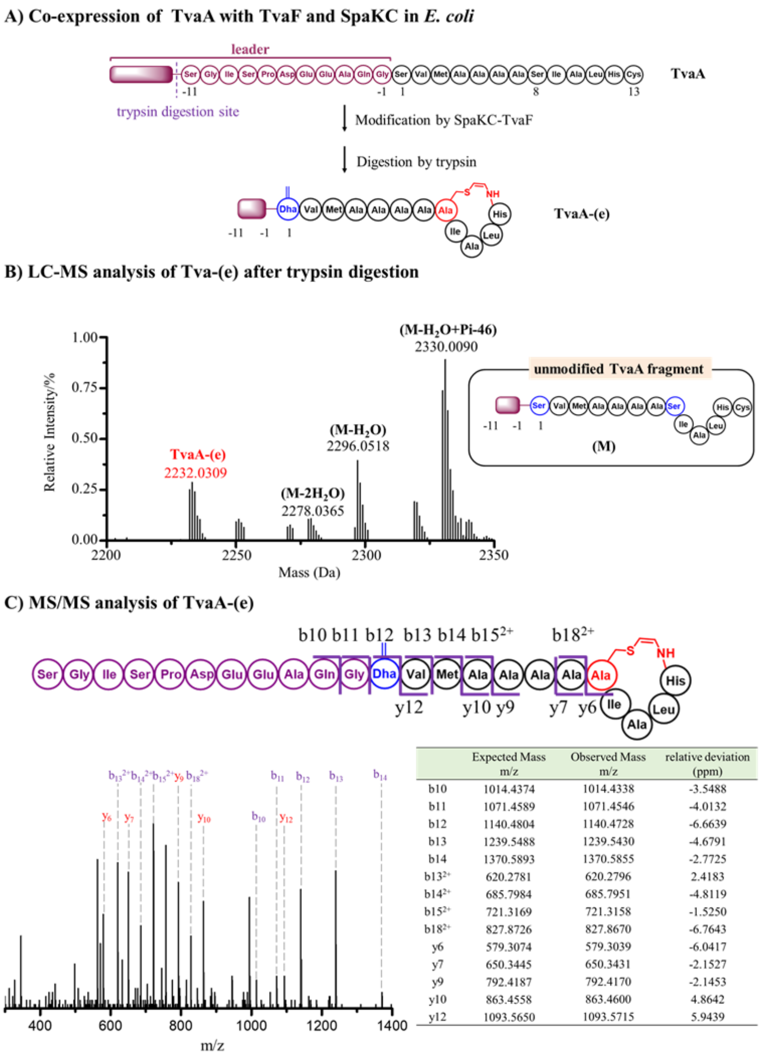


| | Expected Mass m/z | Observed Mass m/z | relative deviation (ppm) |
|---|---|---|---|
| b10 | 1014.4374 | 1014.4338 | -3.5488 |
| b11 | 1071.4589 | 1071.4546 | -4.0132 |
| b12 | 1140.4804 | 1140.4728 | -6.6639 |
| b13 | 1239.5488 | 1239.5430 | -4.6791 |
| b14 | 1370.5893 | 1370.5855 | -2.7725 |
| $b13^{2+}$ | 620.2781 | 620.2796 | 2.4183 |
| $b14^{2+}$ | 685.7984 | 685.7951 | -4.8119 |
| $b15^{2+}$ | 721.3169 | 721.3158 | -1.5250 |
| $b18^{2+}$ | 827.8726 | 827.8670 | -6.7643 |
| y6 | 579.3074 | 579.3039 | -6.0417 |
| y7 | 650.3445 | 650.3431 | -2.1527 |
| y9 | 792.4187 | 792.4170 | -2.1453 |
| y10 | 863.4558 | 863.4600 | 4.8642 |
| y12 | 1093.5650 | 1093.5715 | 5.9439 |

**Figure S42.** SpaKC-TvaF combination installs an AviCys motif in TvaA peptide. (A) $His_6$-tagged TvaA was co-expressed with TvaF and SpaKC in *E. coli*, purified and then digested by trypsin. The digestion mixture was subjected to LC-MS analysis. Digestion conditions: 20 mM Tris -HCl, pH = 8.0, 1.0 µM trypsin, 100 µM peptide product at 37 °C for 6 h. (B) LC-MS analysis of **TvaA-(e)**. (C) LC-MS/MS analysis of **TvaA-(e).** The *b* and *y* ions are listed in the table and marked in the spectrum.

Deconvoluted mass of TvaA derivatives:

**TvaA-(e)**, calcd. 2232.030 Da, obsd. 2232.030 Da;

**(M-2$H_2O$)**: calcd. 2278.035 Da, obsd. 2278.036 Da;

**(M-$H_2O$)**: calcd. 2296.046 Da, obsd. 2296.051 Da;

**(M-$H_2O$+Pi-46)**:calcd. 2330.007 Da, obsd. 2330.009 Da.

**-46**: oxidative decarboxylation of Cys

**Pi**: phosphorylation

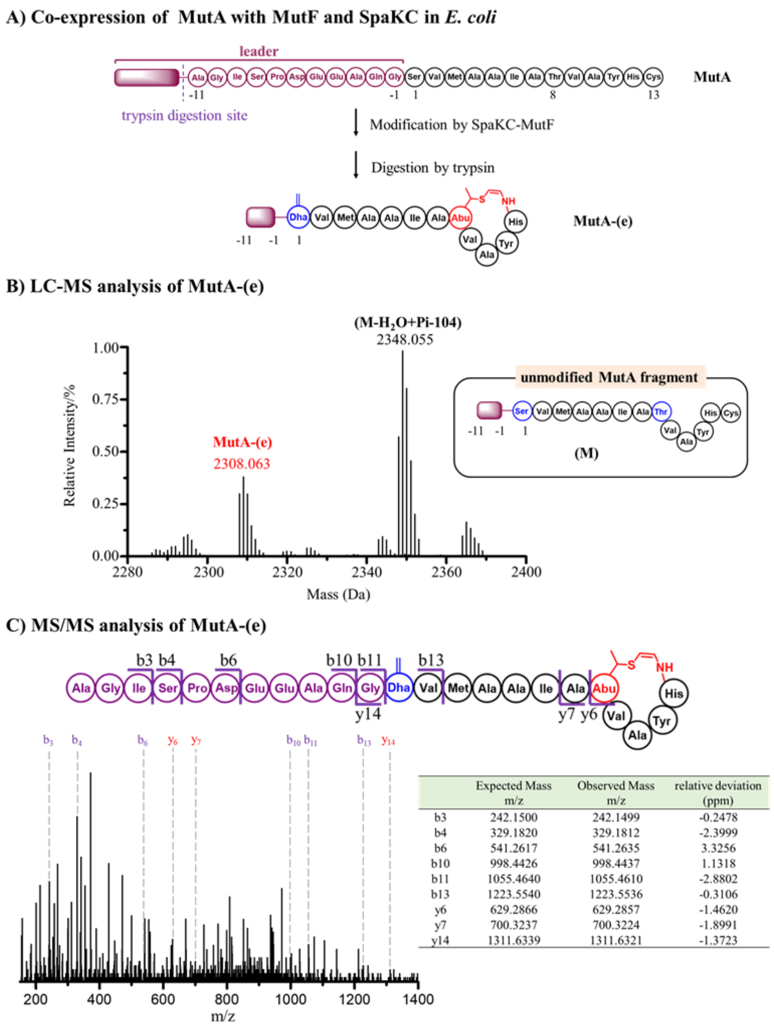


| | Expected Mass m/z | Observed Mass m/z | relative deviation (ppm) |
|---|---|---|---|
| b3 | 242.1500 | 242.1499 | -0.2478 |
| b4 | 329.1820 | 329.1812 | -2.3999 |
| b6 | 541.2617 | 541.2635 | 3.3256 |
| b10 | 998.4426 | 998.4437 | 1.1318 |
| b11 | 1055.4640 | 1055.4610 | -2.8802 |
| b13 | 1223.5540 | 1223.5536 | -0.3106 |
| y6 | 629.2866 | 629.2857 | -1.4620 |
| y7 | 700.3237 | 700.3224 | -1.8991 |
| y14 | 1311.6339 | 1311.6321 | -1.3723 |

**Figure S43.** SpaKC-MutF combination installs an AviMeCys motif in MutA peptide. (A) $His_6$-tagged MutA was co-expressed with MutF and SpaKC in *E. coli*, purified and then digested by trypsin. The digestion mixture was subjected to LC-MS analysis. Digestion conditions: 20 mM Tris-HCl, pH = 8.0, 1.0 μM trypsin, 100 μM peptide product at 37 °C for 6 h. (B) LC-MS analysis of **MutA-(e)**. (C) MS/MS analysis of **MutA-(e)**. The *b* and *y* ions are listed in the table and marked in the spectrum.

Deconvoluted mass of MutA derivatives:

**MutA-(e)**, calc. 2308.061 Da, obs. 2308.063 Da;

**(M-$H_2O$+Pi-104)**: calc. 2348.050 Da, obs. 2348.055 Da.

**-104**: hydrolysis of the decarboxylated Cys

**Pi**: phosphorylation

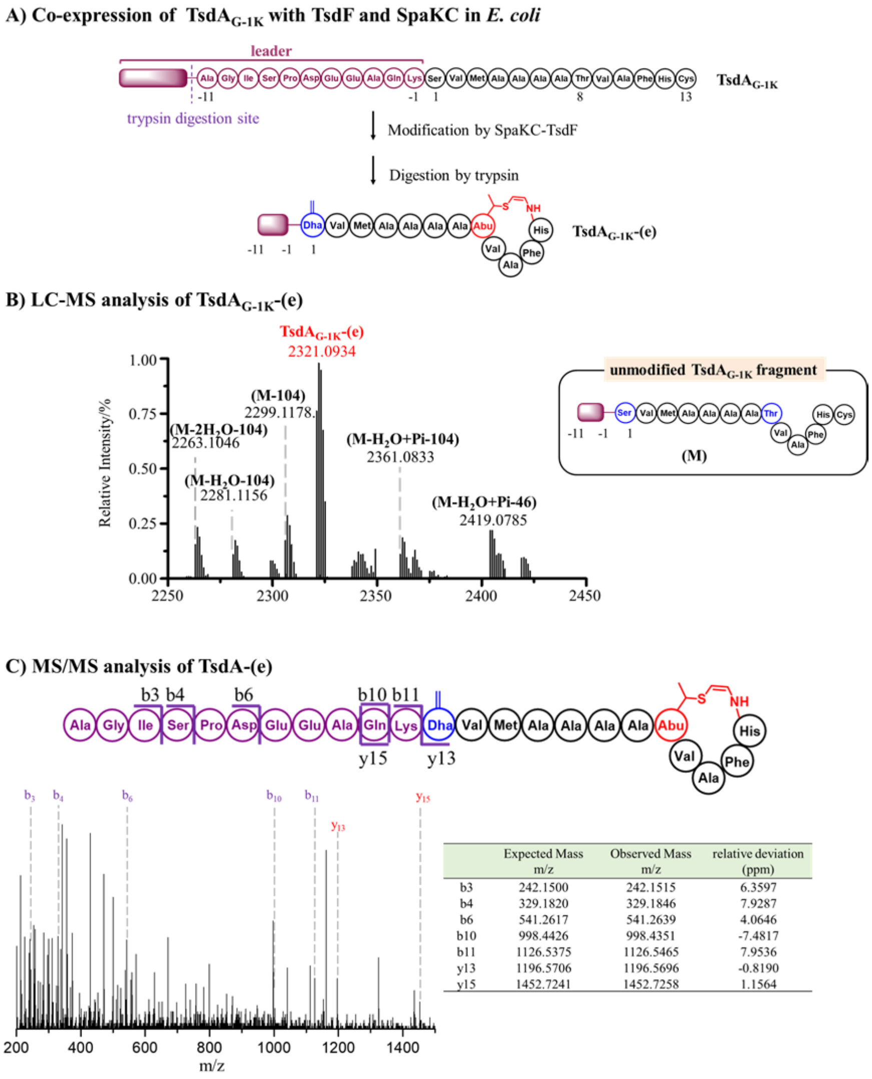


| | Expected Mass m/z | Observed Mass m/z | relative deviation (ppm) |
|---|---|---|---|
| b3 | 242.1500 | 242.1515 | 6.3597 |
| b4 | 329.1820 | 329.1846 | 7.9287 |
| b6 | 541.2617 | 541.2639 | 4.0646 |
| b10 | 998.4426 | 998.4351 | -7.4817 |
| b11 | 1126.5375 | 1126.5465 | 7.9536 |
| y13 | 1196.5706 | 1196.5696 | -0.8190 |
| y15 | 1452.7241 | 1452.7258 | 1.1564 |

**Figure S44.** SpaKC-TsdF combination installs an AviMeCys motif in TsdA peptide. (A) His$_6$-tagged TsdA was co-expressed with TsdF and SpaKC in *E. coli*, and then digested by trypsin. The digestion mixture was subjected to LC-MS analysis. Digestion conditions: 20 mM Tris-HCl, pH = 8.0, 1.0 µM trypsin, 100 µM peptide product at 37 °C for 6 h. (B) LC-MS analysis of **TsdA$_{G-1K}$-(e)**. (C) MS/MS analysis of **TsdA$_{G-1K}$-(e)**. The *b* and *y* ions are listed in the table and marked in the spectrum.

Deconvoluted mass of TsdA$_{G-1K}$ derivatives:

**(M-2H$_2$O-104)**: calc. 2263.081 Da, obs. 2263.104 Da;

**(M-H$_2$O-104)**: calc. 2281.092 Da, obs. 2281.115 Da;

**(M -104)**: calc. 2299.126 Da, obs. 2299.117 Da;

**TsdA-(e)**, calc. 2321.093 Da, obs. 2321.093 Da;

**(M-H$_2$O+Pi-104)**: calc. 2361.058 Da, obs. 2361.083 Da;

**(M-H$_2$O+Pi-46)**: calc. 2419.070 Da, obs. 2419.078 Da.

-46: oxidative decarboxylation of Cys

-62 & -104: hydrolysis of oxidative decarboxylated Cys

**Pi**: phosphorylation

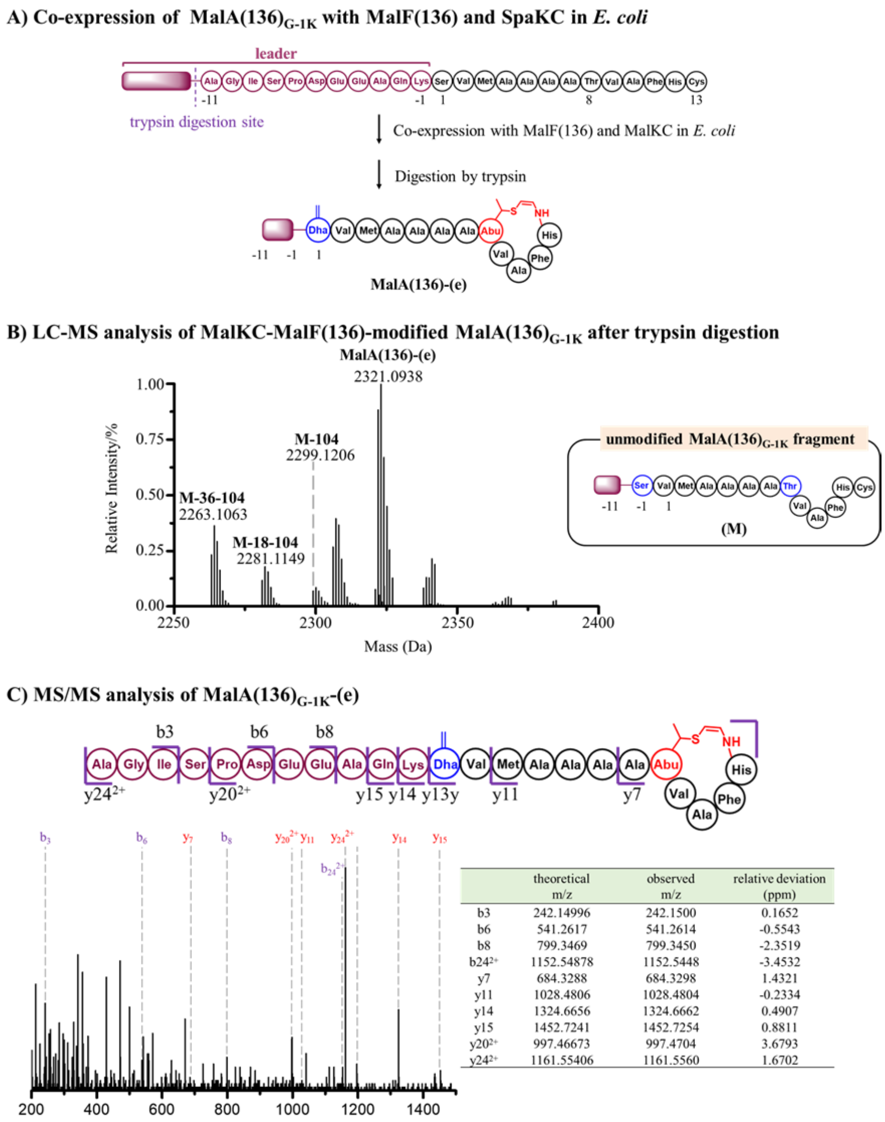


| | theoretical m/z | observed m/z | relative deviation (ppm) |
|---|---|---|---|
| b3 | 242.14996 | 242.1500 | 0.1652 |
| b6 | 541.2617 | 541.2614 | -0.5543 |
| b8 | 799.3469 | 799.3450 | -2.3519 |
| $b24^{2+}$ | 1152.54878 | 1152.5448 | -3.4532 |
| y7 | 684.3288 | 684.3298 | 1.4321 |
| y11 | 1028.4806 | 1028.4804 | -0.2334 |
| y14 | 1324.6656 | 1324.6662 | 0.4907 |
| y15 | 1452.7241 | 1452.7254 | 0.8811 |
| $y20^{2+}$ | 997.46673 | 997.4704 | 3.6793 |
| $y24^{2+}$ | 1161.55406 | 1161.5560 | 1.6702 |

**Figure S45.** SpaKC-MalF(136) combination installs an AviMeCys motif in $MalA_{G\text{-}1K}$ peptide. (A) $His_6$-tagged $MalA(136)_{G\text{-}1K}$ was co-expressed with MalF(136) and SpaKC in *E. coli*, purified and then digested by trypsin. Digestion conditions: 20 mM Tris-HCl, pH = 8.0, 1.0 µM trypsin, 100 µM peptide product at 37 °C for 6 h. (B) LC-MS analysis of **MalA(136)-(e)**. (C) MS/MS analysis of **MalA(136)-(e)**. The *b* and *y* ions are listed in the table and marked in the spectrum.

Deconvoluted mass of $MalA_{G\text{-}1K}$ derivatives:

**(M-2$H_2O$-104)**: calc. 2263.081 Da, obs. 2263.106 Da;

**(M-$H_2O$-104)**: calc. 2281.092 Da, obs. 2281.115 Da;

**(M-104)**: calc. 2299.126 Da, obs. 2299.121 Da;

**MalA(136)-(e)**, calc. 2321.093 Da, obs. 2321.094 Da.

**-104**: hydrolysis of the oxidative decarboxylated Cys

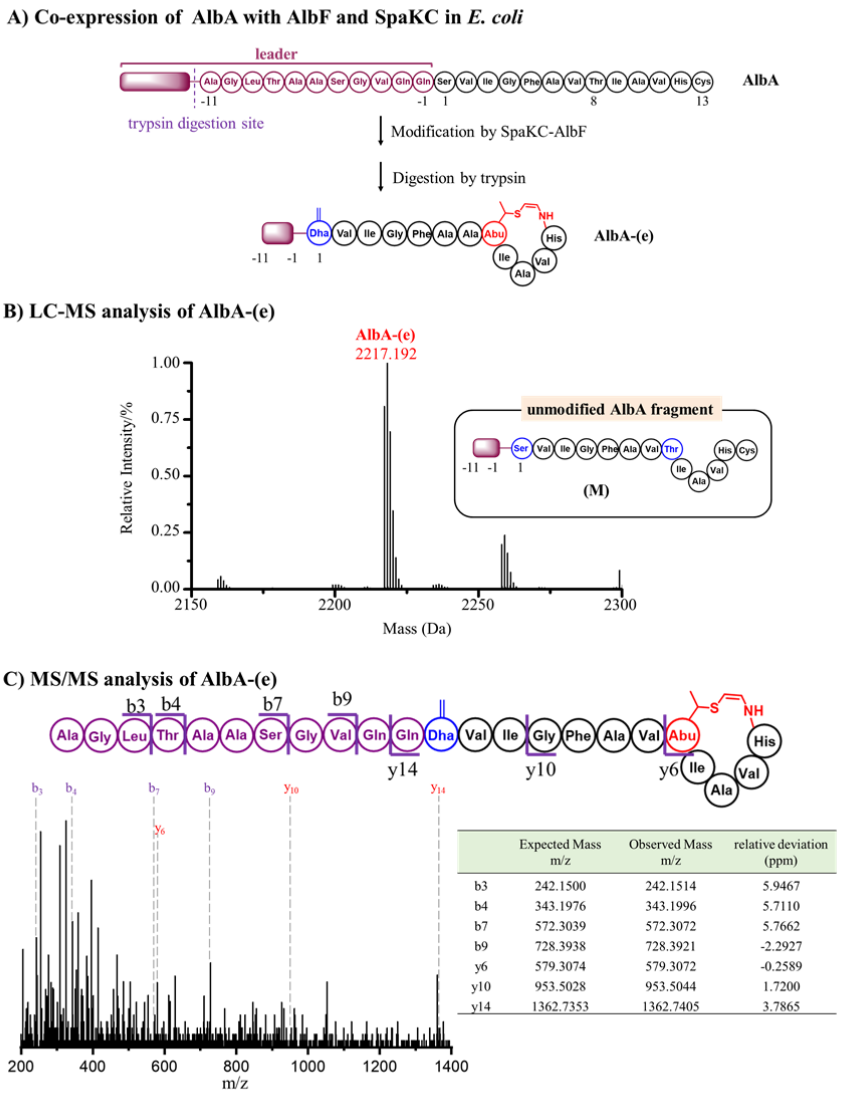


| | Expected Mass m/z | Observed Mass m/z | relative deviation (ppm) |
|---|---|---|---|
| b3 | 242.1500 | 242.1514 | 5.9467 |
| b4 | 343.1976 | 343.1996 | 5.7110 |
| b7 | 572.3039 | 572.3072 | 5.7662 |
| b9 | 728.3938 | 728.3921 | -2.2927 |
| y6 | 579.3074 | 579.3072 | -0.2589 |
| y10 | 953.5028 | 953.5044 | 1.7200 |
| y14 | 1362.7353 | 1362.7405 | 3.7865 |

**Figure S46.** SpaKC-AblF combination installs an AviMeCys motif in AblA peptide. (A) $His_6$-tagged AlbA was co-expressed with AlbF and SpaKC in *E. coli*, purified and then digested by trypsin. (B) LC-MS analysis of **AlbA-(e)**.

Digestion conditions: 20 mM Tris-HCl, pH = 8.0, 1.0 µM trypsin, 100 µM peptide product at 37 °C for 6 h. (C) LC-MS/MS analysis of **AlbA-(e)**. The *b* and *y* ions are listed in the table and marked in the spectrum.

Deconvoluted mass of AlbA derivatives:

**AlbA-(e)**, calc. 2217.172 Da, obs. 2217.192 Da.

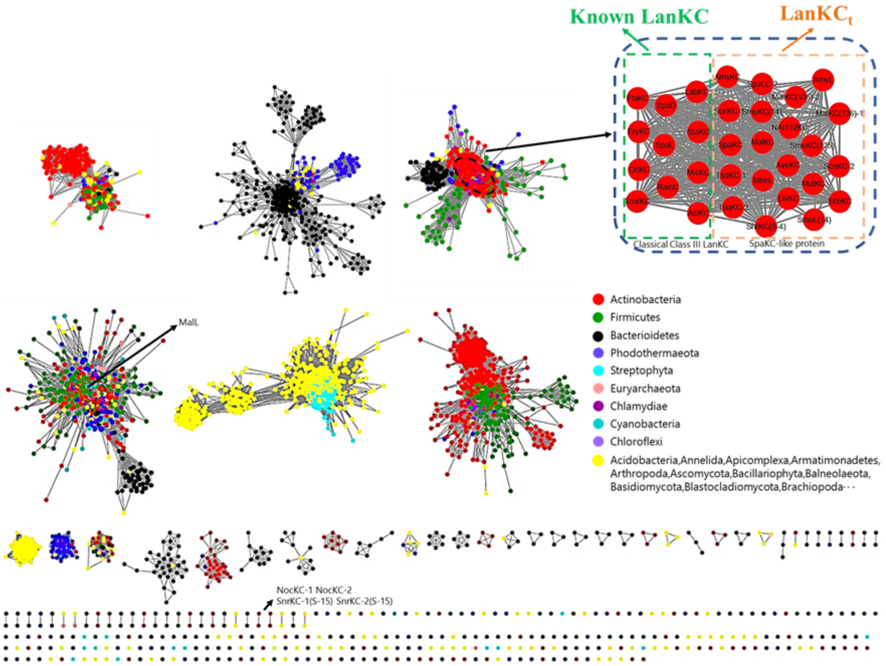


**Figure S47.** Phylogenetic analysis of class III lanthipeptide synthetase LanKC and LanKC-like enzymes was completed using a sequence similarity network (SSN). Sequences for the network were accessed by a 5920 sequence return, 10 classical Class III LanKC and 27 SpaKC homology from BLAST with SpaKC are mixed with 5883 protein from UniRef90 clusters (UniProt IDs clustered at ≥90% sequence identity) of PF05147 Pfam. The SSN was built using the online Enzyme Function Initiative-Enzyme Similarity Tool (EFIEST) using the 5920 sequences that had both an E-value of ≤e−5 The 100% identity representative node network with 1030190 edges, which contains 1687 nodes, was visualized in Cytoscape208 with an alignment score threshold of 40 (~20% sequence identity).

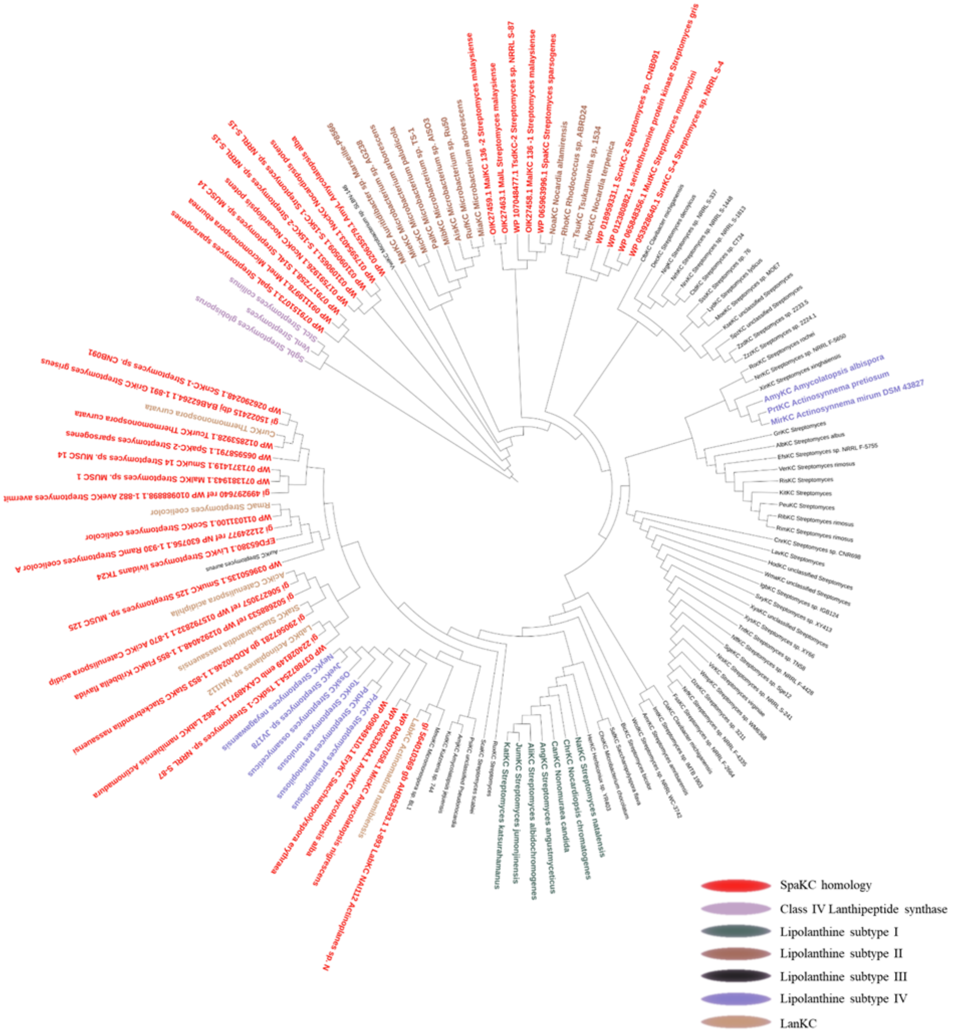


**Figure S48.** Maximum-likehood phylogeny of LanKC$_t$ enzymes discovered from thioamitide-producing strains and selected class III lanthipeptide synthetases, including LanKC enzymes from lipolanthine biosynthesis. Class IV LanL enzymes are shown as the outgroup.

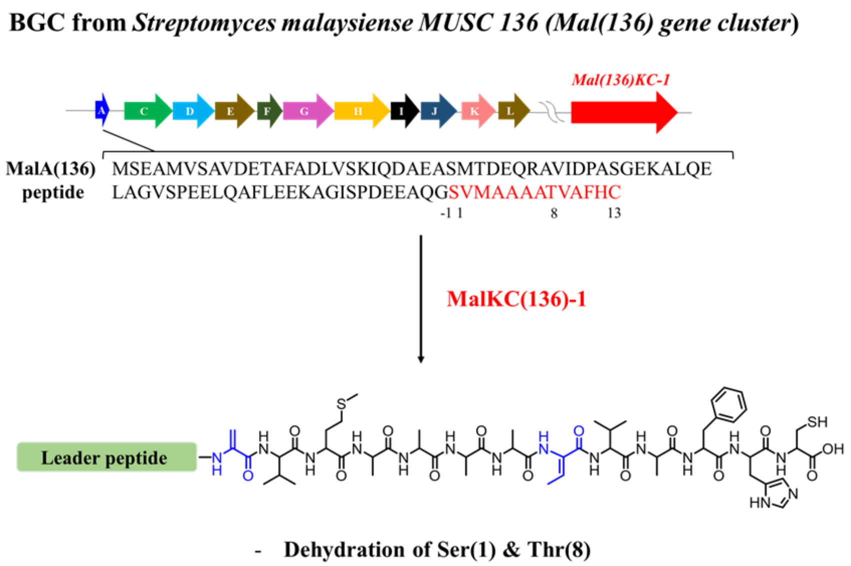


**Figure S49.** Putative thioamitide biosynthetic gene cluster from *S. malaysiense* MUSC 136 and class III lanthipeptide synthetase homolog MalKC(136)-1 encoded outside the *malA* gene cluster. Co-expression of MalKC(136)-1 and MalA(136) peptide in E. coli confirmed its function as a dehydratase.

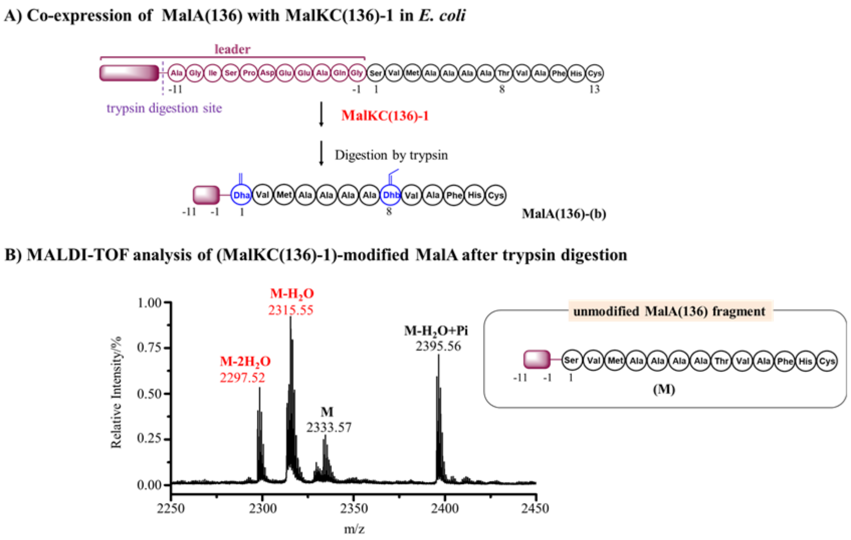


**Figure S50.** Modification of MalA(136) by MalKC(136)-1 from *S. malaysiense* MUSC 136. (A) $His_6$-tagged MalA(136) peptide was co-expressed with MalKC(136)-1 in *E. coli*, purified and then digested by trypsin. (B) MALDI-ToF analysis of the modification products.

Digestion conditions: 20 mM Tris-HCl, pH = 8.0, 1.0 µM trypsin, 100 µM peptide product at 37 °C for 6 h.

**(M-2$H_2$O)** , calc. 2297.03, obs. 2297.52.

**(M-$H_2$O)**, calc. 2315.04; obs. 2315.55.

**(M)**, calc. 2333.05; obs. 2333.57.

**(M-$H_2$O+80)**, calc. 2395.01; obs. 2395.56.

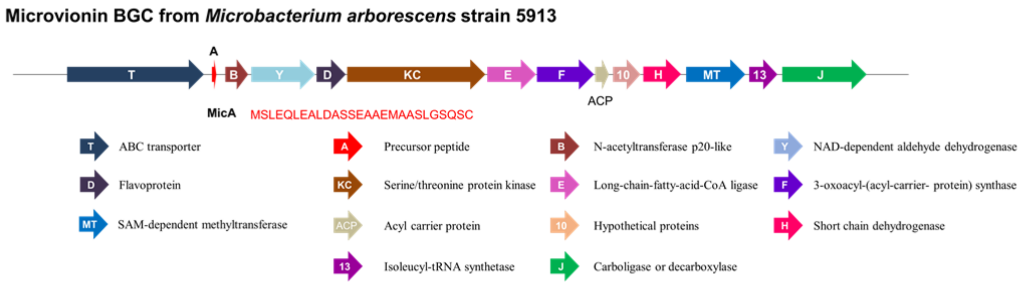


**Figure S51.** The biosynthetic gene cluster of microvionin.
